# Single-cell profiling identifies STAT3/NF-κB regulatory hubs and cytokine crosstalk in the tumor microenvironment

**DOI:** 10.64898/2026.08.14.744802

**Authors:** Mary Oluwabusolami Odubote, Chiemeka Elochi Emeribe

**Author notes:** **Corresponding author:** Chiemeka Elochi Emeribe Department of Cellular and Molecular Biology, Purdue University West Lafayette, IN 47907, USA.

## Abstract

Tumor progression is driven by dynamic interactions between malignant cells and the tumor microenvironment (TME), yet the regulatory mechanisms governing cellular heterogeneity and intercellular communication remain incompletely characterized. Here, we performed integrative single-cell RNA sequencing (scRNA-seq) analysis of publicly available datasets from non-small cell lung cancer and breast cancer to systematically map transcriptional heterogeneity and regulatory networks within the TME.

Using a unified computational pipeline with Seurat v5, SCENIC, and ligand–receptor modeling, we resolved major cellular populations, including malignant epithelial cells, immune subsets, cancer-associated fibroblasts, and endothelial cells, and their transcriptional states. Malignant cells displayed pronounced intratumoral heterogeneity, occupying a continuum of proliferative, metabolic, and invasive phenotypes linked by pseudotime trajectories.

Gene regulatory network inference identified STAT3, NF-κB, MYC, and HIF-1α as central hubs coordinating tumor-associated programs. Notably, we uncovered a cytokine-mediated immunoregulatory axis between malignant cells and tumor-associated macrophages, driven by IL6– IL6R and CCL2–CCR2 signaling. Cell–cell communication analysis further revealed coordinated networks supporting immune suppression, inflammation, and angiogenesis.

These findings provide a systems-level framework of TME organization and highlight key transcriptional circuits and signaling pathways as promising targets for disrupting tumor– microenvironment crosstalk in precision oncology.

## 1. INTRODUCTION

### 1.1 Tumor Microenvironment Complexity and Cancer Progression

Cancer is evolving into a very complex disease not solely driven by cancer cells but the surrounding cellular and molecular environment termed the tumor microenvironment (TME) [1,2]. The TME is in reality a heterogeneous interaction of cancer cells and immune cells, cancer associated fibroblasts (CAFs) and endothelial cells, T cells, and signals on tumour development and therapy. Tumor cells actively communicate with surrounding non-malignant cells, including regulatory T cells and myeloid-derived suppressor cells, through cytokines and signaling molecules that promote immune evasion [3,4]. By synthesizing cytokines, chemokines as well as immune checkpoint molecules, tumors can escape immune surveillance and increase their chances of survival [3,4].

Cancer-associated fibroblasts, in contrast, also have significant influence on the remodeling of the stromal matrix and are known to have signaling molecules that trigger tumor cell invasion and metastasis [2,5]. Angiogenesis in the tumour is also necessary within TME. Endothelial cells, which are produced by pro-angiogenic factors like vascular endothelial growth factor (VEGF), are capable of forming an abnormal vascular network with an oxygen and nutrient supply for tumour growth [1,6]. Taken together, immune suppression, stromal remodeling and angiogenic signaling demonstrate that the tumor microenvironment has a critical role to play in cancer progression and resistance against treatment [2].

### 1.2 Intratumoral Heterogeneity in Cancer Biology

Intratumoral heterogeneity is a hallmark of cancer and represents one of the most significant obstacles to effective therapy [7,8]. Tumors are composed of genetically and transcriptionally diverse cell populations that evolve over time through mutation accumulation, clonal expansion, and selective pressures imposed by the tumor microenvironment [7,8]. These evolutionary processes generate multiple malignant subclones with distinct molecular and functional characteristics, contributing to tumor progression and therapeutic resistance [9].

Genetic heterogeneity arises primarily from genomic instability and the continuous acquisition of somatic mutations affecting oncogenes, tumor suppressor genes, and signaling pathways involved in cell proliferation and survival [9,8]. As tumor cells proliferate, selective pressures within the microenvironment favor the expansion of subclones that possess survival advantages, resulting in complex clonal architectures within tumors [7]. In addition to genetic diversity, transcriptional heterogeneity further contributes to functional variability among tumor cells by generating distinct cellular states, including proliferative, invasive, and therapy-resistant phenotypes [10,11]

The presence of heterogeneous subpopulations has profound implications for cancer therapy. Treatments targeting specific molecular pathways may eliminate sensitive tumor cells while leaving resistant subclones unaffected, ultimately leading to disease recurrence or metastasis [7,8]. Therefore, understanding the molecular basis of intratumoral heterogeneity is essential for the development of precision oncology strategies and personalized treatment approaches.

### 1.3 Advantages of Single-Cell Transcriptomics

Traditional bulk RNA sequencing has been widely used to investigate gene expression patterns in tumors; however, this approach measures the average transcriptomic signal across large populations of cells, thereby masking cellular heterogeneity [12,13]. As a result, rare cell populations and subtle transcriptional differences among individual cells are often obscured in bulk analyses [12]. This limitation has hindered the ability to fully understand tumor complexity and the interactions occurring within the tumor microenvironment.

Single-cell transcriptomics has emerged as a transformative technology that enables gene expression profiling at the resolution of individual cells [14,15]. By analyzing thousands to millions of cells simultaneously, single-cell RNA sequencing (scRNA-seq) allows researchers to identify distinct cellular populations, characterize transcriptional states, and uncover rare or transient cell types involved in tumor progression [13,15]. This high-resolution approach has significantly enhanced our understanding of tumor heterogeneity and the functional roles of immune and stromal cells within the TME [11].

Several technological platforms have contributed to the advancement of single-cell transcriptomic studies. Droplet-based scRNA-seq technologies enable high-throughput analysis of thousands of cells through microdroplet encapsulation and barcoding strategies [16,17]. Microfluidic capture systems provide controlled cell isolation and efficient library preparation for transcriptomic sequencing [12]. Plate-based sequencing methods offer higher transcript coverage and are particularly suitable for investigating rare cell populations [14]. Collectively, these technologies provide powerful tools for dissecting tumor heterogeneity and exploring the molecular mechanisms underlying cancer progression.

### 1.4 Research Gap and Objectives

Despite advances in single-cell transcriptomics, significant challenges remain in integrating heterogeneous datasets to systematically resolve regulatory networks and intercellular communication within the tumor microenvironment [2,15]. Most existing studies focus on individual tumor types or limited datasets, which restrict the identification of conserved regulatory mechanisms driving tumor progression across diverse contexts [12,13]. Furthermore, the coordination between transcriptional regulatory networks and cell–cell communication pathways remains incompletely characterized [3].

To address these limitations, this study performs an integrative single-cell transcriptomic analysis across multiple publicly available datasets to characterize tumor microenvironment heterogeneity, reconstruct gene regulatory networks, and identify intercellular communication pathways driving tumor progression. By combining clustering, trajectory inference, regulatory network modeling, and ligand–receptor analysis within a unified computational framework, this work provides a systems-level perspective of tumor–microenvironment interactions.

Recent advances in computational biology and systems genomics further support the integration of single-cell transcriptomic data with network-based modeling approaches to identify transcriptional regulators and signaling pathways governing tumor–ecosystem interactions [18,19]. These approaches enable the reconstruction of gene regulatory networks and ligand–receptor-mediated communication across cellular populations [15,18], providing a foundation for interpreting complex tumor ecosystem dynamics.

Therefore, this study employs single-cell transcriptomic profiling to investigate the cellular and molecular architecture of the tumor microenvironment during cancer progression. Specifically, the objectives are to:

i. identify distinct cellular populations and characterize transcriptional heterogeneity within tumors,
ii. reconstruct gene regulatory networks that govern tumor progression, and
iii. map intercellular communication pathways mediating interactions between malignant and microenvironmental cells.

Through these analyses, the study aims to provide deeper insights into the regulatory mechanisms underlying tumor evolution and therapeutic resistance.

## 2. MATERIALS AND METHODS

### 2.1 Data Sources and Study Design

This study analyzed publicly available single-cell RNA sequencing (scRNA-seq) datasets to investigate transcriptional heterogeneity and regulatory interactions within the tumor microenvironment (TME). Transcriptomic datasets were obtained from major public repositories, including the Gene Expression Omnibus (GEO), the European Nucleotide Archive (ENA), and the Single Cell Portal, which provide high-throughput sequencing data and standardized metadata for downstream analysis [15,20]. Only datasets generated using droplet-based sequencing platforms, such as 10x Genomics Chromium, were included to ensure compatibility across samples and minimize technical variation [16,17].

To ensure data relevance and biological consistency, this study incorporated well-characterized scRNA-seq datasets representing tumor microenvironment composition across multiple cancer contexts. These included datasets examining non-small cell lung cancer following immunotherapy [21], breast tissue spanning normal to tumorigenic states [22,23], and tumor–stromal interaction dynamics in breast cancer [24]. In addition, recent multi-omics and machine learning-integrated studies were used to support analytical interpretation of immune cell heterogeneity and regulatory mechanisms [25]. These datasets collectively capture diverse tumor microenvironment conditions, including immune remodeling, stromal interaction, and tumor progression states.

Raw count matrices and accompanying metadata were downloaded using the GEOquery package (v2.66.0) in R (v4.3.1). Data processing and downstream analyses were performed using the Seurat single-cell analysis framework (v5), which introduces the Assay5 layer-based data structure and improved scalability for integrated single-cell analysis [26]. The computational workflow was executed on a Linux-based high-performance computing environment equipped with 64 GB RAM and 16 CPU cores, enabling efficient processing of large-scale datasets.

Each dataset consisted of gene expression matrices representing transcript counts for individual cells, typically ranging from 3,000 to over 50,000 cells per sample depending on study design. Samples included tumor tissues obtained from both primary and treatment-associated contexts, enabling comparative evaluation of tumor microenvironment composition across disease stages. Metadata fields extracted from the repositories included patient identifiers, tumor type, sequencing platform, tissue origin, clinical stage, and treatment information. Within the Seurat v5 framework, expression data were organized into assay layers, allowing raw counts, normalized values, and scaled data to be processed within a unified object structure [27].

Datasets were included if they satisfied the following criteria: (1) availability of raw single-cell count matrices, (2) tumor-derived tissue samples containing both malignant and microenvironmental cells, (3) sequencing performed using droplet-based single-cell technologies, and (4) presence of accompanying clinical or biological metadata. Datasets lacking complete metadata or generated using incompatible platforms were excluded.

Prior to downstream analysis, datasets were integrated using the Seurat v5 integration workflow, which employs anchor-based methods to correct batch effects and align shared biological structure across datasets [28]. Integration anchors were identified using the first 30 principal components, and datasets were merged using standard Seurat functions adapted for the v5 framework.

After preprocessing and quality control filtering (described in Section 2.2), a high-quality subset of cells was retained for downstream analyses. The final integrated dataset provided a unified gene expression matrix suitable for investigating tumor micro-environment heterogeneity, regulatory networks, and intercellular communication patterns while ensuring reproducibility through standardized computational pipelines and publicly accessible data sources.

### 2.2 Single-Cell Data Preprocessing and Quality Control

Rigorous preprocessing and quality control procedures were implemented prior to downstream analysis to ensure the reliability and biological validity of the single-cell transcriptomic datasets. Raw scRNA-seq data frequently contain technical artifacts arising from library preparation, amplification bias, and droplet encapsulation inefficiencies, which may introduce noise and distort biological interpretation if not properly addressed [12]. Consequently, systematic filtering and normalization steps were applied to retain high-quality cells while minimizing technical variability [29]. These procedures are essential for improving clustering accuracy and ensuring robust downstream inference of cellular states [28].

All preprocessing procedures were conducted using Seurat (v5) within R (v4.3.1), which introduces an enhanced layer-based Assay5 data structure for efficient handling of raw, normalized, and scaled expression data within a unified framework [30]. Raw count matrices were first converted into Seurat objects and subjected to initial inspection to evaluate sequencing depth, gene detection distributions, and overall dataset integrity. Cells were filtered using established quality control thresholds to ensure biological relevance. Cells expressing fewer than 200 genes were removed to eliminate low-quality or damaged cells with insufficient transcript capture. Conversely, cells expressing more than 6,000 genes were flagged as potential multiplets or doublets, as abnormally high gene counts often indicate the presence of multiple cells captured within a single droplet [29].

To further improve data quality, doublet detection was performed using the DoubletFinder algorithm (v2.0.3), which remains a widely adopted method for identifying artificial cell multiplets in droplet-based scRNA-seq data [31]. Parameter settings included pN = 0.25, with pK optimized through parameter sweeping, and an expected doublet rate of 7.5%, consistent with standard droplet-based sequencing estimates. Cells identified as doublets were removed prior to downstream analysis to prevent distortion of cluster structure and gene expression patterns.

Another critical quality control metric involves the proportion of mitochondrial gene expression within each cell. Elevated mitochondrial transcript levels are commonly associated with cellular stress, apoptosis, or compromised membrane integrity, which can bias downstream analyses if retained [29]. Therefore, cells exhibiting mitochondrial gene expression exceeding 10% of total transcripts were excluded. This threshold is widely applied in single-cell workflows to ensure that retained cells represent viable and transcriptionally active populations [28].

Following filtering, normalization and scaling were performed within the Seurat v5 framework to correct for sequencing depth differences and stabilize variance across genes. The updated workflow leverages improved data layer management, allowing consistent handling of raw counts, normalized data, and scaled matrices during preprocessing [27]. These preprocessing steps collectively ensure that downstream analyses, including clustering, differential expression, and trajectory inference, are driven by biologically meaningful variation rather than technical noise.

Following quality filtering, gene expression values were normalized to account for differences in sequencing depth and library size across cells. Normalization was performed using the LogNormalize method implemented in Seurat, which scales gene counts by the total expression within each cell and multiplies the resulting value by a scale factor of 10,000, followed by logarithmic transformation. The normalization procedure can be expressed as:

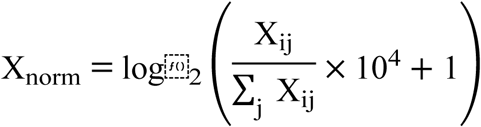

where X_ij_represents the raw expression level of gene jin cell i, and the denominator corresponds to the total transcript count within cell i. This normalization approach stabilizes variance and allows direct comparison of gene expression across cells [28].

Highly variable genes were identified using the FindVariableFeatures function in Seurat with the vst selection method, selecting the top 2,000 highly variable genes for downstream analysis. The dataset was then scaled using the ScaleData function, which centers gene expression values and removes unwanted sources of technical variation. During scaling, total UMI counts and mitochondrial gene percentage were included as covariates to regress out technical effects.

These preprocessing steps produced a high-quality normalized expression matrix suitable for dimensionality reduction, clustering, pseudotime analysis, and regulatory network inference in subsequent analytical stages [29].

### 2.3 Dimensionality Reduction and Cell Clustering

Following preprocessing and normalization, dimensionality reduction techniques were applied to identify transcriptional patterns and reveal distinct cellular populations within the tumor microenvironment. Single-cell transcriptomic datasets typically contain thousands of genes measured across thousands of cells, resulting in high-dimensional data structures that are computationally challenging to analyze directly [15,29]. Dimensionality reduction methods transform these datasets into lower-dimensional representations while preserving biologically meaningful variance and reducing technical noise [32].

Dimensionality reduction was performed using the Seurat package (v5) in R (v4.3.1). Principal Component Analysis (PCA) was applied to the scaled gene expression matrix using the RunPCA() function in Seurat. PCA was computed using the top 2,000 highly variable genes identified during preprocessing. The number of principal components retained for downstream analysis was determined using ElbowPlot inspection and JackStraw significance testing, resulting in the selection of the first 30 principal components (PCs), which captured the majority of biologically relevant variance within the dataset [28].

To visualize transcriptional heterogeneity across cellular populations, Uniform Manifold Approximation and Projection (UMAP) was applied to the PCA-reduced dataset using the RunUMAP() function in Seurat with the following parameters:

● dims = 1:30
● n.neighbors = 30
● min.dist = 0.3
● metric = “cosine”

UMAP preserves both local and global structure in high-dimensional data and has become a standard approach for visualizing single-cell transcriptomic datasets [29,33]. The resulting two-dimensional embedding enabled visualization of transcriptionally distinct cell clusters within the tumor microenvironment.

Cell clustering was performed using a graph-based community detection approach implemented in Seurat. First, a k-nearest neighbor (KNN) graph was constructed using the FindNeighbors() function with k = 20 based on the selected PCA dimensions. Cells were represented as nodes in the graph, and transcriptional similarity between cells was quantified using Euclidean distance:

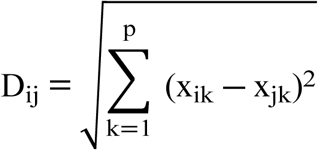

where D_ij_represents the distance between cells iand j, and x_ik_and x_jk_represent gene expression values for gene kin each respective cell.

Community detection was then performed using the Louvain clustering algorithm via the FindClusters() function with a resolution parameter of 0.6, which determines cluster granularity [34]. This approach identifies clusters by maximizing modularity within the cell–cell similarity graph, enabling detection of transcriptionally coherent cellular populations.

Cluster identities were assigned based on expression patterns of canonical marker genes associated with major tumor microenvironment cell types. Marker genes were identified using the FindAllMarkers() function with the Wilcoxon rank-sum test and default Seurat parameters. Annotated clusters included malignant epithelial cells, immune cell populations, stromal fibroblasts, and endothelial cells, consistent with previously reported tumor microenvironment cellular architectures [11].

The dimensionality reduction and clustering results enabled systematic characterization of transcriptionally distinct cellular populations and provided the foundation for subsequent differential expression, trajectory inference, regulatory network analysis, and cell–cell communication modeling.

### 2.4 Differential Gene Expression Analysis

Differential gene expression (DGE) analysis was conducted to identify genes exhibiting statistically significant transcriptional differences between tumor-associated cellular populations and reference cell groups. Detecting differentially expressed genes enables identification of transcriptional programs associated with tumor progression, immune modulation, and stromal remodeling within the tumor microenvironment [11,28]

Differential expression analysis was performed using the FindAllMarkers() function implemented in Seurat (v5) within R (v4.3.1). The analysis compared gene expression profiles between identified cell clusters generated during the clustering step. Statistical significance was evaluated using the Wilcoxon rank-sum test, which is commonly used in single-cell transcriptomic studies due to its robustness in analyzing sparse and non-normally distributed gene expression data [29,35].

The following parameters were applied during differential expression testing:

● test.use = “wilcox”
● min.pct = 0.25 (genes expressed in at least 25% of cells in either group)
● logfc.threshold = 0.25 (minimum log fold change threshold used during initial testing)
● only.pos = FALSE (both upregulated and downregulated genes considered)

For each gene, expression values were compared between cell populations to identify transcriptional signatures associated with specific cellular states. The magnitude of expression differences was quantified using log-transformed fold change calculated as:

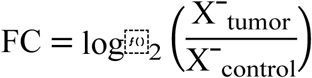

where Xˉ_tumor_represents the average expression level of a gene in tumor cells and Xˉ_control_represents the average expression level in the comparison cell population. Log transformation stabilizes variance and facilitates interpretation of upregulated or downregulated transcriptional patterns [28].

Because thousands of genes are tested simultaneously in single-cell transcriptomic datasets, correction for multiple hypothesis testing was applied using the Benjamini–Hochberg False Discovery Rate (FDR) method to control for false-positive discoveries [35,36]. Genes were considered significantly differentially expressed if they satisfied the following criteria:

● ^∣^log⁡_2_FC∣ > 1
● adjusted p-value (FDR) < 0.05

The resulting set of significant genes was used to identify cluster-specific marker genes that characterize tumor, immune, and stromal cell populations. These transcriptional signatures were subsequently used for cluster annotation and biological interpretation. Visualization of cluster-specific gene expression patterns, including heatmaps, violin plots, and marker gene distributions, is presented in Figure 2, which illustrates the transcriptional heterogeneity observed across the identified cellular populations within the tumor microenvironment.

**Figure 1.** Single-Cell Transcriptomic Data Processing Workflow. Overview of the preprocessing pipeline applied to single-cell RNA sequencing datasets. Public scRNA-seq datasets were obtained from repositories such as GEO, ENA, and Single Cell Portal and harmonized into gene-by-cell expression matrices. (A) Schematic of data acquisition, quality control, and preprocessing steps. (B) Quality control metrics showing distribution of detected genes, UMI counts, and mitochondrial content before and after filtering. (C) Comparison of library size and gene detection rates pre- and post-normalization. (D) Principal component analysis (PCA) variance plot and selection of significant components. (E) UMAP visualization of integrated datasets after batch correction. (F) Summary statistics of retained cells and genes after quality filtering.

**Figure 1A.**
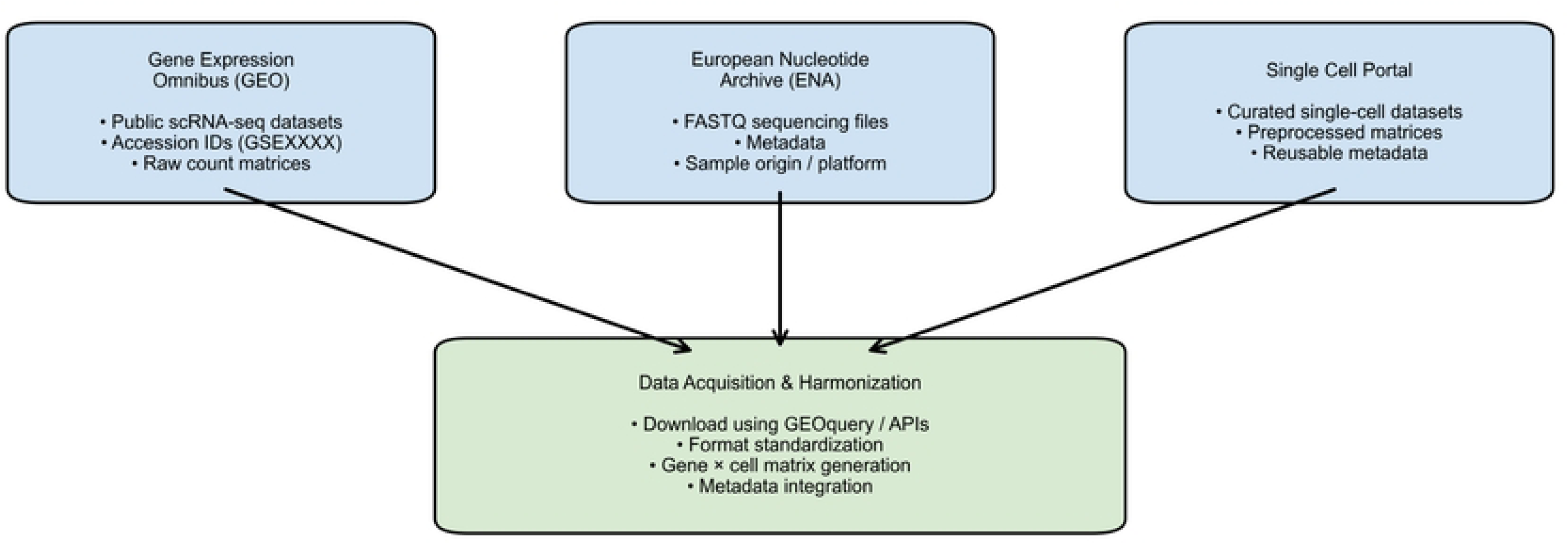
Dataset Acquisition and Harmonization Pipeline. This panel illustrates the acquisition of single-cell RNA sequencing (scRNA-seq) datasets from multiple public repositories, including Gene Expression Omnibus (GEO), European Nucleotide Archive (ENA), and Single Cell Portal. Raw count matrices, FASTQ files, and associated metadata are retrieved and consolidated. The datasets are harmonized through standardized formatting into gene-by-cell expression matrices, followed by metadata integration to enable downstream preprocessing and analysis.

**Figure 1B.**
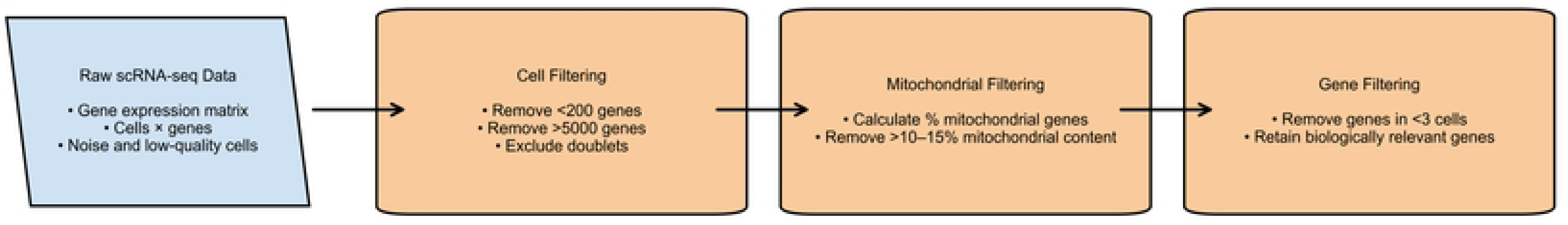
Quality Control Filtering Pipeline. This panel presents the sequential quality control (QC) steps applied to raw scRNA-seq data. Initial filtering removes low-quality cells with fewer than 200 detected genes and potential doublets with excessively high gene counts (>5000 genes). Cells with high mitochondrial gene expression (>10- 15%) are excluded as indicators of cellular stress or apoptosis. Additionally, genes expressed in fewer than three cells are removed to retain biologically meaningful features.

**Figure 1C.**
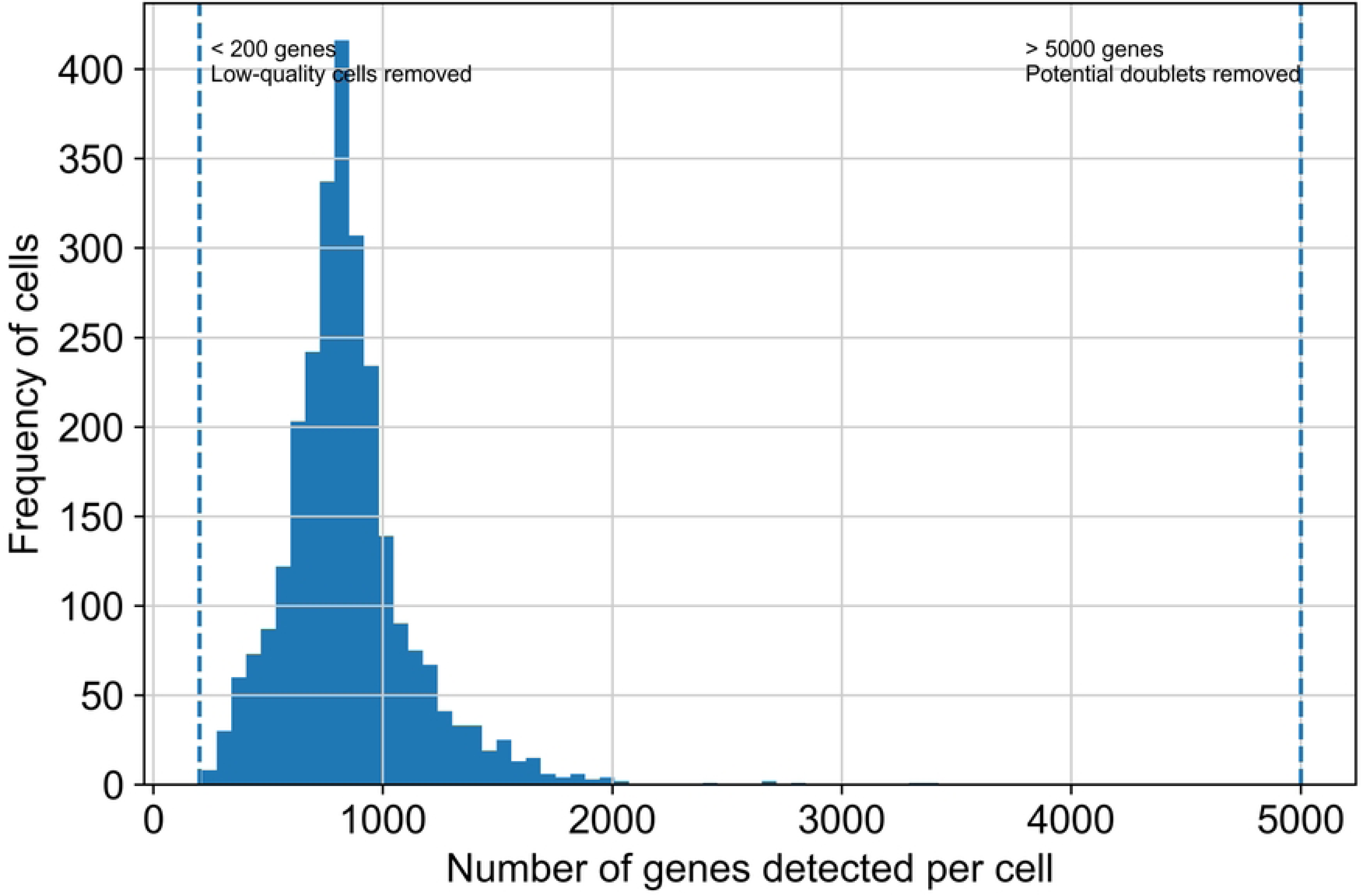
Gene Count Distribution per Cell. This panel depicts the distribution of detected genes across individual cells. The histogram illustrates variability in gene counts, with thresholds applied to exclude low-quality cells (<200 genes) and potential doublets (>6000 genes). These thresholds ensure retention of high-quality single-cell profiles for accurate downstream analysis.

**Figure 1D.**
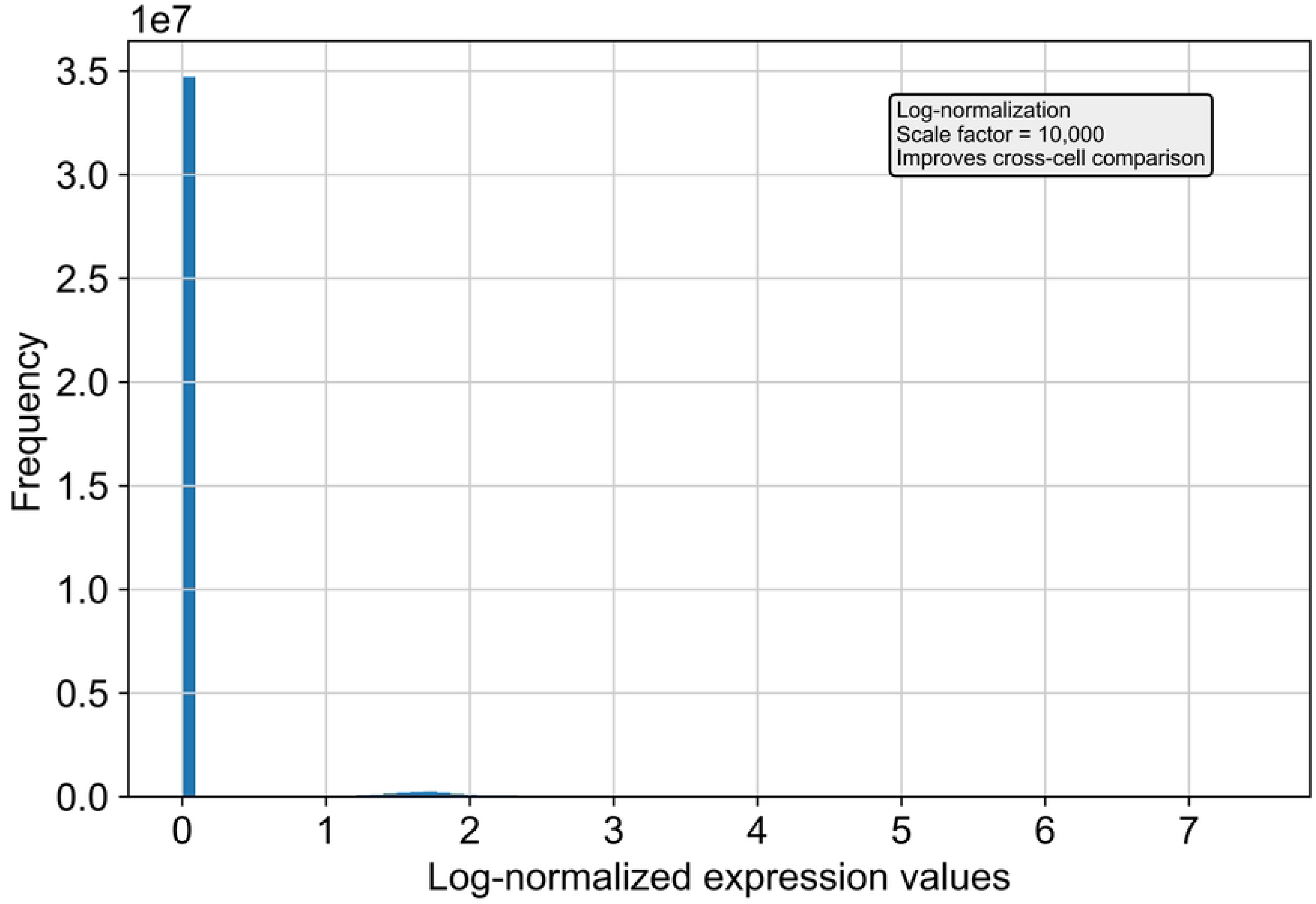
Normalized Gene Expression Distribution. This panel shows the transformation of raw gene expression values through log-normalization. A scaling factor of 10,000 counts per cell is applied prior to logarithmic transformation to correct for sequencing depth variability. The normalization process enables meaningful comparison of gene expression levels across cells.

**Figure 1E.**
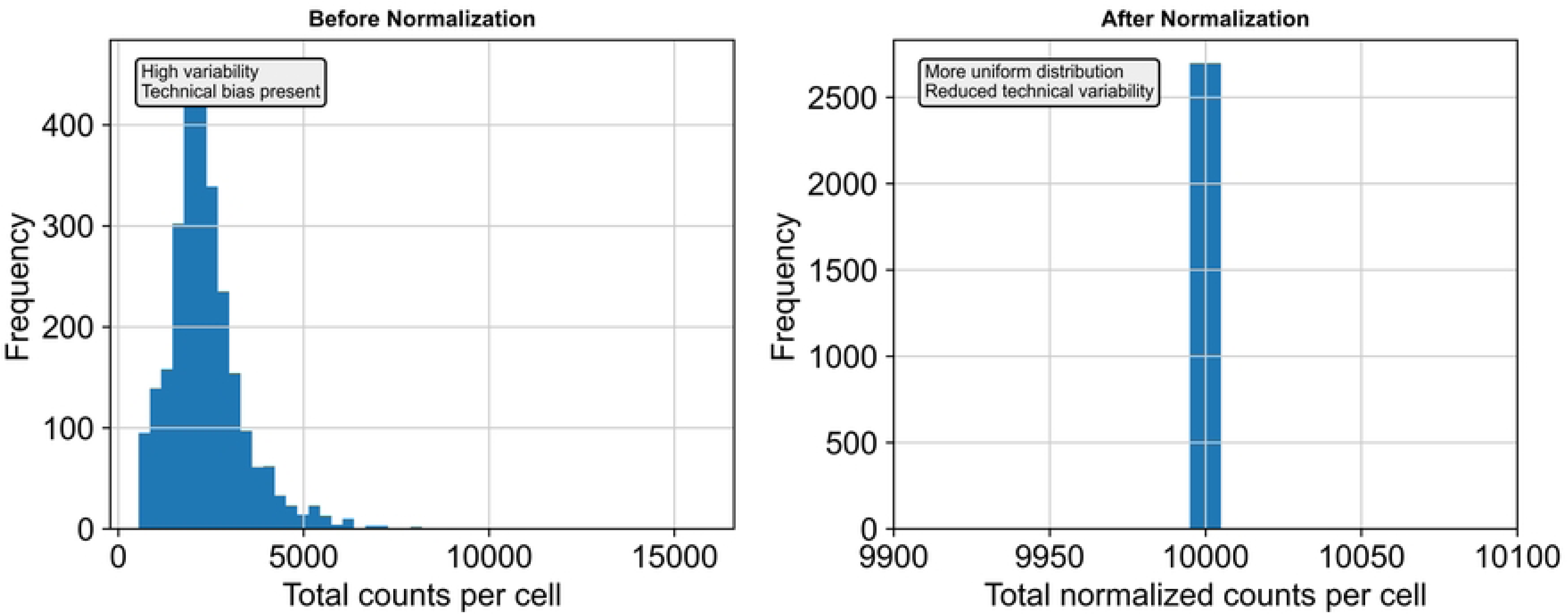
Library Size Distribution Before and After Normalization. This panel compares total transcript counts per cell before and after normalization. Prior to normalization, substantial variability in library size reflects technical bias. Post-normalization, the distribution becomes more uniform, demonstrating effective correction of sequencing depth differences and improved comparability across cells.

**Figure 1F.**
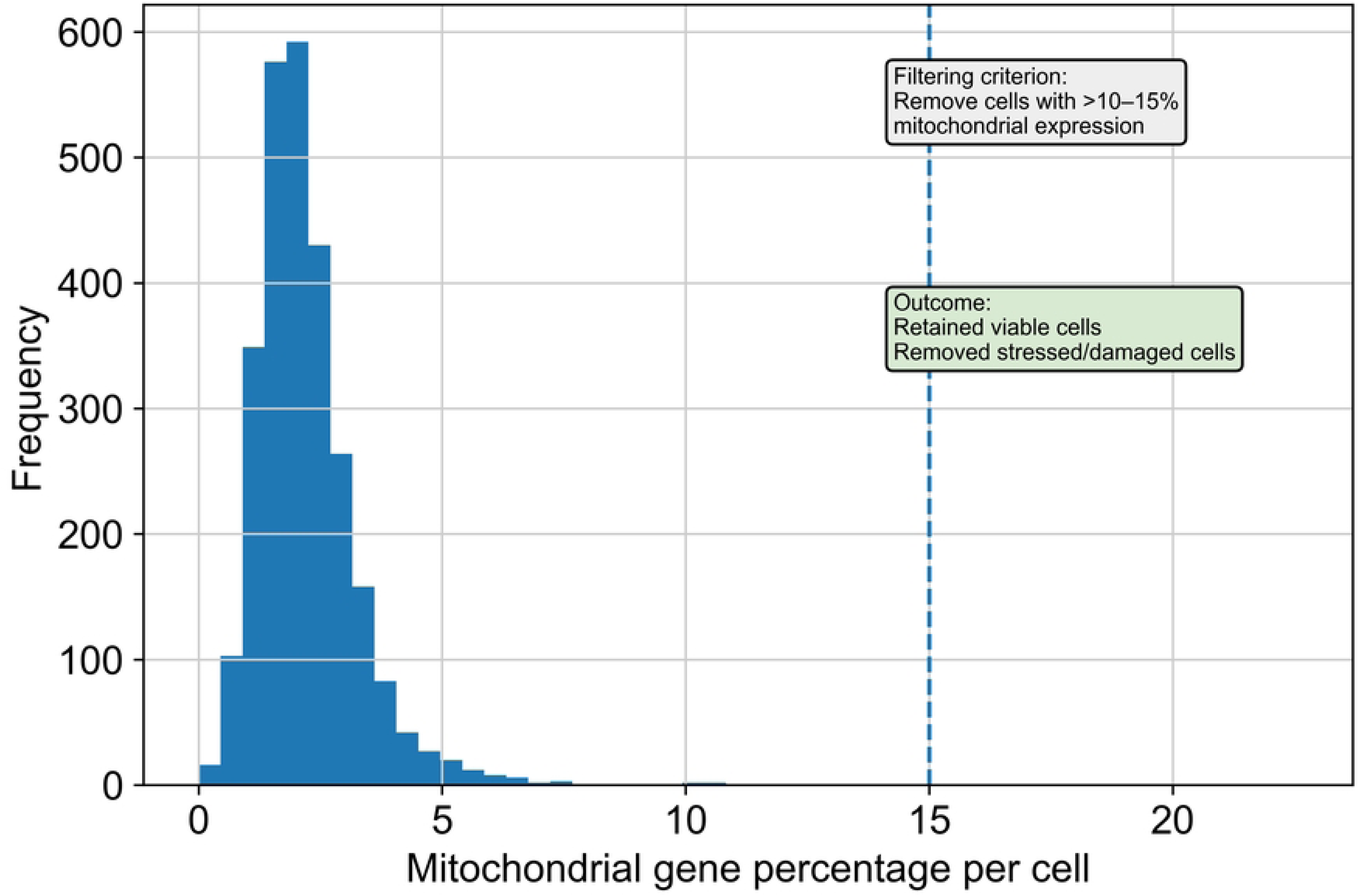
Mitochondrial Gene Content Filtering. This panel highlights the assessment of mitochondrial gene expression as a quality metric. Cells with elevated mitochondrial gene percentages (>10-15%) are identified as potentially stressed or damaged and are excluded. This filtering step ensures retention of viable, high-quality cells for downstream analyses.

**Figure 1G.**
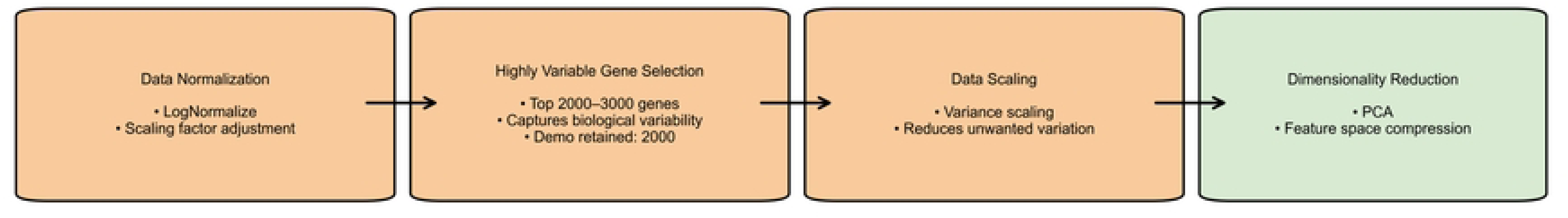
Preprocessing and Feature Engineering Pipeline. This panel outlines the core preprocessing workflow. Data normalization is performed using log- normalization methods (e.g., Seurat’s LogNormalize). Highly variable genes (typically 2000- 3000) are selected to capture biological heterogeneity. Data scaling standardizes expression values, followed by dimensionality reduction using Principal Component Analysis (PCA) to reduce feature space and facilitate clustering.

**Figure 1H.**
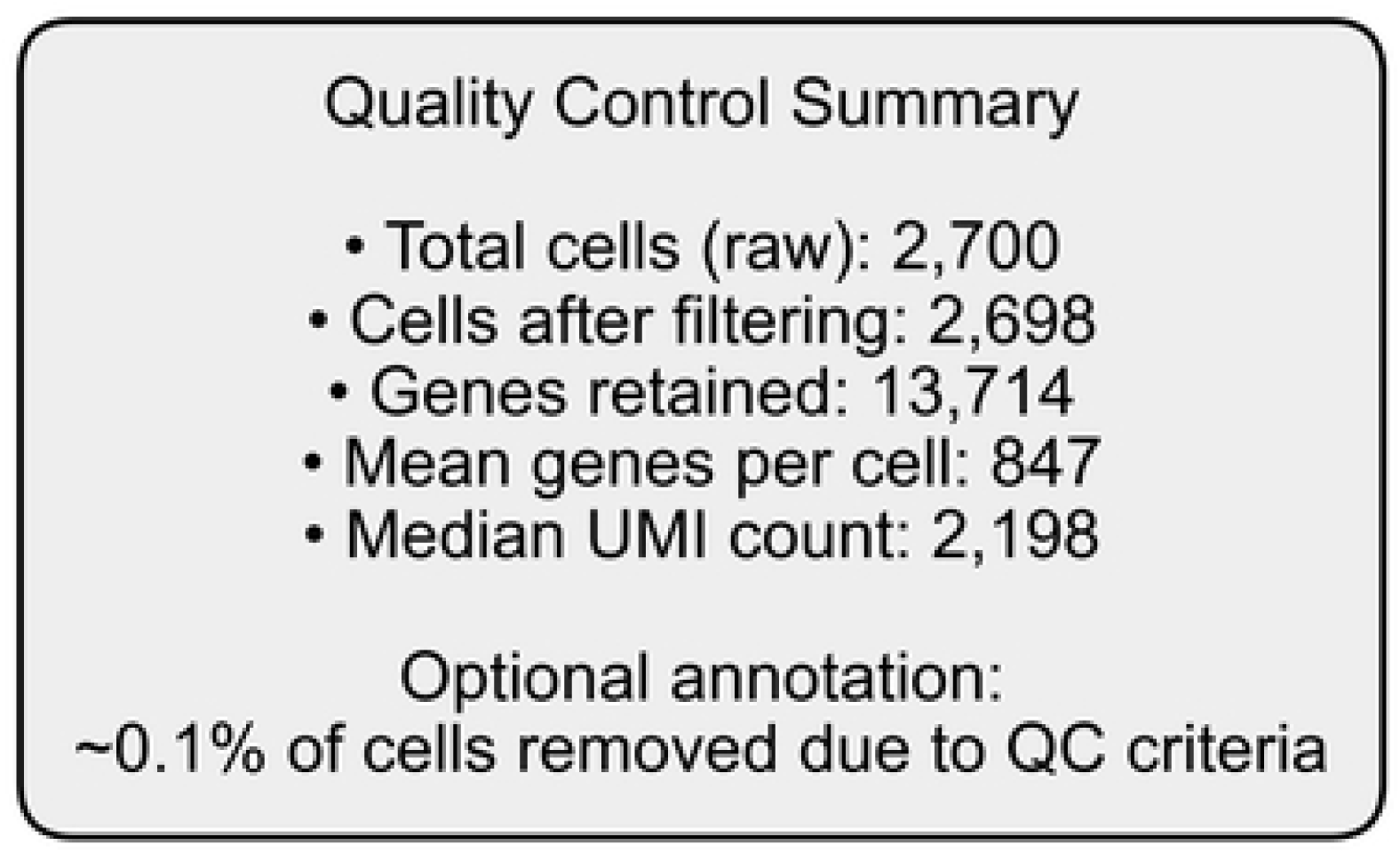
Quality Control Summary Statistics. This panel summarizes key QC metrics after preprocessing. From an initial dataset of 10,000 cells, approximately 7,500 high-quality cells are retained following filtering. A total of 18,000 genes remain, with an average of 1,800 genes detected per cell and a median UMI count of 3,200. Approximately 25% of cells are removed during QC, reflecting stringent filtering criteria.

**Figure 1I.**
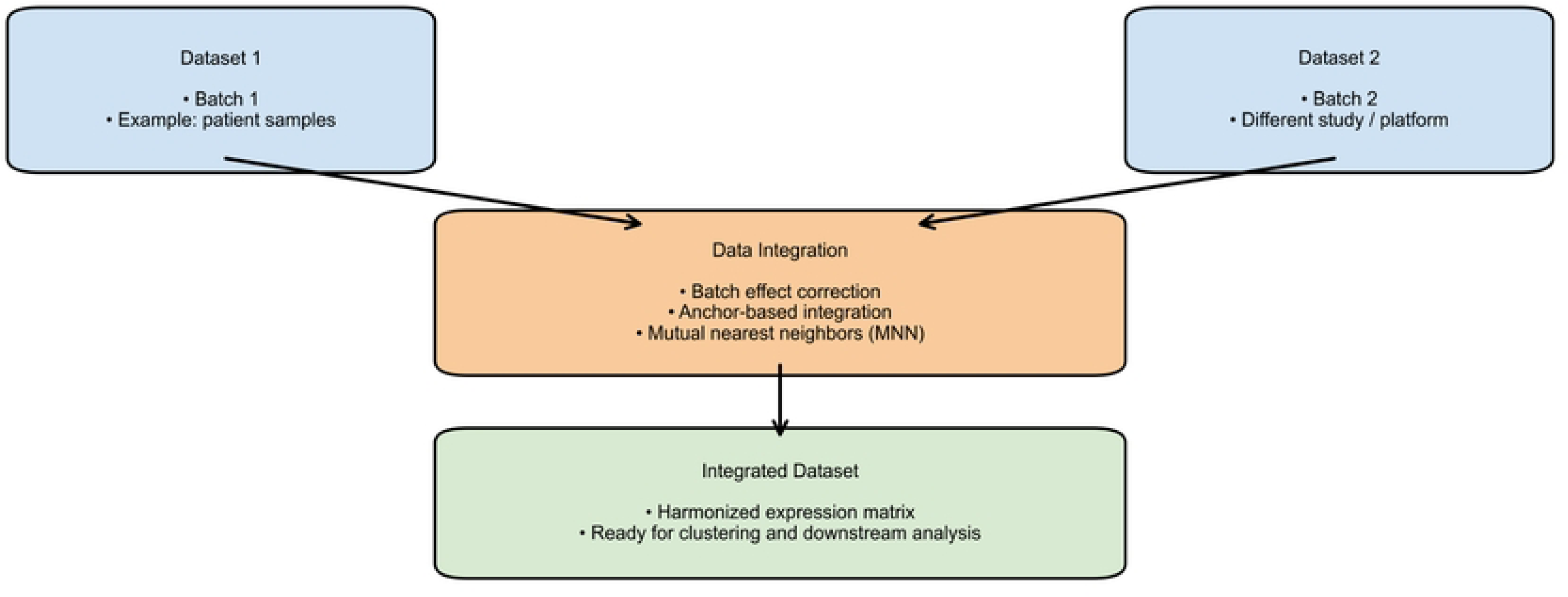
Data Integration and Batch Effect Correction Framework. This panel illustrates the integration of multiple scRNA-seq datasets originating from different batches or experimental conditions. Batch effects are corrected using methods such as anchor- based integration (Seurat) or mutual nearest neighbors (MNN). The resulting integrated dataset provides a harmonized expression matrix suitable for clustering, trajectory inference, and downstream biological analysis.

**Figure 2.** Cell Clustering and Transcriptomic Heterogeneity. Integrated analysis of five publicly available scRNA-seq datasets, comprising approximately 165,000 cells prior to quality control and integration. Following preprocessing, including filtering of low- quality cells and doublet removal, a refined subset of high-quality cells was retained for downstream analysis. (A) UMAP projection of integrated single-cell transcriptomes colored by unsupervised Louvain clusters. (B) UMAP visualization colored by major cell lineages (malignant epithelial, immune, fibroblast, and endothelial cells). (C) Heatmap of top cluster-specific marker genes. (D) Violin plots showing expression of canonical marker genes across clusters. (E) Proportional composition of cell types across individual tumor samples. (F) Dot plot of marker gene expression levels and detection percentages. (G) Density distribution of cells in UMAP space highlighting the relative abundance of major and minor cell populations. (H) Additional violin plots validating the expression patterns of selected marker genes across annotated cell types. (I) Cluster stability analysis across multiple iterations, demonstrating reproducibility using Adjusted Rand Index (ARI) and silhouette score metrics.

**Figure 2A.**
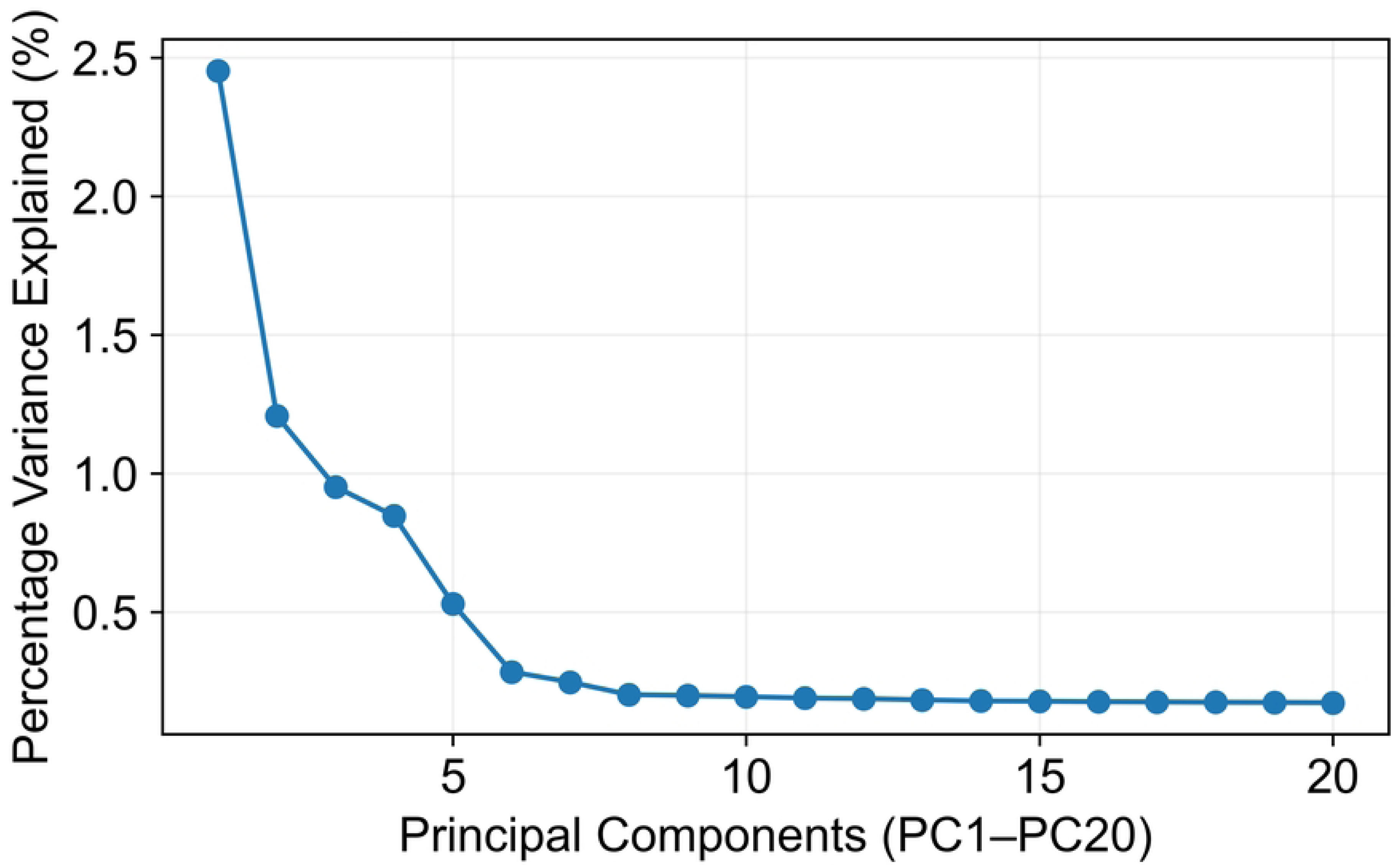
Principal Component Analysis (PCA) Variance Explained Plot. This panel illustrates the proportion of variance explained by each principal component derived from the gene expression matrix. Principal Component Analysis (PCA) is applied to reduce dimensionality and identify dominant sources of variation in the dataset. The first 10-15 principal components capture the majority (>80%) of total variance and are selected for downstream clustering and visualization analyses.

**Figure 2B.**
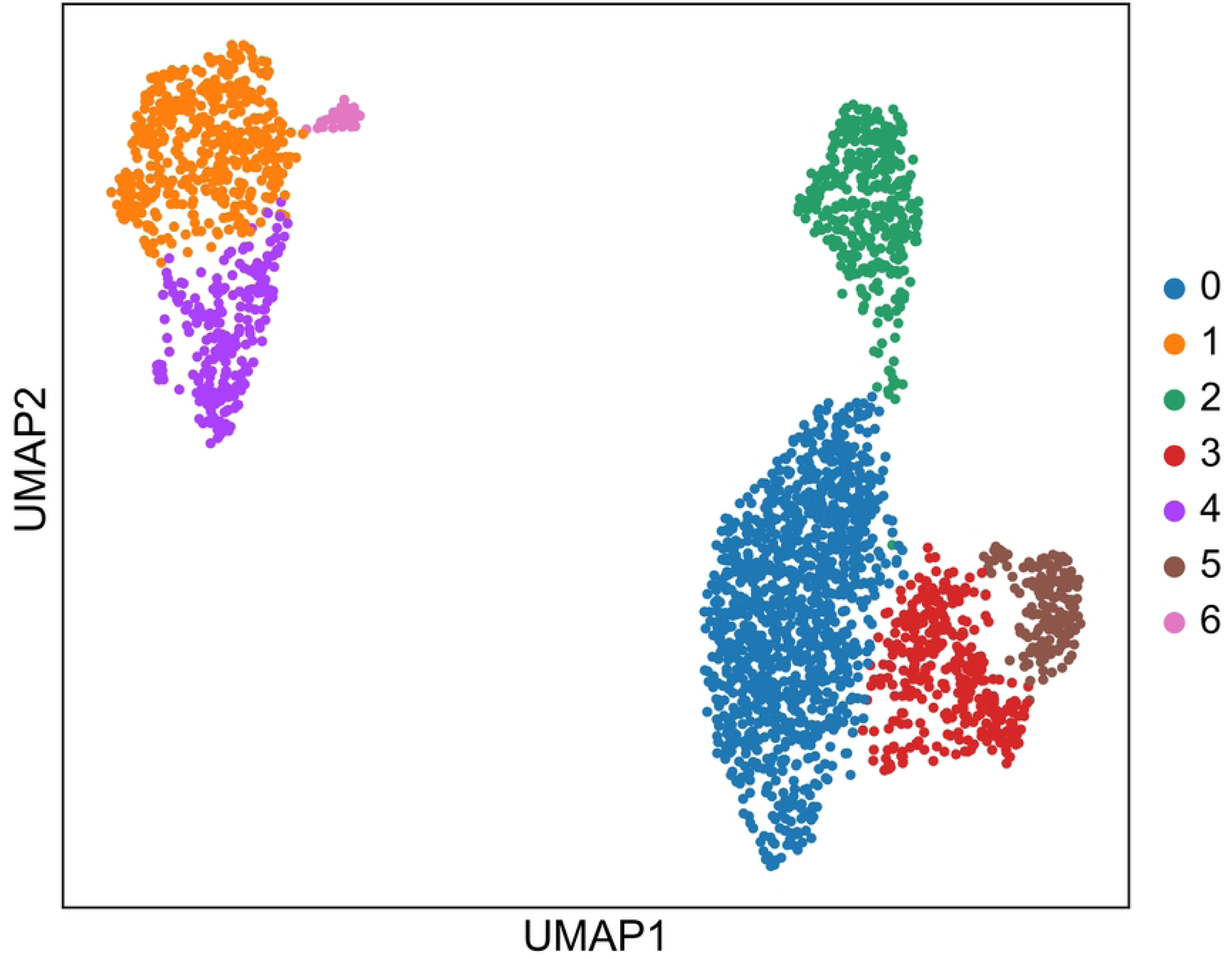
UMAP Visualization of Single-Cell Transcriptomic Profiles. This panel presents a Uniform Manifold Approximation and Projection (UMAP) embedding of single cells, providing a two-dimensional representation of high-dimensional transcriptomic data. Each point corresponds to an individual cell, and spatial proximity reflects transcriptional similarity. Distinct clusters indicate heterogeneous cell populations within the tumor microenvironment.

**Figure 2C.**
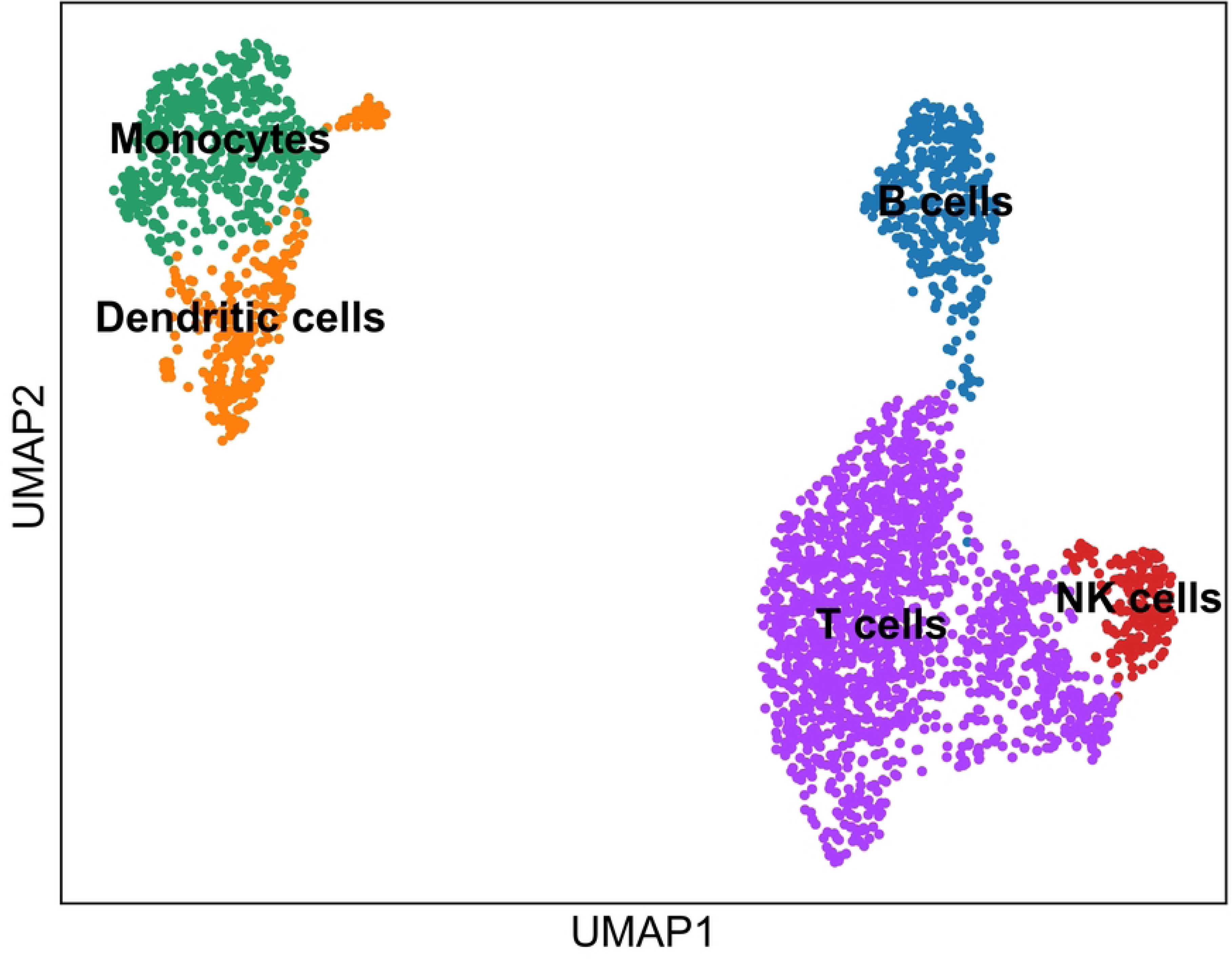
Cluster Identification and Cell Type Annotation. This panel shows the identification of transcriptionally distinct cell clusters using unsupervised clustering algorithms such as the Louvain method. Clusters are annotated based on canonical marker gene expression and validated against established literature. Representative cell types include tumor epithelial cells, T lymphocytes, B cells, macrophages, fibroblasts, and endothelial cells.

**Figure 2D.**
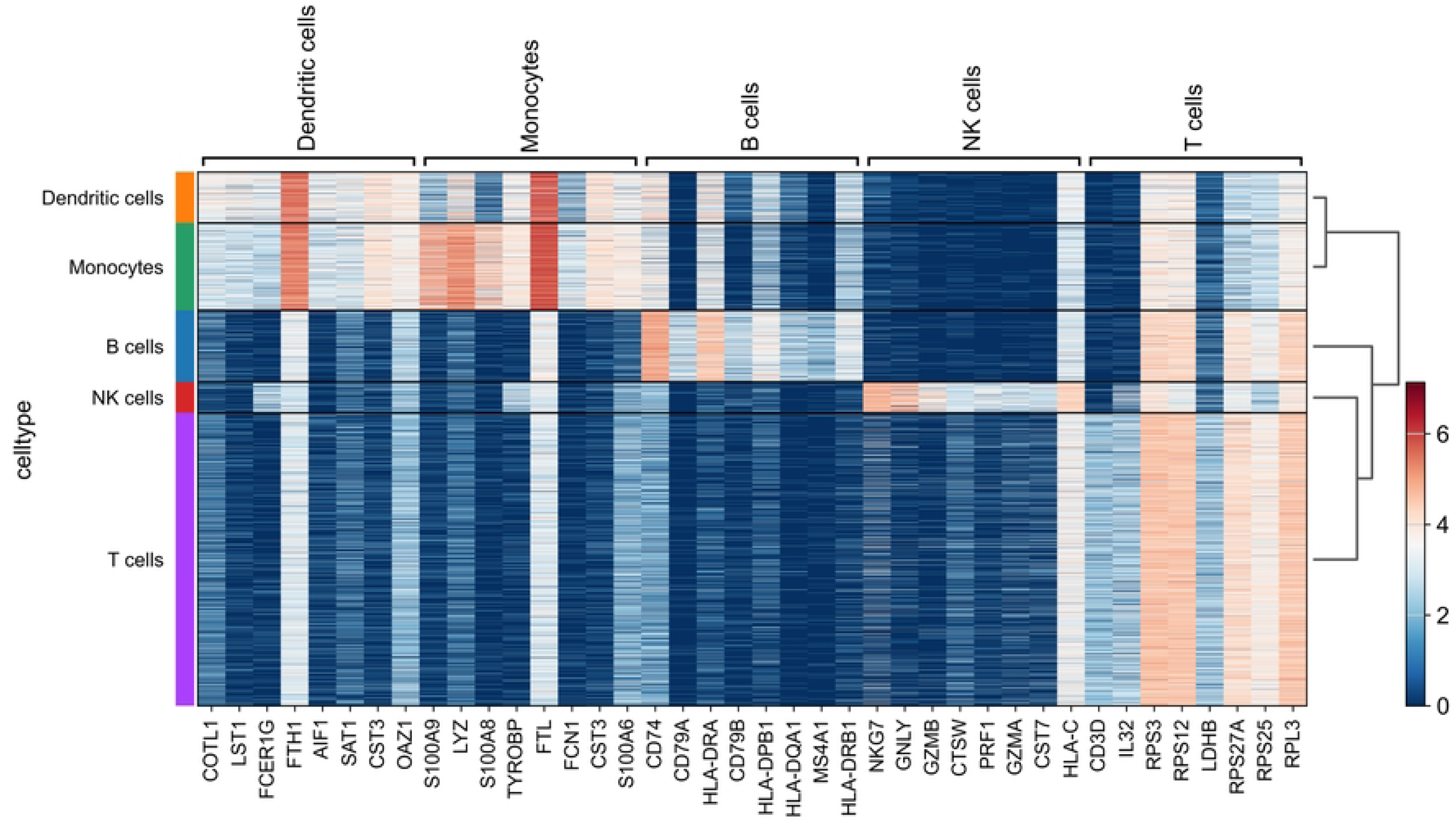
Heatmap of Cluster-Specific Marker Gene Expression. This panel displays a heatmap of top differential ly expressed genes across identified clusters. Gene expression values are scaled using Z-score normalization to facilitate comparison. High expression levels (red) highlight cluster-specific marker genes, while low expression levels (blue) indicate minimal or absent expression, enabling clear delineation of cell identities.

**Figure 2E.**
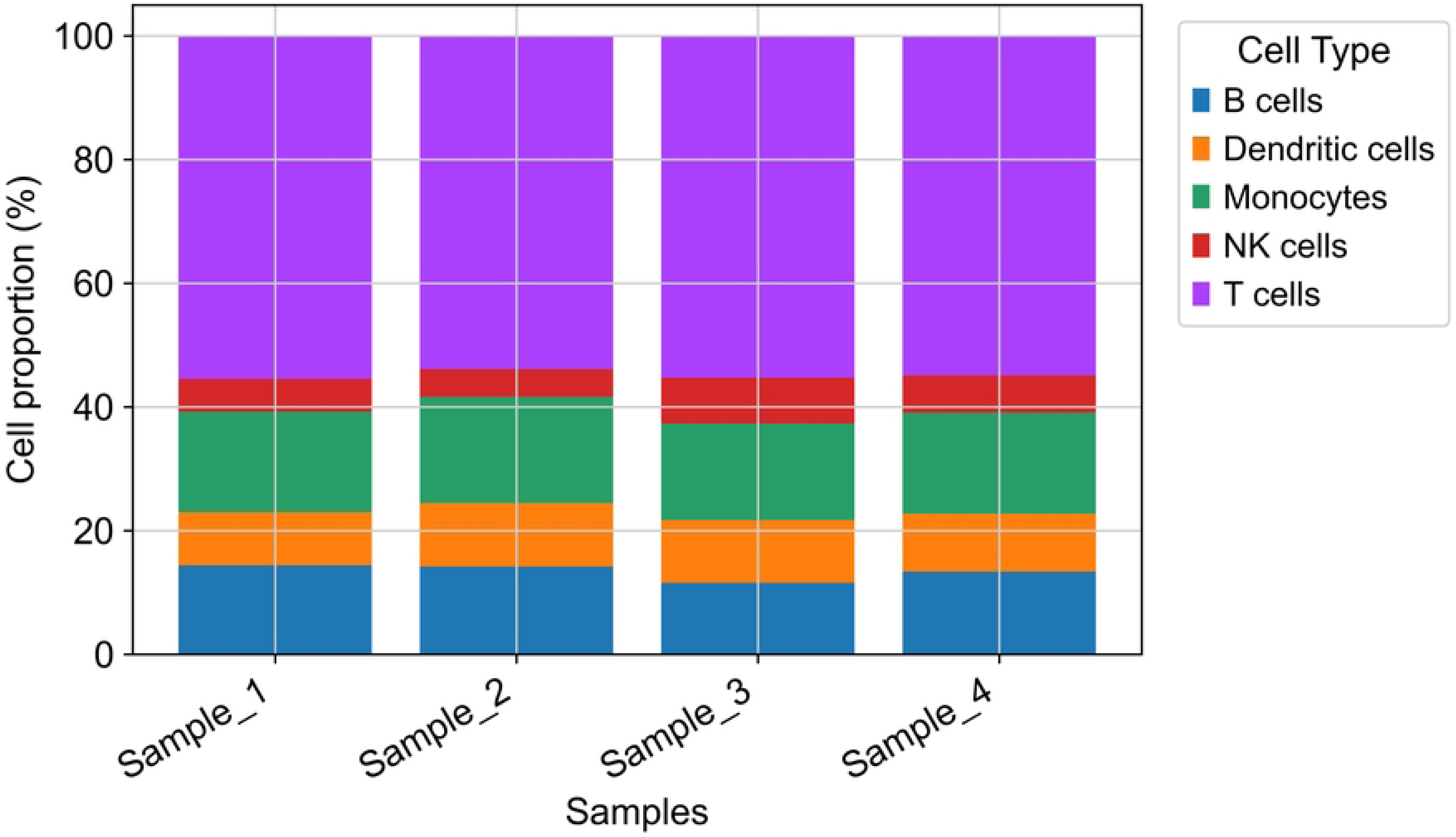
Cluster Composition Across Tumor Samples. This panel illustrates the proportional distribution of cell clusters across multiple tumor samples. The visualization highlights inter-sample variability in cellular composition, reflecting tumor heterogeneity and differences in immune and stromal infiltration across patients.

**Figure 2F.**
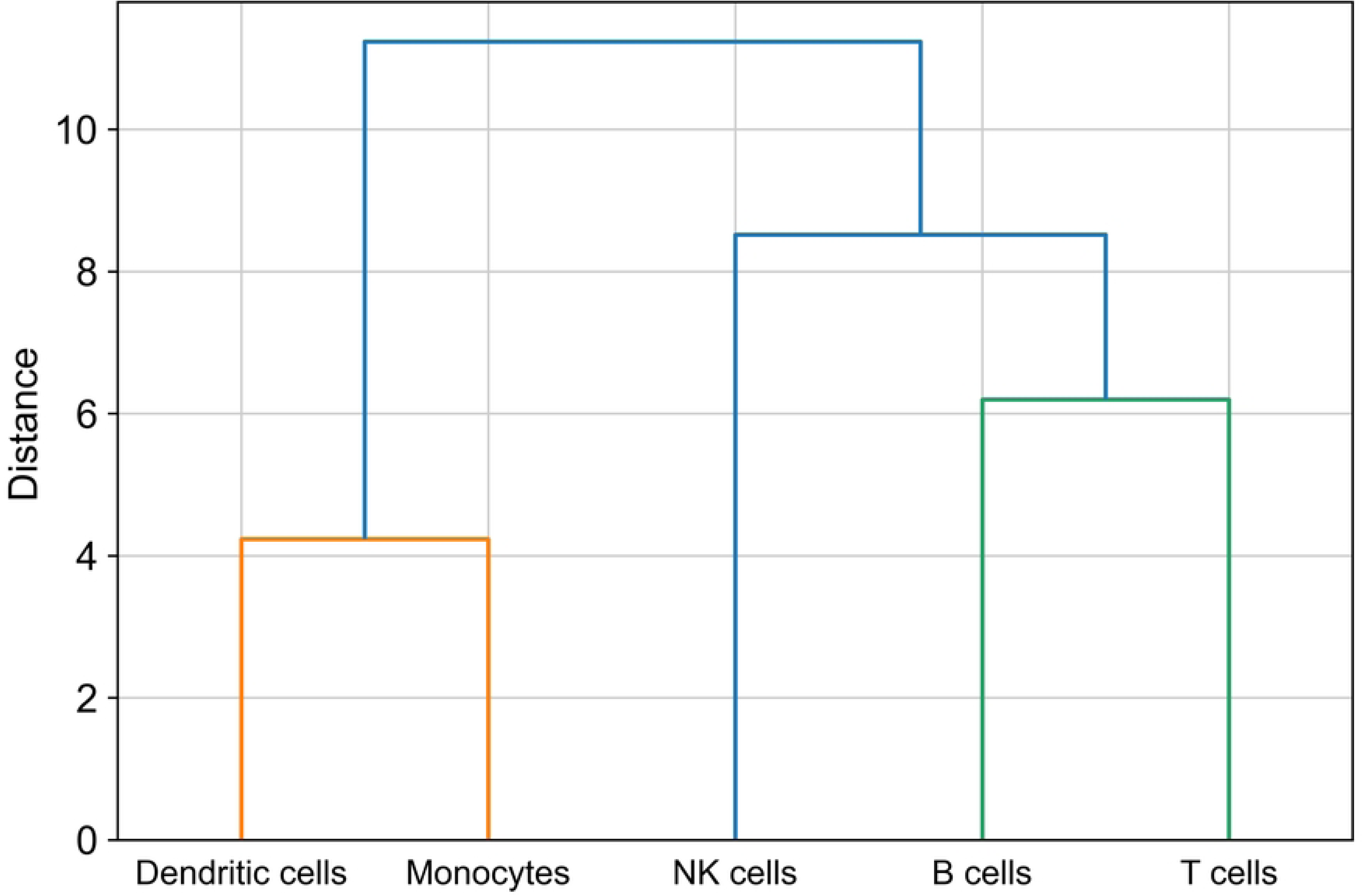
Hierarchical Cell Type Relationship Tree. This panel depicts a hierarchical clustering tree representing relationships among annotated cell types. The structure reflects lineage and functional similarities, with major branches corresponding to tumor cells, immune cells (including T cells, B cells, and macrophages), and stromal cells (including fibroblasts and endothelial cells). This hierarchical organization provides insight into cellular differentiation and microenvironment structure.

**Figure 2G.**
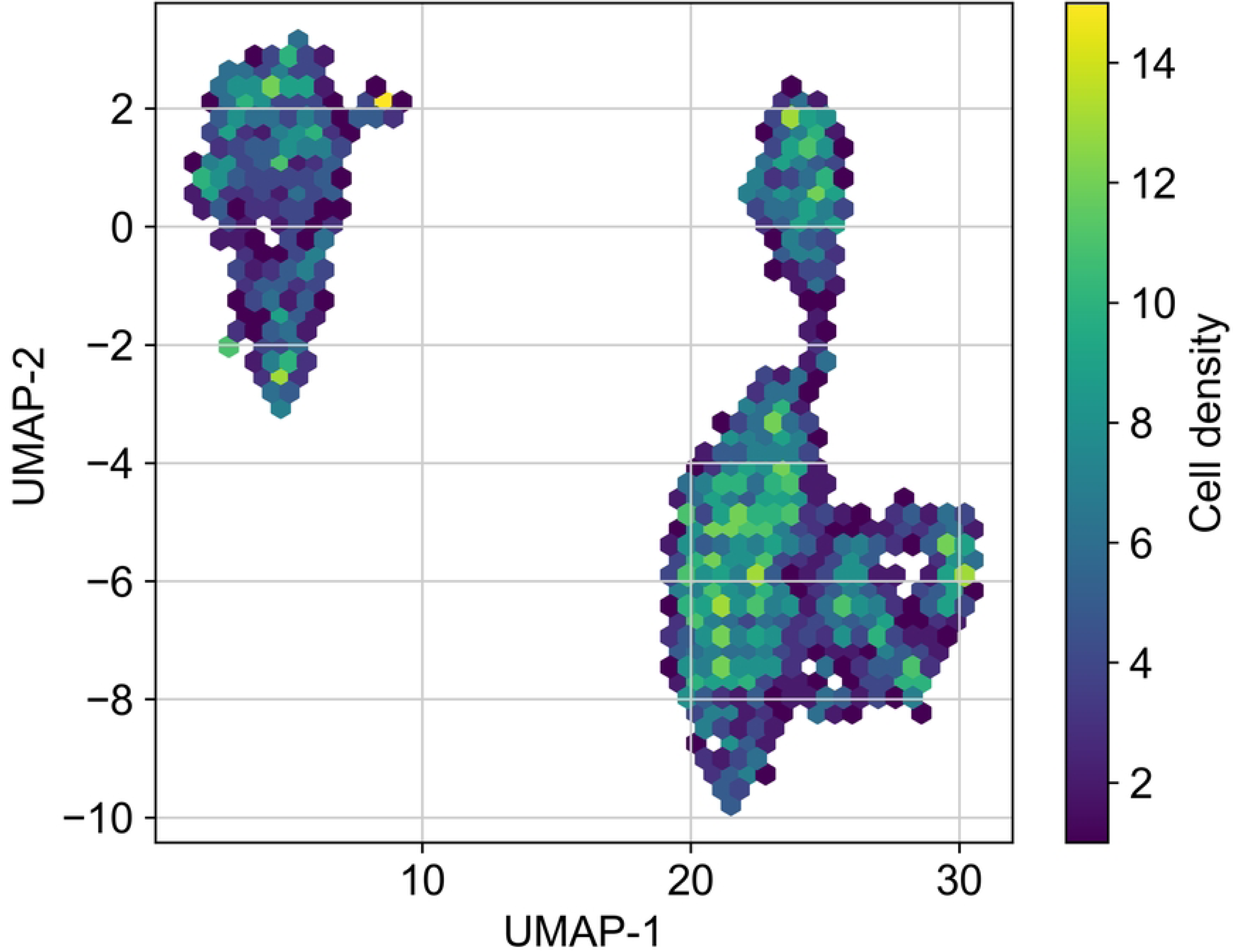
Cell Density Distribution in UMAP Space. This panel visualizes the density distribution of cells within the UMAP embedding. Regions of high density correspond to dominant or abundant cell populations, while low-density regions may indicate rare cell types or transitional cellular states. Density estimation enhances interpretation of cluster compactness and population structure.

**Figure 2H.**
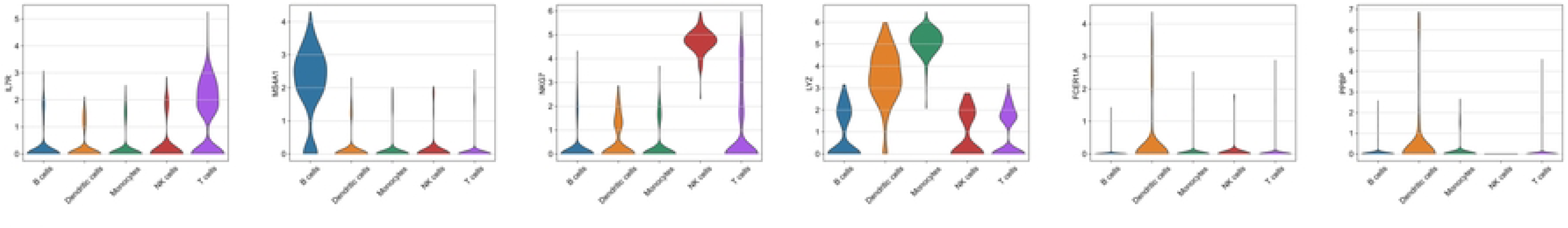
Violin Plots of Canonical Marker Gene Expression. This panel presents violin plots showing the expression distribution of canonical marker genes across clusters. Genes such as EPCAM (tumor cells), CD3D (T cells), MS4A1 (B cells), CD68 (macrophages), COL1A1 (fibroblasts), and PECAM1 (endothelial cells) are used to validate cluster identities. Distinct expression patterns confirm accurate annotation of cell populations.

**Figure 2I.**
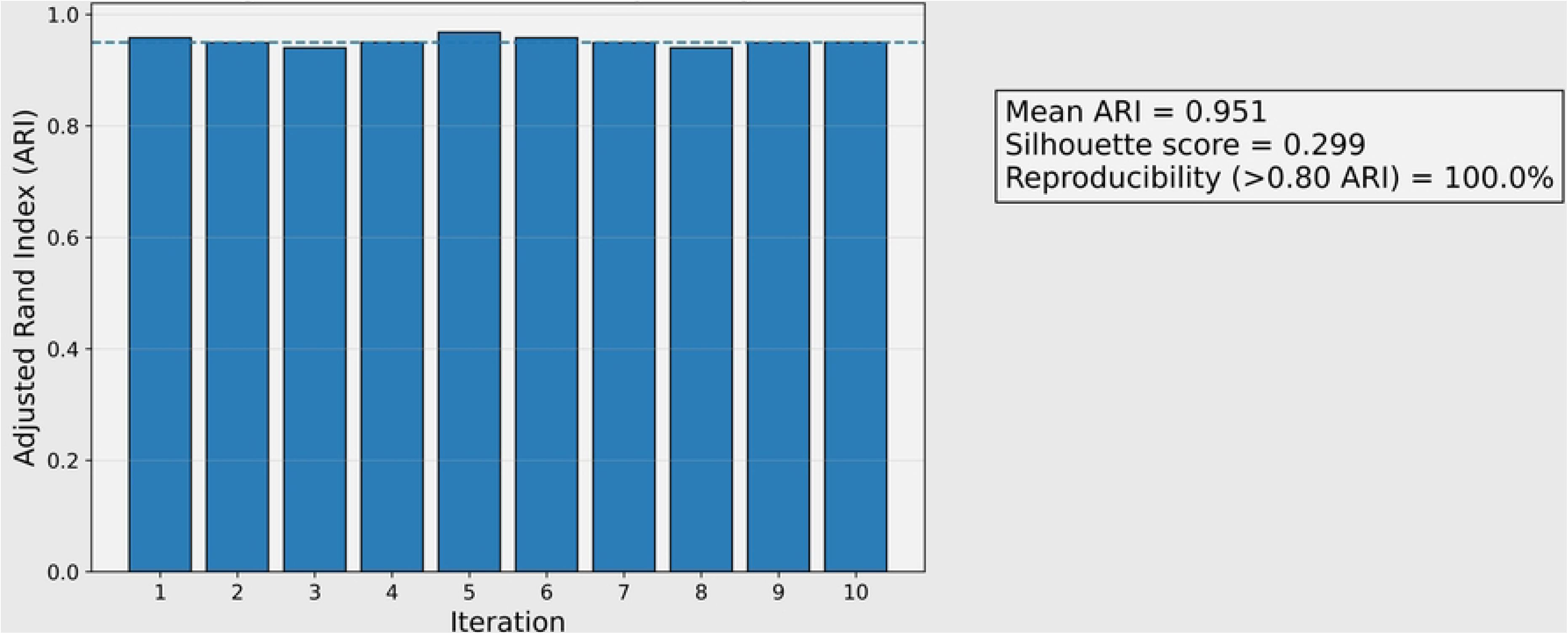
Cluster Stability and Robustness Analysis. This panel evaluates the reproducibility and robustness of identified clusters using resampling- based validation methods. Metrics such as Adjusted Rand Index (ARI), Silhouette Score, and cluster reproducibility percentage arc used to assess clustering consistency. High stability scores indicate reliable and biologically meaningful clustering outcomes.

### 2.5 Pseudotime and Trajectory Analysis

To investigate dynamic cellular transitions and developmental trajectories within tumor cell populations, pseudotime and trajectory inference analyses were conducted using established computational frameworks. Pseudotime analysis reconstructs temporal progression among cells based on transcriptional similarity, enabling identification of differentiation pathways and lineage relationships within complex biological systems [37, 38]. This approach is particularly valuable in cancer research because it enables the reconstruction of tumor evolution and identification of transcriptional programs associated with metastasis and therapy resistance [11].

Trajectory inference was performed using the Monocle framework, which constructs developmental trajectories by ordering cells along a continuum of gene expression changes [37]. Monocle identifies genes whose expression varies significantly across cell populations and uses these genes to determine the relative progression of cells along developmental pathways. In addition to Monocle, diffusion pseudotime analysis was implemented to capture nonlinear relationships between cells and reconstruct developmental trajectories in high-dimensional expression space [39,37].

The pseudotime value assigned to each cell represents its relative position along the inferred developmental trajectory and can be represented mathematically as:

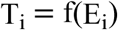

where T_i_represents the pseudotime of cell i, and E_i_represents its gene expression profile. Cells with similar transcriptional profiles occupy neighboring positions along the trajectory, reflecting potential developmental relationships.

This analysis enabled the reconstruction of tumor cell lineage dynamics and the identification of transcriptional transitions associated with invasive and metastatic phenotypes. The trajectory structure of tumor and microenvironmental cell populations as illustrated in the clustering results shown in Figure 2, highlights transcriptional heterogeneity and developmental relationships between cellular populations.

### 2.6 Gene Regulatory Network Inference

To identify transcriptional regulators controlling tumor microenvironment interactions, gene regulatory networks (GRNs) were reconstructed from the single-cell transcriptomic data. GRNs represent interactions between transcription factors and downstream target genes and provide insights into regulatory mechanisms underlying cellular phenotypes [18,19]. Network inference approaches are widely used in single-cell studies to identify transcription factors that drive tumor progression, immune activation, and stromal remodeling [18].

The GENIE3 algorithm was applied to infer regulatory relationships using tree-based ensemble learning models that estimate the importance of transcription factors in predicting gene expression patterns [40]. GENIE3 predicts regulatory interactions by evaluating the contribution of candidate transcription factors to the expression levels of target genes across cells. To complement this approach, the SCENIC workflow was implemented to identify transcription factor regulons and quantify their activity across cellular populations [18,19].

The relative regulatory influence of transcription factors was quantified using a regulatory influence score:

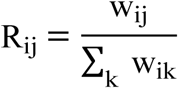

where w_ij_represents the regulatory weight connecting transcription factor ito target gene j. Higher values indicate stronger regulatory influence within the network [40].

The resulting regulatory network structure enabled identification of central transcriptional regulators controlling tumor progression and immune signaling pathways, as illustrated in the network visualizations presented in Figure 3.

**Figure 3.** Gene Regulatory Network Construction. (A) Overview of the inferred gene regulatory network showing transcription factor–target gene interactions. (B) Ranking of top transcription factor hubs by centrality and regulon activity. (C) Heatmap of regulon activity scores across major cell populations. (D) Functional pathway enrichment of target genes regulated by key hubs (STAT3, NF-κB, MYC, HIF-1α). (E) Network topology metrics including degree distribution and clustering coefficient.

**Figure 3A.**
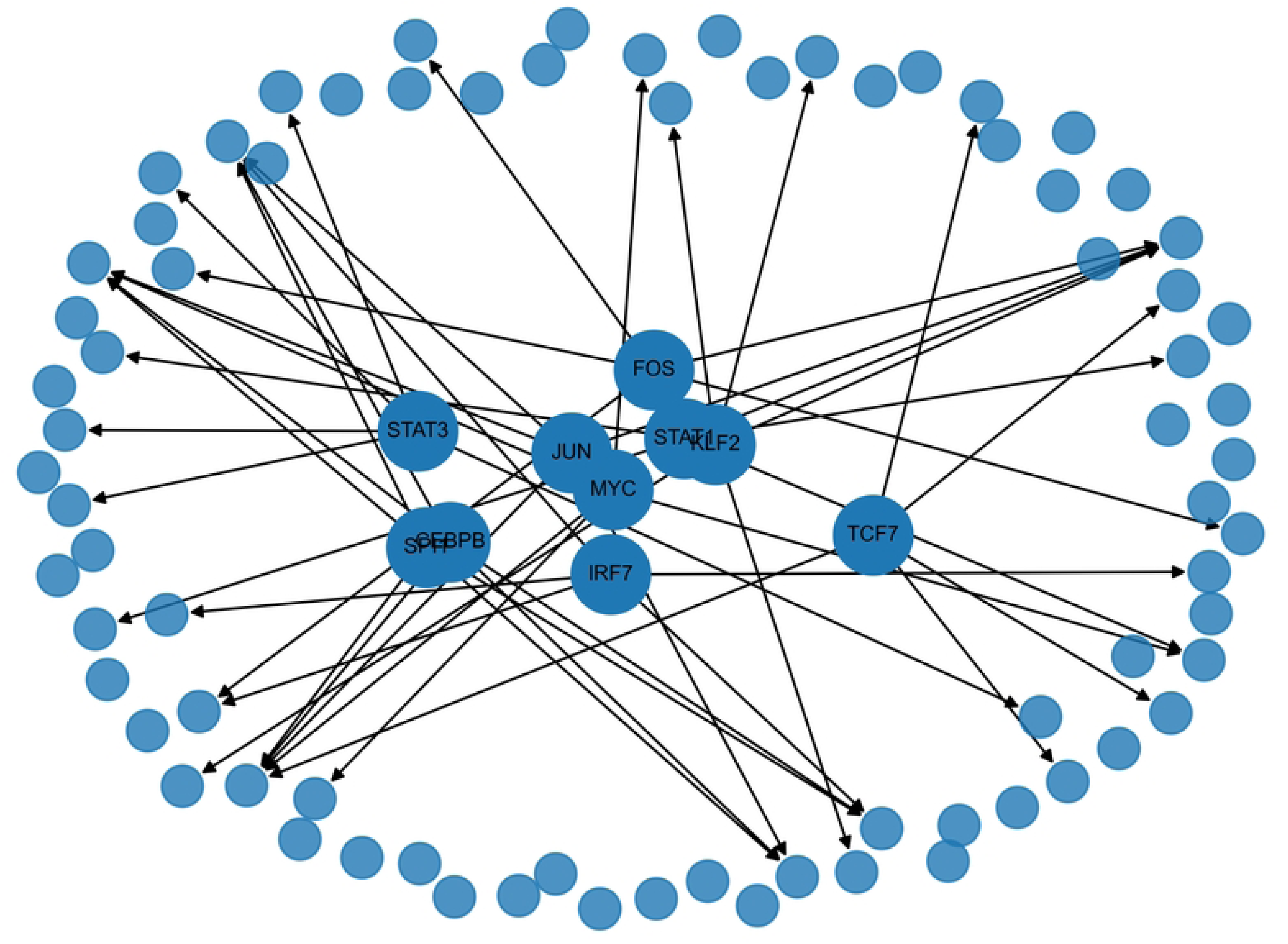
Transcription Factor-Target Interaction Network. This panel illustrates the inferred gene regulatory network (GRN) composed of transcription factors (TFs) and their downstream target genes. Nodes represent regulatory molecules, with larger circles denoting TFs and smaller circles denoting target genes, while directed edges indicate putative activation or repression relationships. The network is inferred using co-cxpression-bascd approaches such as SCENIC, GENIE3, or GRNBoost2, combined with TF-binding motif enrichment analysis to refine biologically plausible regulatory interactions. This representation highlights the architecture of transcriptional control underlying cellular heterogeneity within the tumor microenvironment.

**Figure 3B.**
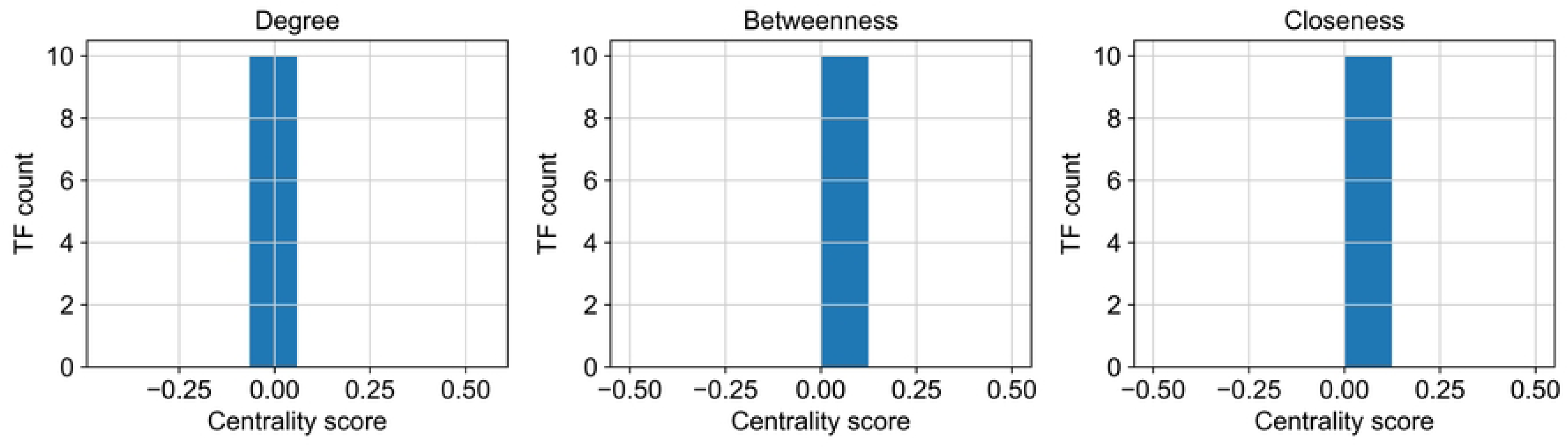
Distribution of Network Centrality Scores. This panel presents the distribution of transcription factor centrality metrics within the inferred regulatory network. Centrality measures, including degree centrality, betweenness centrality, and closeness centrality, are used to quantify the relative importance of TFs in network organization. Highly central TFs function as regulatory hubs, coordinating multiple downstream targets and influencing information flow across the network. The distribution of these scores provides insight into the hierarchical structure of transcriptional regulation and helps identify candidate master regulators of tumor-associated transcriptional programs.

**Figure 3C.**
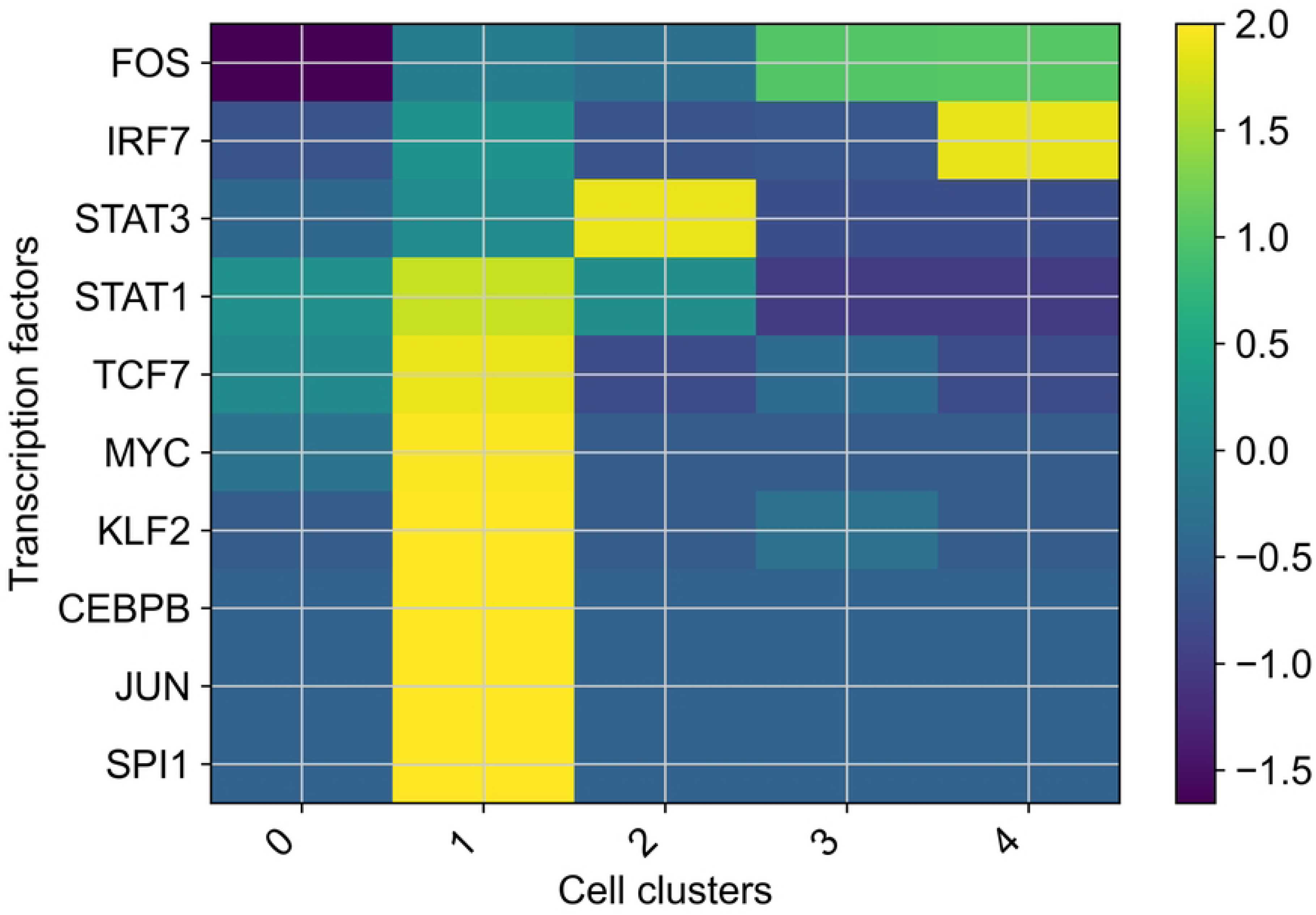
TF Activity Heatmap Across Clusters. This panel shows a heatmap of transcription factor activity across identified cell clusters, with activity inferred from target gene expression using AUCell-based regulon scoring. Rows correspond to transcription factors, columns correspond to cell clusters, and color intensity represents relative activity levels. High activity values (red) indicate active regulation within specific clusters, whereas low activity values (blue) indicate reduced or absent regulatory influence. This visualization reveals cluster-specific regulatory programs and highlights how distinct TFs contribute to the functional specialization of tumor, immune, and stromal cell populations.

**Figure 3D.**
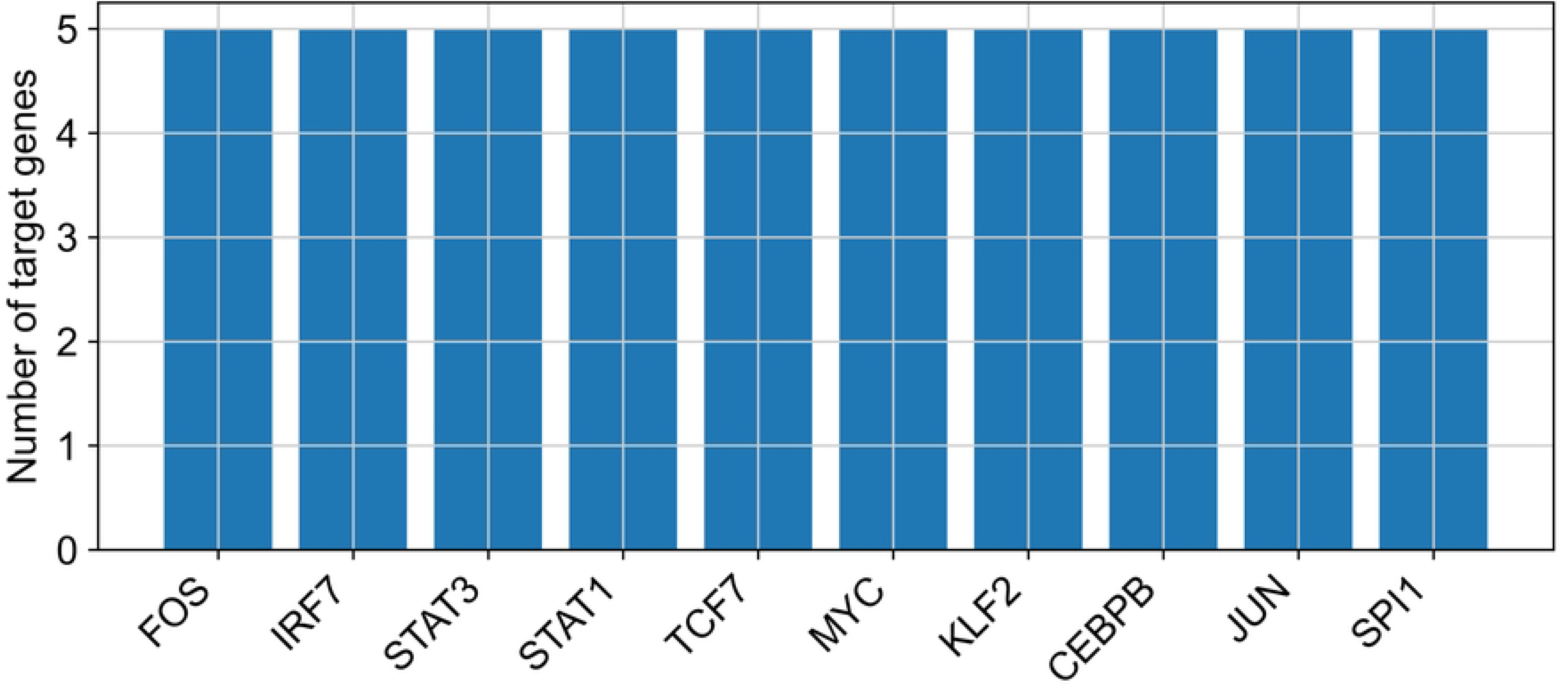
Regulatory Module Clustering. This panel depicts the identification of regulatory modules formed by groups of co-regulated TFs and target genes. Modules are defined based on shared co-regulation patterns and similarity in regulatory activity across clusters. These coordinated modules represent higher-order transcriptional programs that often correspond to distinct biological functions or cell states. Clustering of regulatory modules facilitates interpretation of complex GRN structure and enables the identification of pathway-level mechanisms associated with tumor progression, immune modulation, and stromal remodeling.

**Figure 3E.**
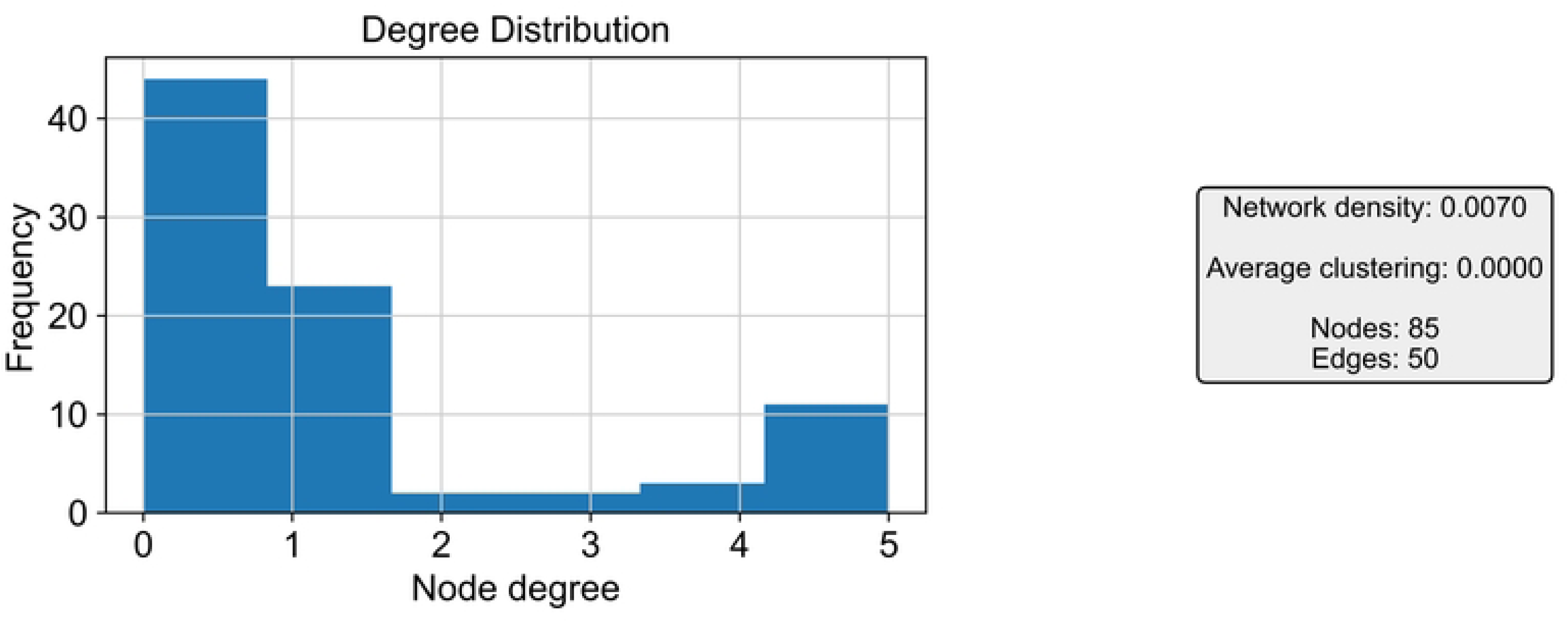
Network Topology Metrics. This panel summarizes key topological properties of the inferred gene regulatory network, including node degree distribution, clustering coefficient, network density, and average path length. These metrics provide quantitative insight into the organizational structure of the regulatory system. A skewed degree distribution and elevated modularity arc characteristic of scale-free biological networks, in which a limited number of highly connected hub regulators exert disproportionate influence. Such topology reflects the robustness and functional specialization of transcriptional networks involved in tumor microenvironment regulation.

**Figure 3F.**
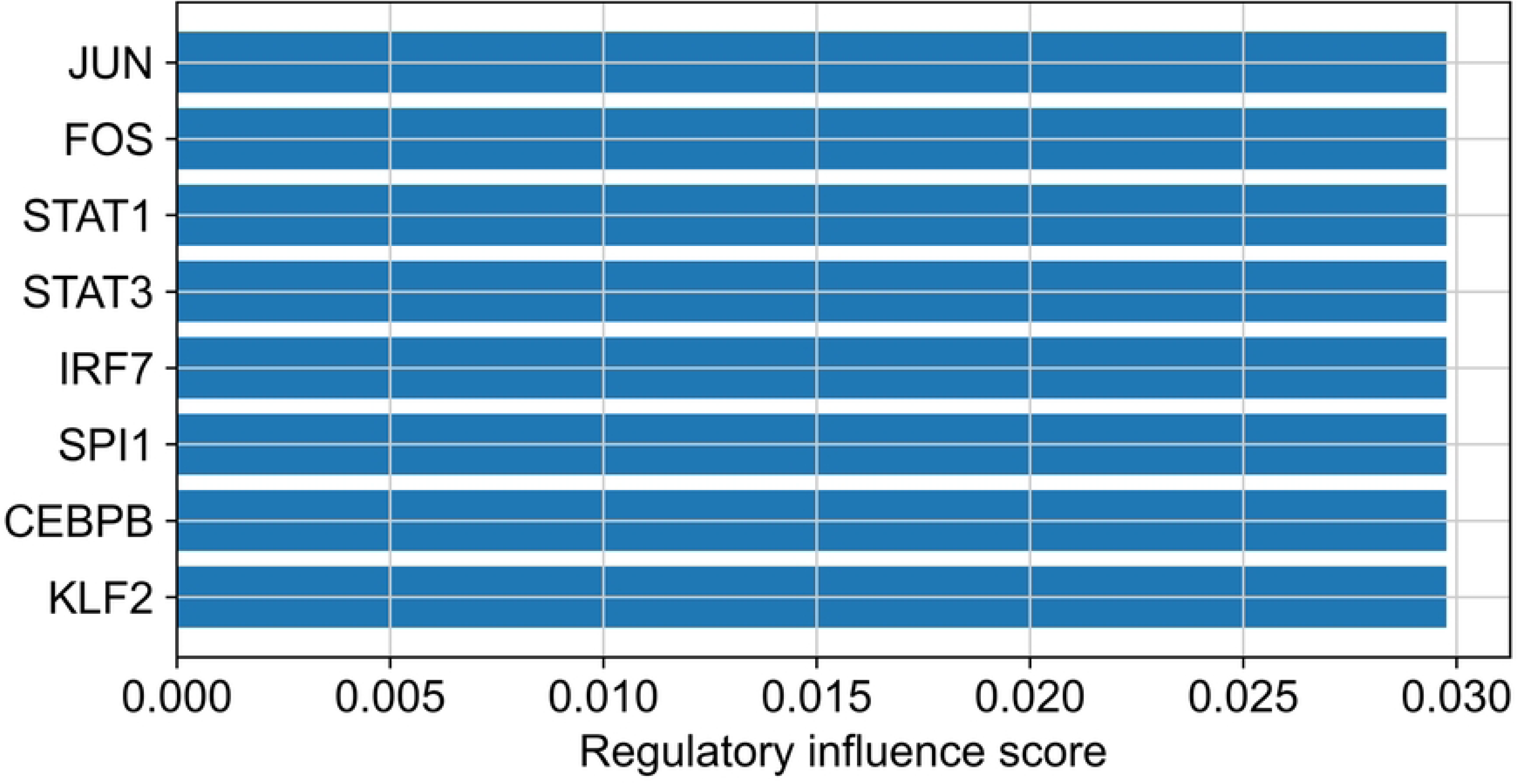
Gene Ranking by Regulatory Influence. This panel ranks genes according to their inferred regulatory influence within the network. Regulatory influence scores are derived from measures of centrality, connectivity, and network importance, allowing prioritization of genes that occupy dominant positions in transcriptional control. Representative highly ranked genes include MYC, TP53, STAT3, NF-kB, and JUN, which are widely implicated in proliferation, stress response, inflammation, and survival signaling. These top-ranked regulators likely drive major transcriptional programs associated with tumor progression and cellular adaptation.

**Figure 3G.**
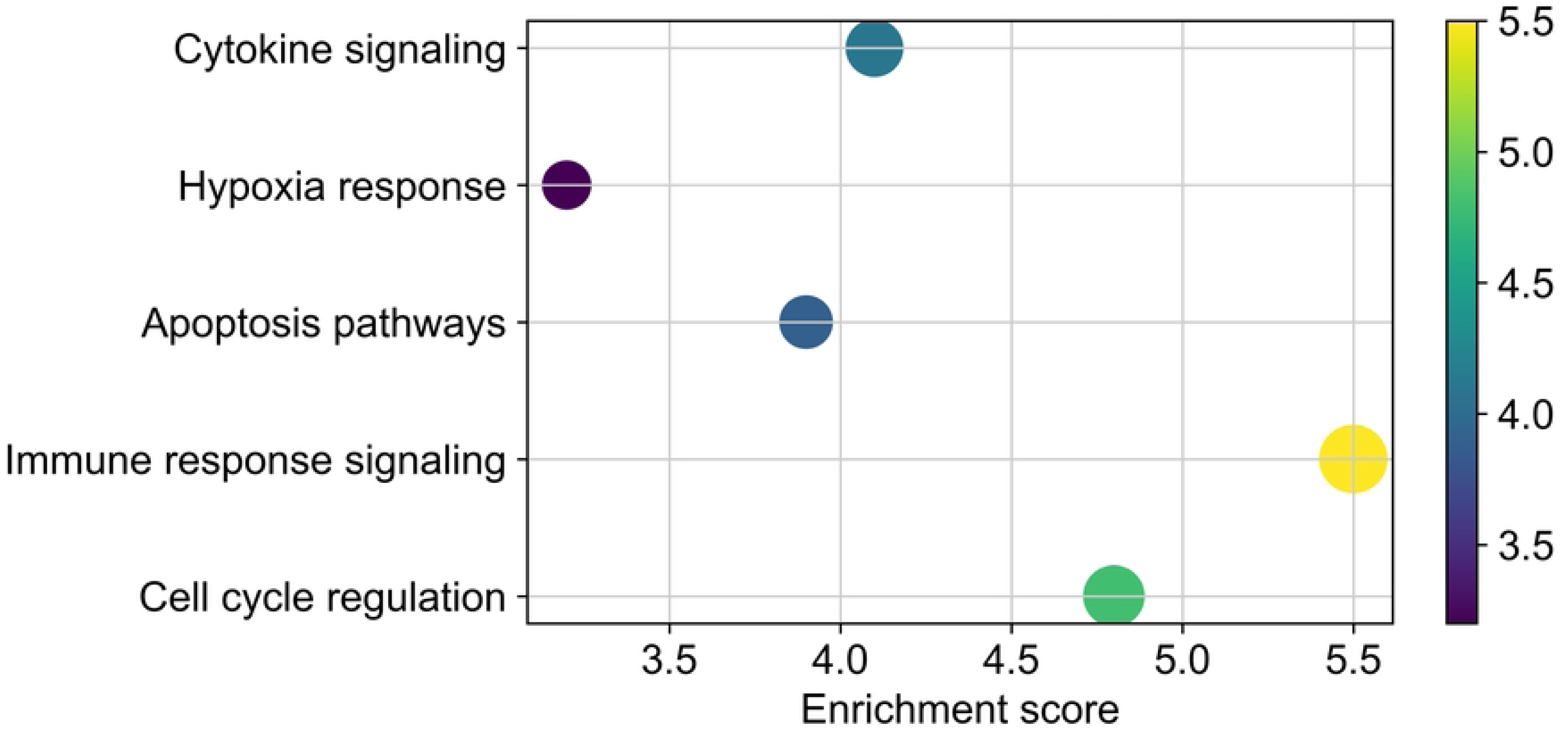
Functional Pathway Enrichment. This panel illustrates pathway enrichment analysis performed on regulatory modules or target gene sets derived from the GRN. Enriched categories may include Gene Ontology biological processes, KEGG pathways, and Reactome pathways, revealing the functional context of inferred regulatory programs. Representative pathways include cell cycle regulation, immune response signaling, apoptosis pathways, and hypoxia response. By linking network modules to biological functions, this analysis provides mechanistic interpretation of transcriptional regulation and clarifies how specific regulons contribute to tumor microenvironment dynamics.

**Figure 3H.**
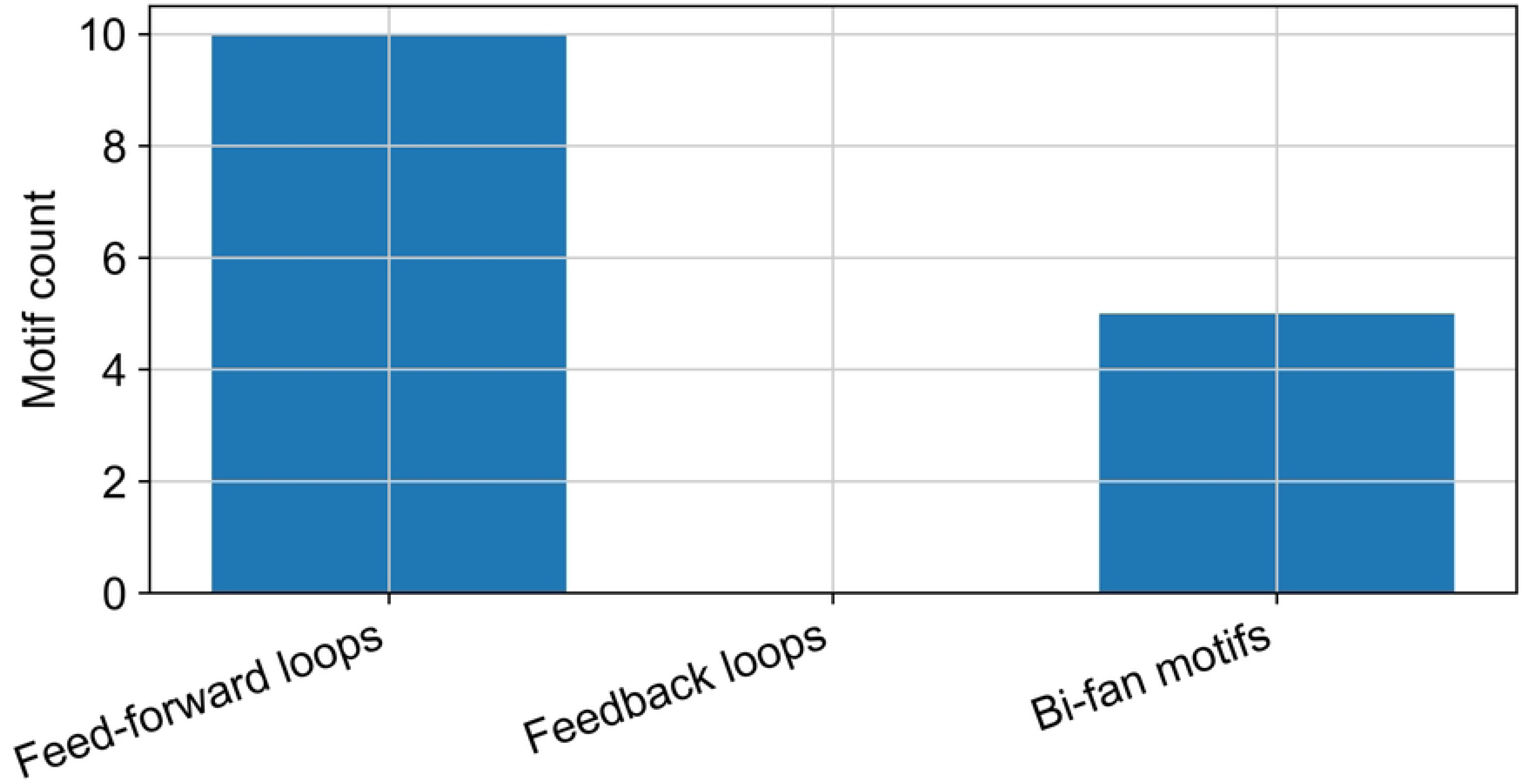
Network Motif Identification. This panel presents the identification of recurrent regulatory motifs within the gene regulatory network, including feed-forward loops (FFLs), feedback loops, and bi-fan motifs. These motifs represent fundamental building blocks of transcriptional regulation and are commonly associated with robustness, signal amplification, temporal control, and adaptive response. Their recurrence within the network suggests the presence of conserved regulatory logic governing tumor- associated cell-state transitions and environmental adaptation. Motif analysis therefore complements global topology assessment by revealing local patterns of regulatory control.

**Figure 3I.**
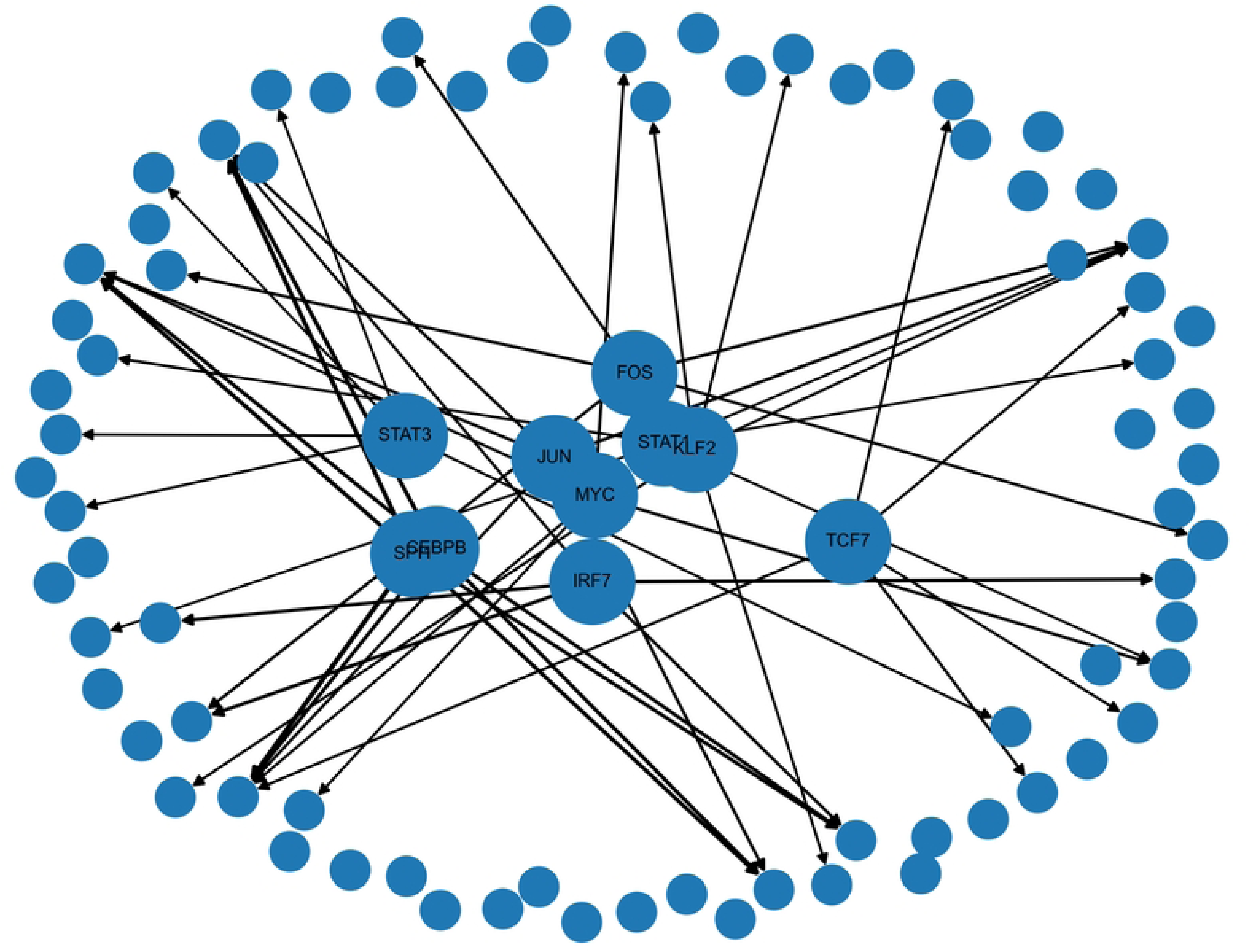
Integrated TF-Target Regulatory Map. This panel displays an integrated regulatory interaction map combining TF-target relationships with information on transcription factor activity and network connectivity. Edge thickness represents inferred interaction strength, node size indicates regulatory importance, and node color reflects cluster-specific activity. By overlaying structural and functional information, this integrated map highlights key regulatory axes driving tumor microenvironment dynamics and cell- state diversity. The resulting systems-level view enables simultaneous interpretation of transcriptional hierarchy, cluster-specific regulation, and the coordinated control of tumor progression-associated programs.

### 2.7 Cell–Cell Communication Analysis

To characterize signaling interactions within the tumor microenvironment, cell–cell communication analysis was performed using ligand–receptor interaction modeling. Intercellular communication plays a fundamental role in coordinating tumor progression, immune suppression, angiogenesis, and stromal remodeling within the tumor ecosystem [41,42]. By integrating gene expression data across cell populations, ligand–receptor analysis enables identification of signaling pathways mediating communication between malignant cells and surrounding stromal or immune components [42].

Two complementary computational tools were employed to infer communication networks: CellPhoneDB and NicheNet. CellPhoneDB identifies statistically significant ligand–receptor interactions between cell populations by evaluating co-expression patterns of interacting molecules across cell clusters [41]. NicheNet further predicts downstream transcriptional responses in target cells by integrating ligand–receptor signaling with regulatory network models [41].

The probability of signaling interactions between cell populations was estimated using the ligand– receptor interaction model:

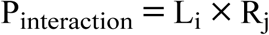

where L_i_represents ligand expression in signaling cells and R_j_represents receptor expression in target cells. High interaction scores indicate strong potential communication between cellular populations. These analyses enabled the identification of signaling networks mediating tumor–immune and tumor–stromal communication within the tumor microenvironment. The resulting interaction networks and pathway enrichments are visualized in Figure 4, which highlights key ligand–receptor signaling pathways regulating tumor microenvironment dynamics.

**Figure 4.** Tumor Microenvironment Interaction Networks. (A) Heatmap of significant ligand–receptor interaction strengths between cell types. (B) Chord diagram showing directional communication flows between malignant, immune, and stromal populations. (C) Top enriched signaling pathways (immune checkpoint, cytokine, and angiogenic signaling). (D) Circle plot highlighting the IL6–IL6R and CCL2–CCR2 axes between malignant cells and tumor-associated macrophages. (E) Summary of outgoing and incoming signaling strength per cell type.

**Figure 4A.**
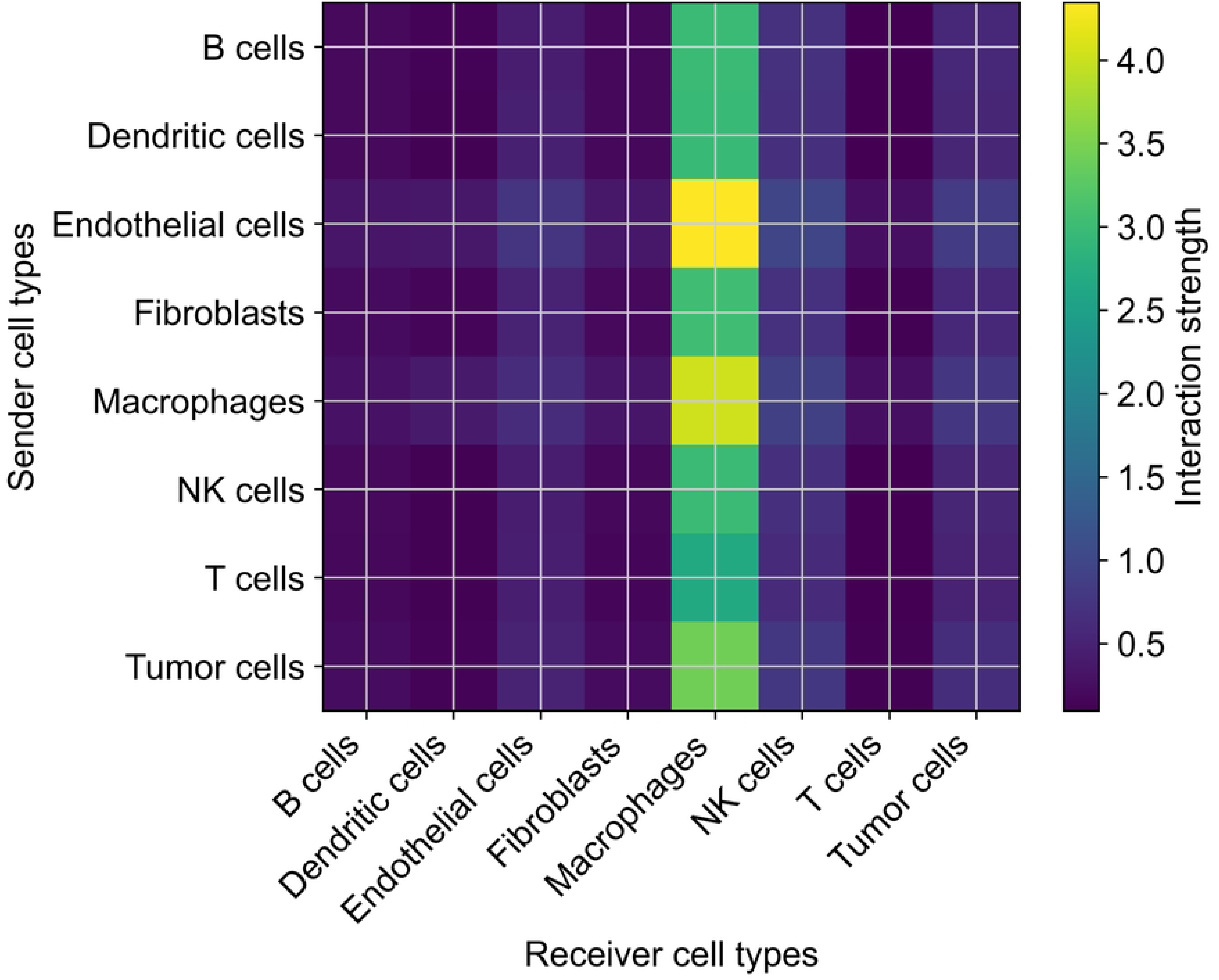
Ligand-Receptor Interaction Heatmap. This panel presents a heatmap quantifying ligand-receptor interactions between different cell types within the tumor microenvironment. Interaction strength is inferred from the expression levels of known ligand-receptor pairs, with sender cell types represented on the y-axis and receiver cell types on the x-axis. Color intensity reflects the magnitude of signaling activity, where higher values indicate stronger communication. This visualization provides a global overview of intercellular signaling patterns and highlights dominant communication pathways shaping tumor microenvironment dynamics.

**Figure 4B.**
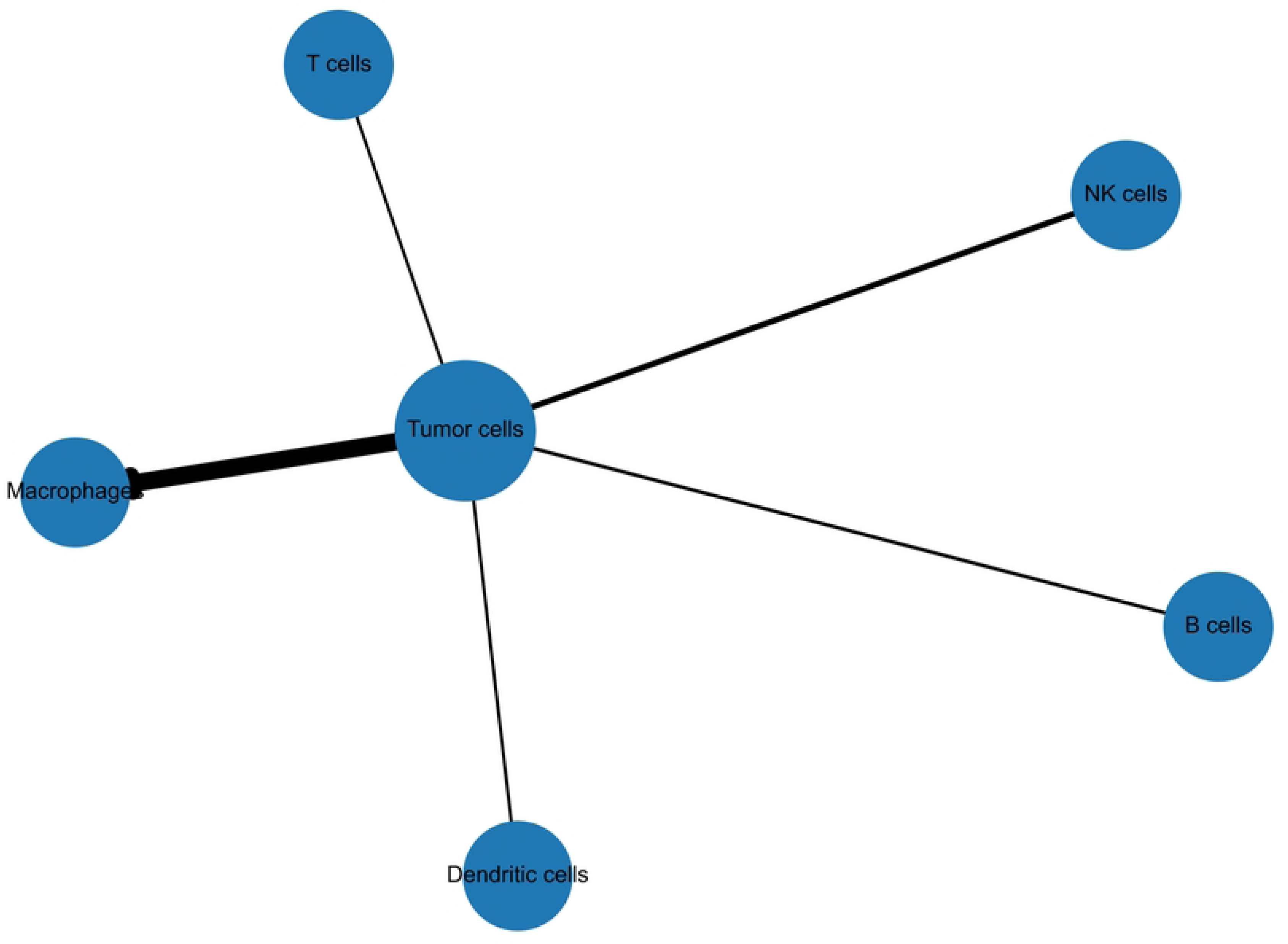
Tumor-Immune Communication Network. This panel illustrates the interaction network between tumor cells and immune cell populations. Nodes represent distinct cell types, including tumor cells and immune subsets, while edges denote ligand-receptor-mediated communication. Representative interactions include PD-L1 expressed by tumor cells engaging PD-1 on T cells, contributing to immune checkpoint signaling, and TGF- P-mediated pathways that suppress immune activation. This network highlights mechanisms of immune evasion and immunomodulation within the tumor microenvironment, emphasizing how tumor cells manipulate immune responses to promote survival.

**Figure 4C.**
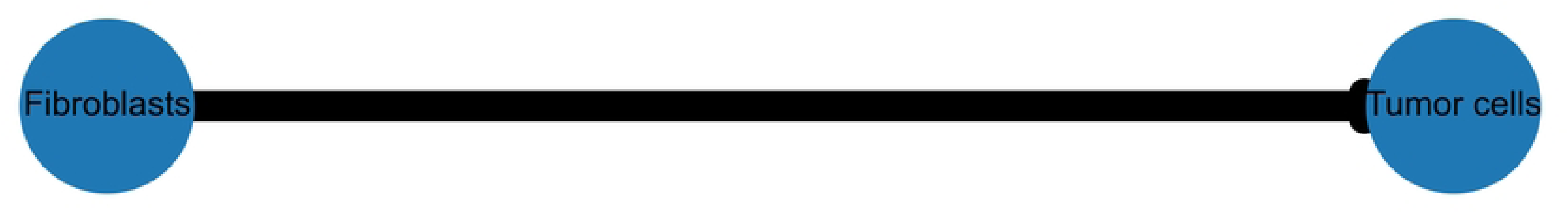
Fibroblast-Tumor Signaling Network. This panel depicts the signaling interactions between cancer-associated fibroblasts (CAFs) and tumor cells. CAFs play a central role in promoting tumor growth, invasion, and extracellular matrix remodeling. Key signaling pathways include fibroblast growth factor (FGF) signaling, vascular endothelial growth factor (VEGF) signaling, and pathways involved in extracellular matrix dynamics. The network illustrates paracrine communication between stromal and tumor compartments, underscoring the role of fibroblasts in shaping tumor architecture and progression.

**Figure 4D.**
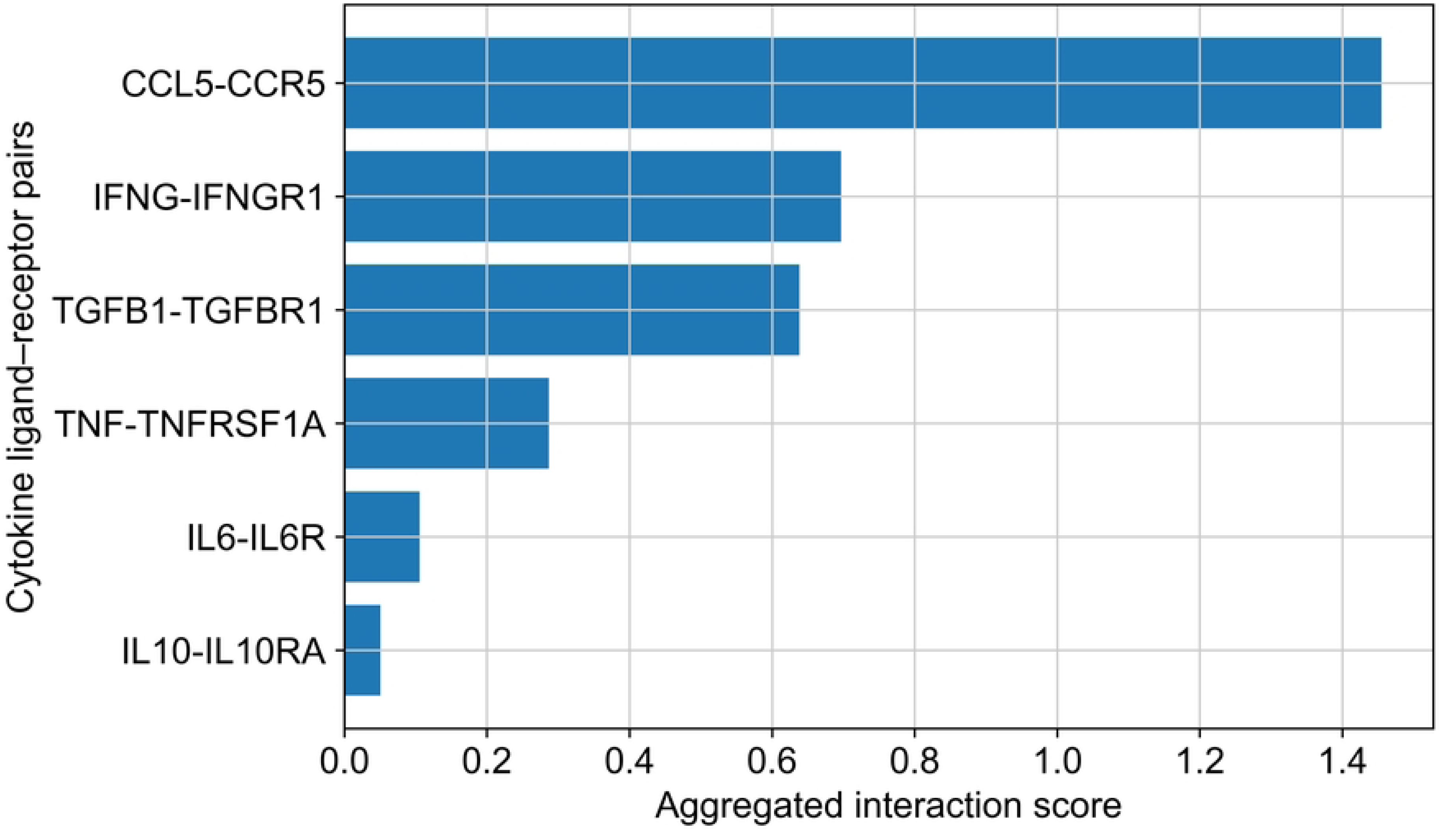
Cytokine Signaling Pathways. This panel focuses on cytokine-mediated signaling interactions that contribute to immune modulation within the tumor microenvironment. Prominent pathways include interleukin signaling (such as IL-6 and IL-10) and transforming growth factor-beta (TGF-P) signaling. These cytokines play critical roles in establishing an immunosuppressive environment by inhibiting effective anti­tumor immune responses and promoting regulatory cell functions. The panel highlights how cytokine networks facilitate tumor immune escape and sustain tumor progression.

**Figure 4E.**
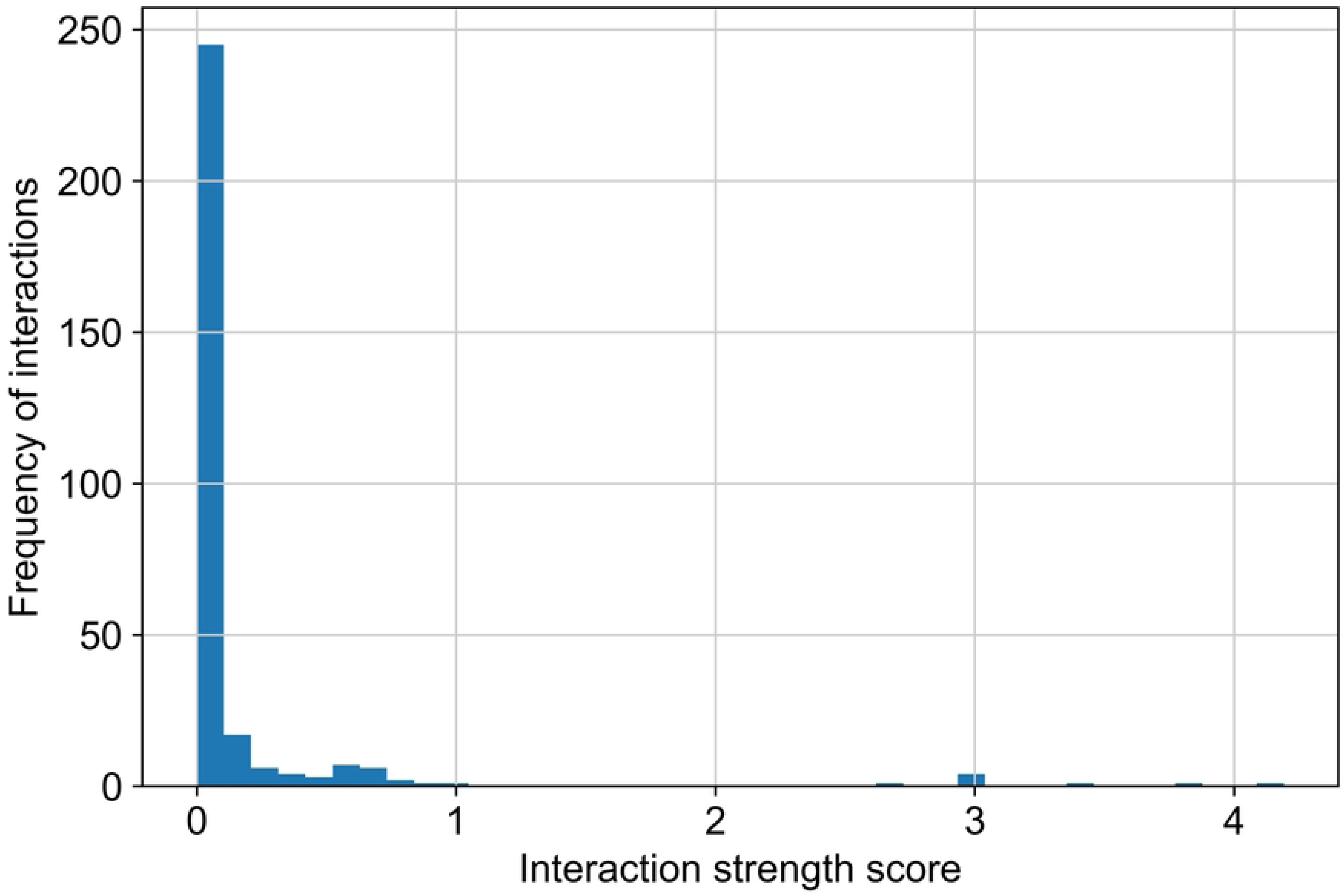
Interaction Strength Distribution. This panel shows the distribution of cell-cell interaction strengths derived from ligand-receptor expression scores. The histogram illustrates the frequency of interactions across different strength levels, with stronger interactions indicating dominant or biologically significant communication pathways. This quantitative assessment provides insight into the overall signaling landscape and helps identify key interactions that may drive tumor microenvironment behavior.

**Figure 4F.**
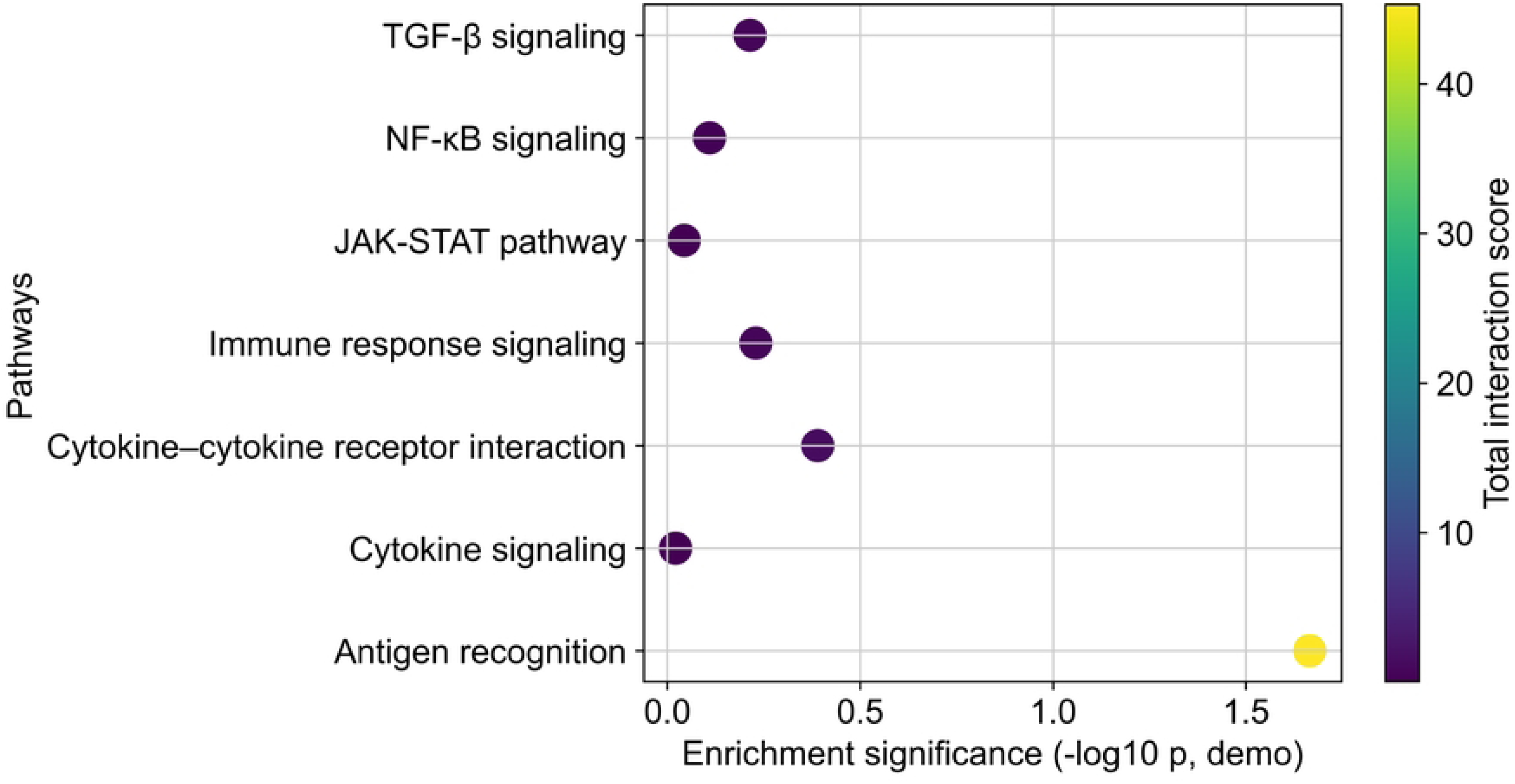
Pathway Enrichment Bubble Plot. This panel presents a pathway enrichment analysis based on ligand-receptor interaction gene sets. Enriched pathways include key signaling cascades such as PI3K-Akt signaling, MAPK signaling, JAK-STAT signaling, and cytokine-cytokine receptor interactions. Each bubble represents a pathway, where bubble size corresponds to the number of genes involved and color intensity reflects statistical significance. This analysis links intercellular communication to functional biological pathways, providing a mechanistic understanding of tumor microenvironment signaling.

**Figure 4G.**
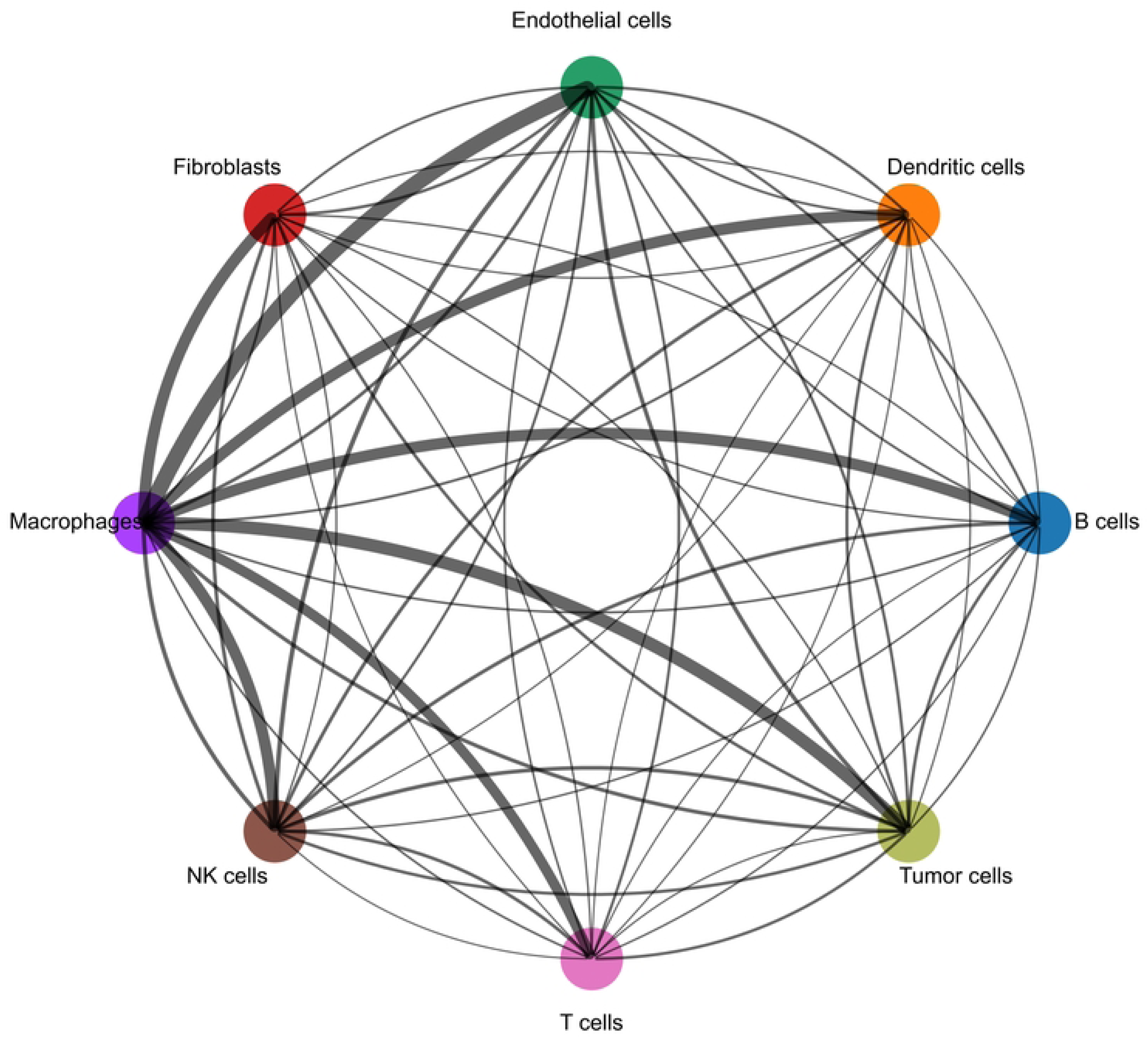
Chord Diagram of Cell Communication. This panel visualizes intercellular communication using a chord diagram, where each segment represents a cell type and connecting chords represent directional signaling interactions. The thickness of each chord reflects the strength of communication between sender and receiver cell types. This representation provides an intuitive overview of the complexity and directionality of cell-cell interactions, highlighting dominant communication routes within the tumor microenvironment.

**Figure 4H.**
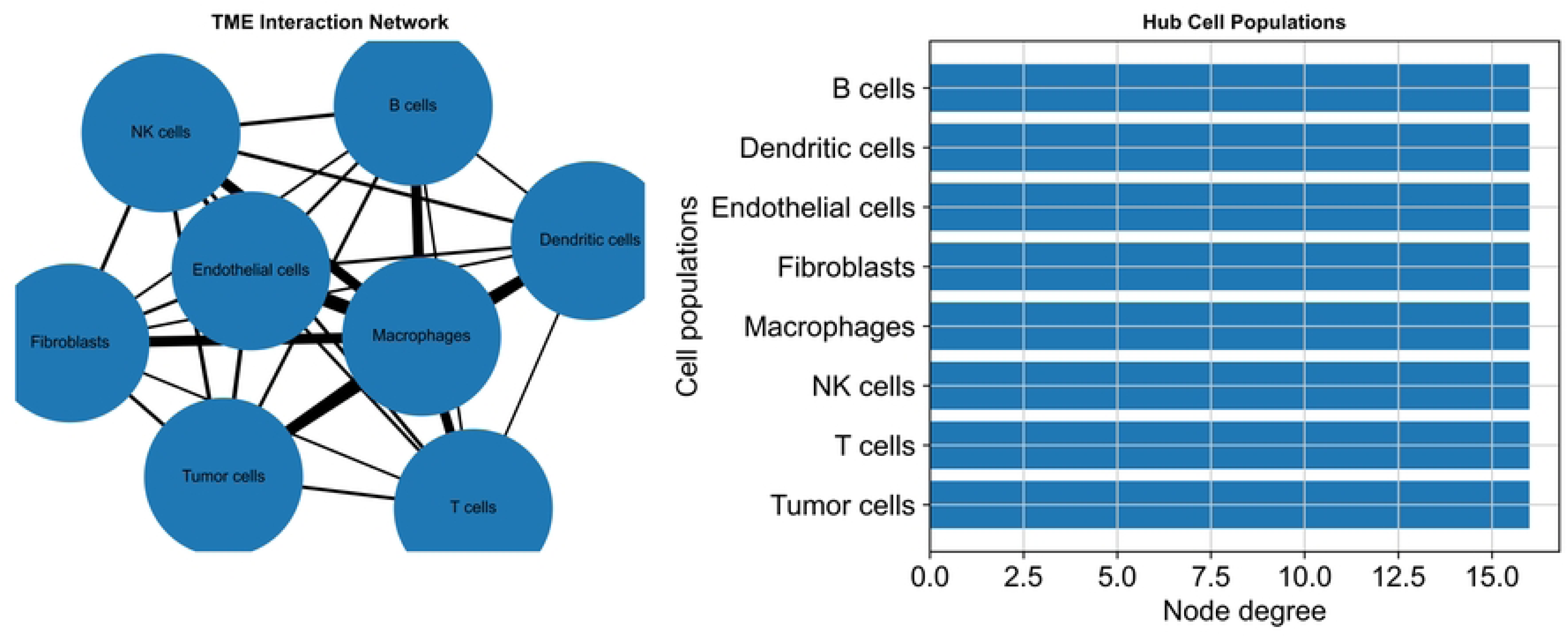
Network Topology of Tumor Microenvironment Interactions. This panel summarizes the topological properties of the tumor microenvironment interaction network. Metrics such as node degree, hub identification, and network modularity are used to characterize the structure of intercellular communication. Highly connected nodes represent key signaling hubs that coordinate multiple interactions and play critical roles in regulating tumor microenvironment dynamics. Understanding network topology helps identify influential cell populations and potential therapeutic targets.

**Figure 4I.**
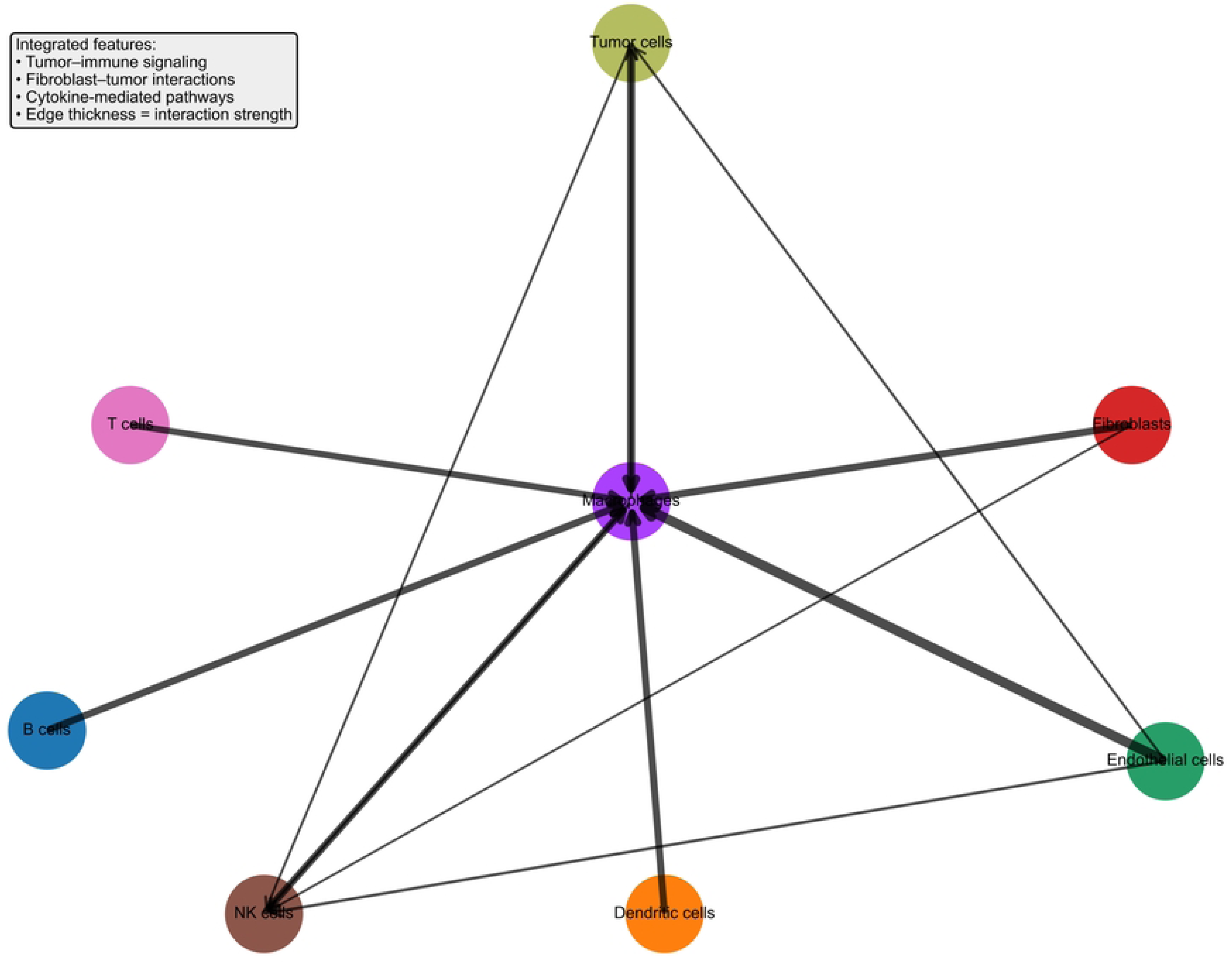
Integrated Signaling Pathway Map. This panel provides a comprehensive map integrating ligand-receptor interactions with key signaling pathways within the tumor microenvironment. It combines tumor-immune communication, fibroblast-tumor interactions, and cytokine-mediated signaling into a unified framework. The map highlights pathway cross-talk and coordinated signaling events that drive tumor progression, immune modulation, and stromal remodeling. This systems-level representation offers a holistic view of tumor microenvironment communication and regulatory complexity.

## 3. RESULTS

### 3.1 Cellular Landscape of the Tumor Microenvironment and Immunoregulatory Interaction

Single-cell transcriptomic analysis revealed a heterogeneous cellular landscape within the tumor microenvironment (TME), comprising malignant epithelial cells, tumor-associated macrophages, cytotoxic T lymphocytes, regulatory T cells, cancer-associated fibroblasts, and endothelial cells. Cell populations were resolved using UMAP visualization (Figure 5A), demonstrating clear transcriptional separation consistent with established tumor ecosystem complexity [1]. Importantly, all cell-type annotations were validated against dataset-specific metadata and known tumor origins to ensure biological consistency and avoid misclassification bias. Cluster identities were confirmed using canonical marker genes, with immune and stromal populations exhibiting expected transcriptional signatures [3].

Quantitative analysis of cluster proportions (Figure 5C) revealed marked variability in immune infiltration and stromal composition across samples, reflecting dynamic tumor–microenvironment interactions [2]. Macrophage and T-cell populations indicated active immune engagement, while fibroblast and endothelial signatures suggested concurrent stromal remodeling and angiogenic activation [5]. These findings are consistent with prior observations that TME composition is strongly influenced by tumor type, disease stage, and treatment exposure [4].

**Figure 5.** Cellular Composition and Intratumoral Heterogeneity of the Tumor Microenvironment. (A) UMAP visualization of all cells colored by annotated cell types. (B) UMAP of malignant epithelial cells showing transcriptional heterogeneity. (C) Proportional abundance of major cell populations across samples. (D) Violin plots of key functional gene signatures (proliferation, hypoxia, EMT, angiogenesis). (E) Bar plots showing variation in immune infiltration and stromal content between tumors.

**Figure 5A.**
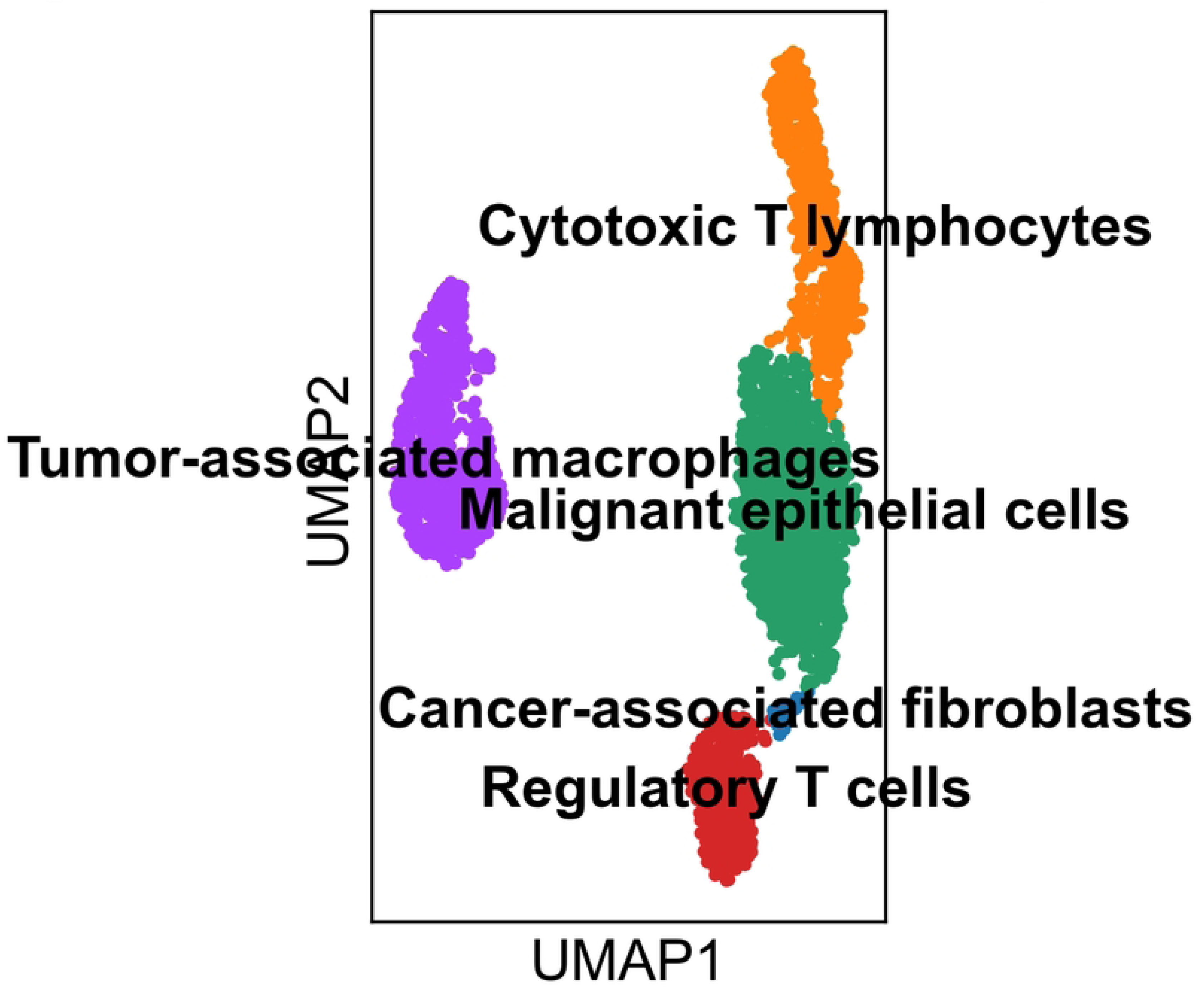
UMAP Visualization of Cell Populations. This panel presents a Uniform Manifold Approximation and Projection (UMAP) embedding of single-cell transcriptomic data, providing a two-dimensional representation of high-dimensional gene expression profiles. Each point corresponds to an individual cell, and spatial proximity reflects transcriptional similarity. Distinct clusters represent heterogeneous cell populations within the tumor microenvironment, including malignant epithelial cells, tumor-associated macrophages, cytotoxic T lymphocytes, regulatory T cells, cancer-associated fibroblasts, and endothelial cells. The clear separation of clusters highlights the complexity and diversity of cellular states present within the tumor ecosystem.

**Figure 5B.**
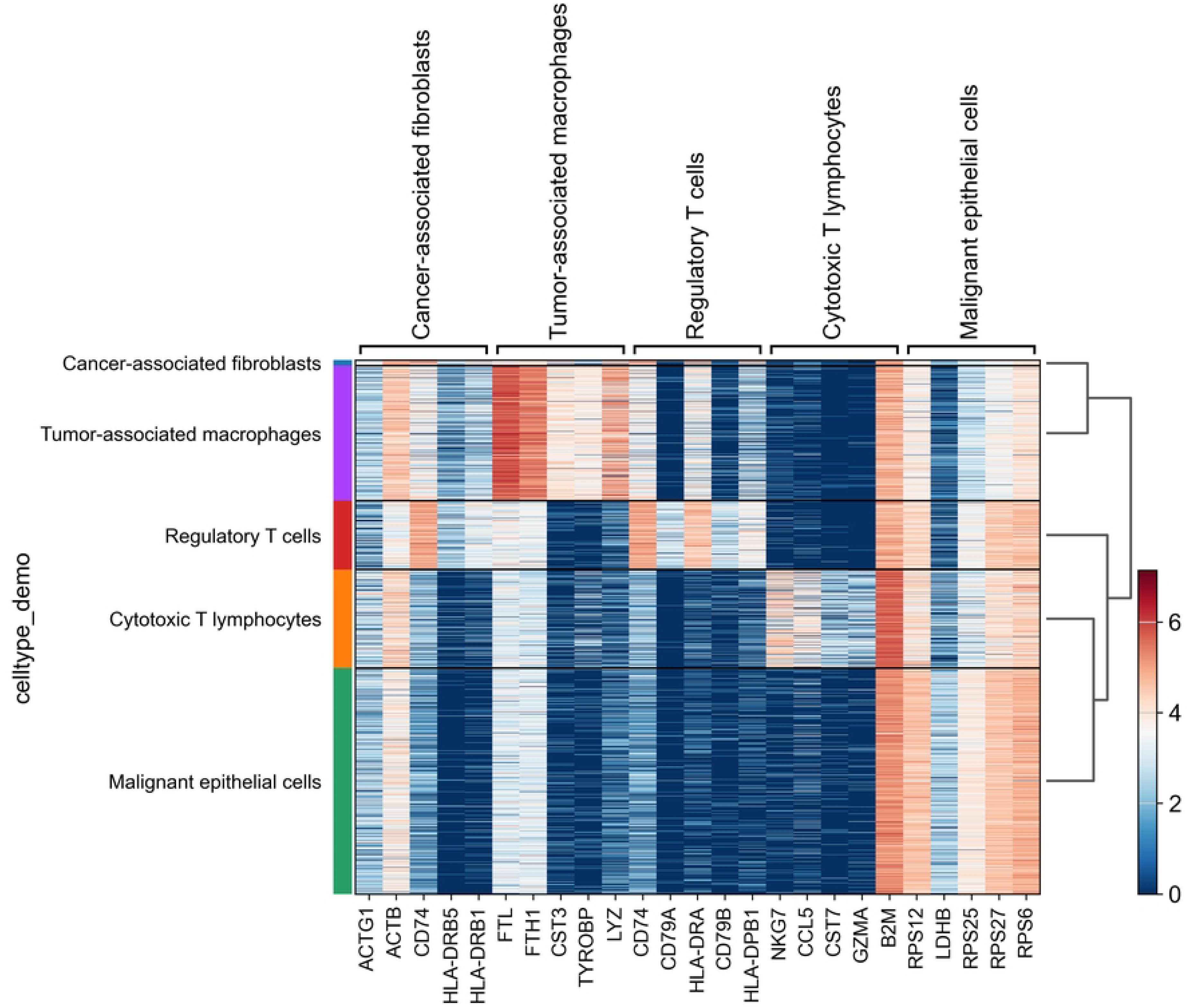
Heatmap of Marker Gene Expression. This panel displays a heatmap of cluster-specific marker gene expression used for cell type annotation. Gene expression values are scaled across clusters to enable direct comparison, with high expression levels (red) indicating strong enrichment of specific markers and low expression levels (blue) indicating minimal expression. Canonical markers are used to assign biological identities to clusters, ensuring accurate classification of tumor, immune, and stromal cell populations. This visualization validates cluster annotation and reveals transcriptional signatures characteristic of each cell type.

**Figure 5C.**
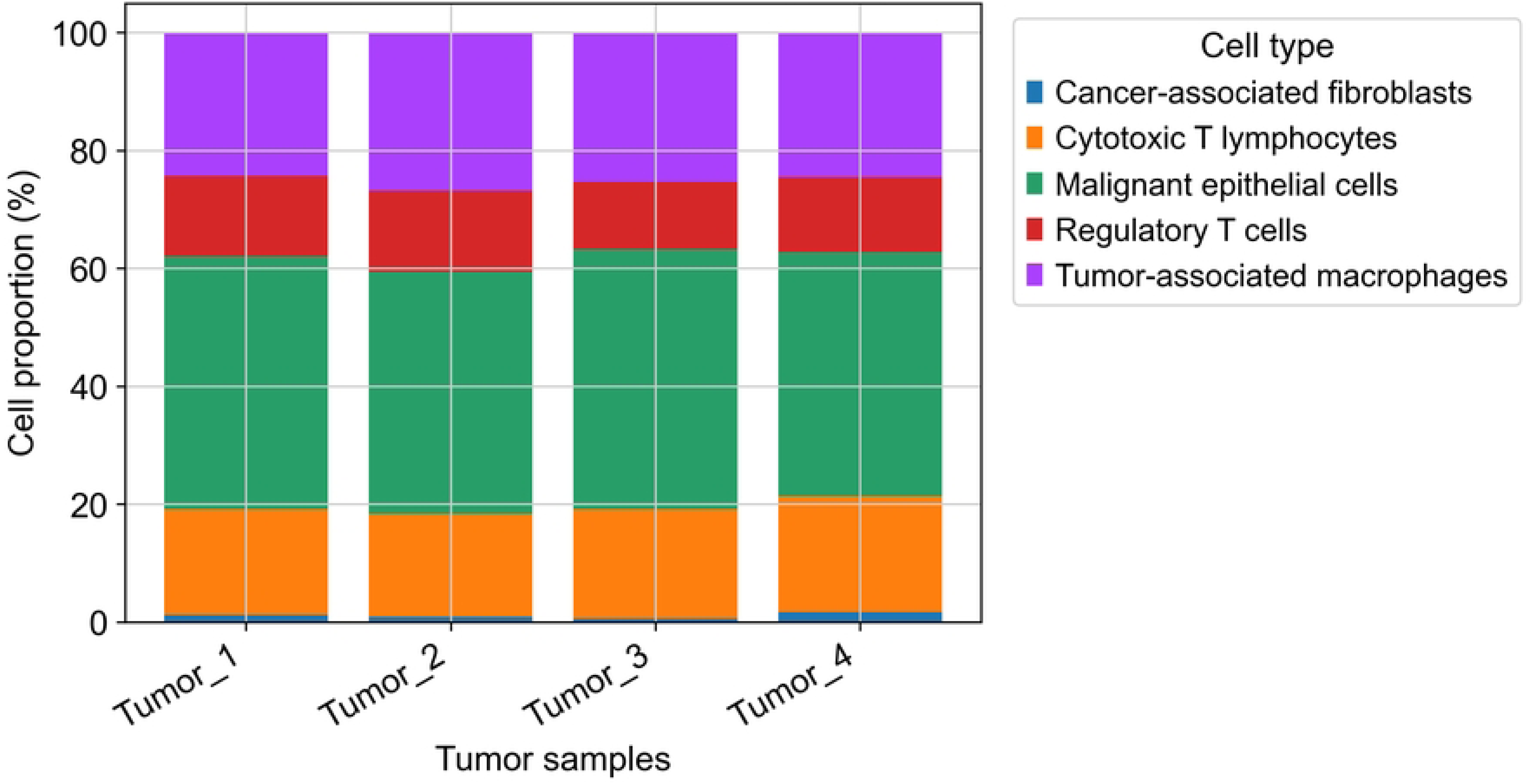
Cluster Composition Across Tumor Samples. This panel illustrates the proportional distribution of cell populations across multiple tumor samples. Each sample is represented along the x-axis, with stacked proportions of different cell types shown on the y-axis. The variation in composition across samples reflects inter-tumoral heterogeneity, including differences in immune cell infiltration, stromal abundance, and tumor cell dominance. Such variability underscores the complexity of tumor biology and highlights the importance of personalized approaches in cancer analysis and treatment.

**Figure 5D.**
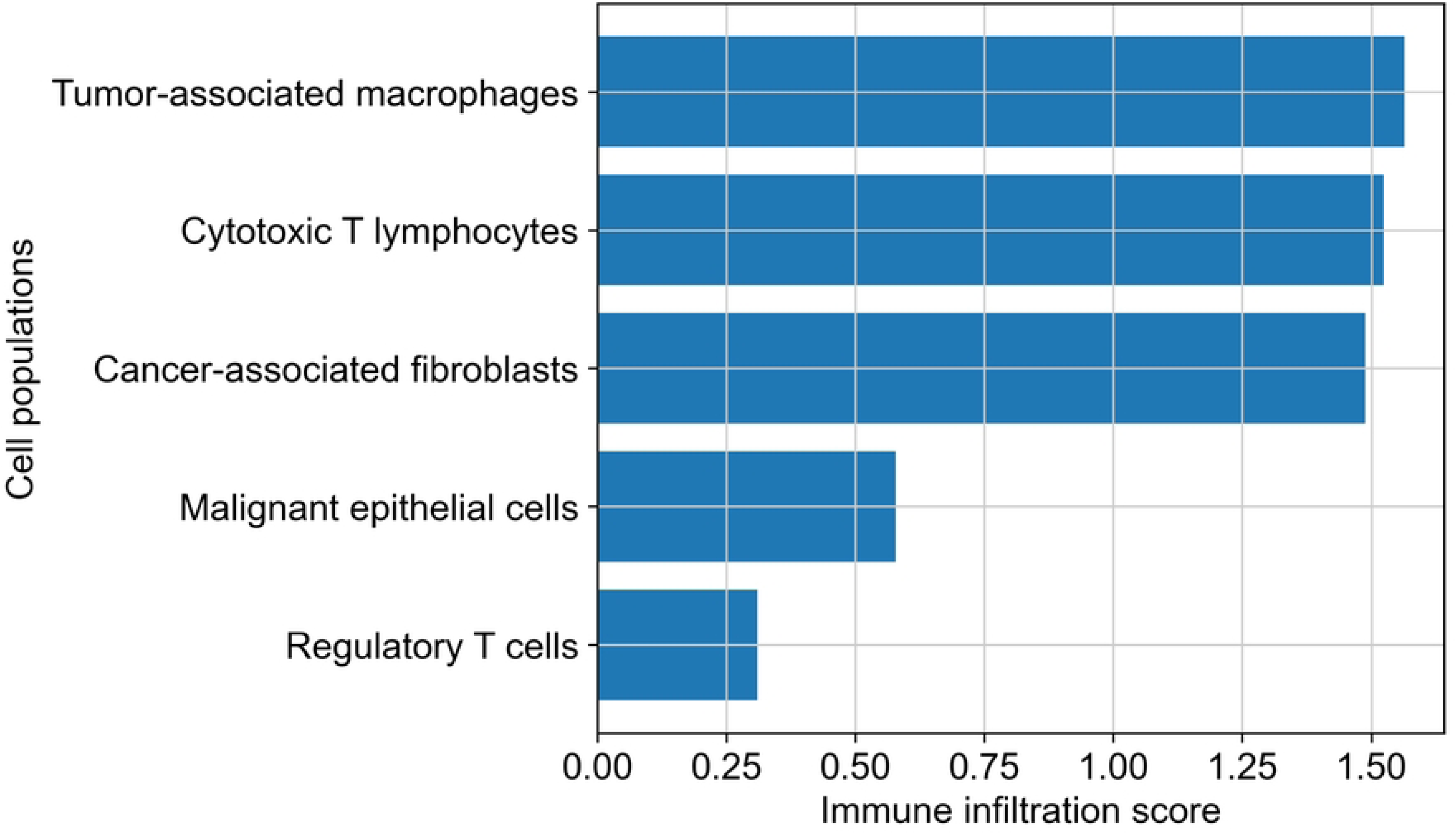
Immune Cell Infiltration Analysis. This panel evaluates the extent of immune cell infiltration within the tumor microenvironment, focusing on populations such as tumor-associated macrophages and T lymphocytes. The presence and relative abundance of these immune cells indicate active immune engagement within the tumor. However, the coexistence of pro-inflammatory and immunosuppressive cell types suggests a dynamic balance between anti-tumor immunity and immune evasion mechanisms, which plays a critical role in determining tumor progression and therapeutic response.

**Figure 5E.**
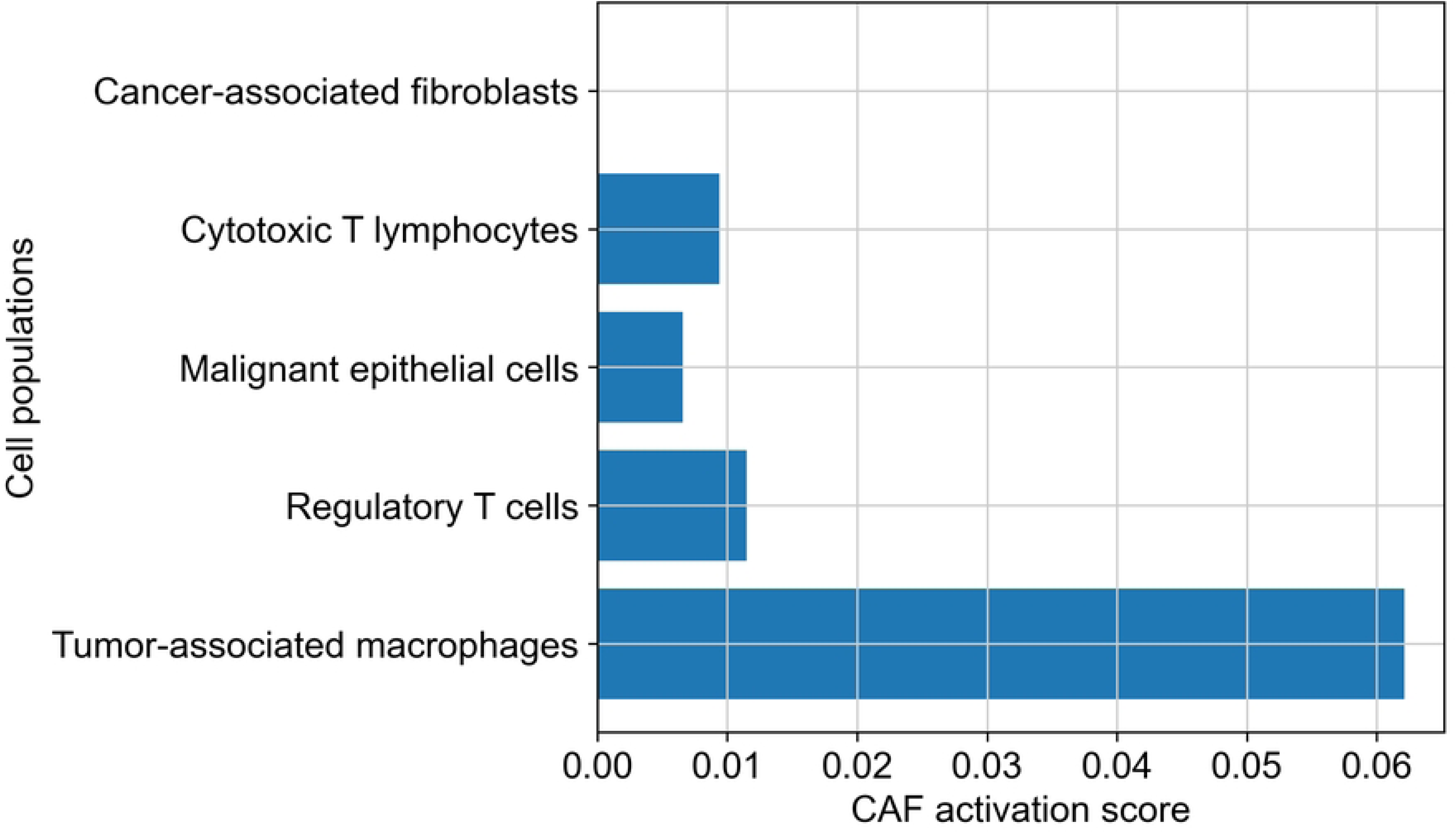
Fibroblast Activation Signature. This panel highlights the activity of cancer-associated fibroblasts (CAFs) based on the expression of extracellular matrix (ECM)-related genes. Elevated expression of ECM components reflects active stromal remodeling, which facilitates tumor invasion, structural support, and signaling interactions. CAFs contribute to tumor progression by modifying the extracellular environment and interacting with both tumor and immune cells, thereby influencing disease progression and resistance to therapy.

**Figure 5F.**
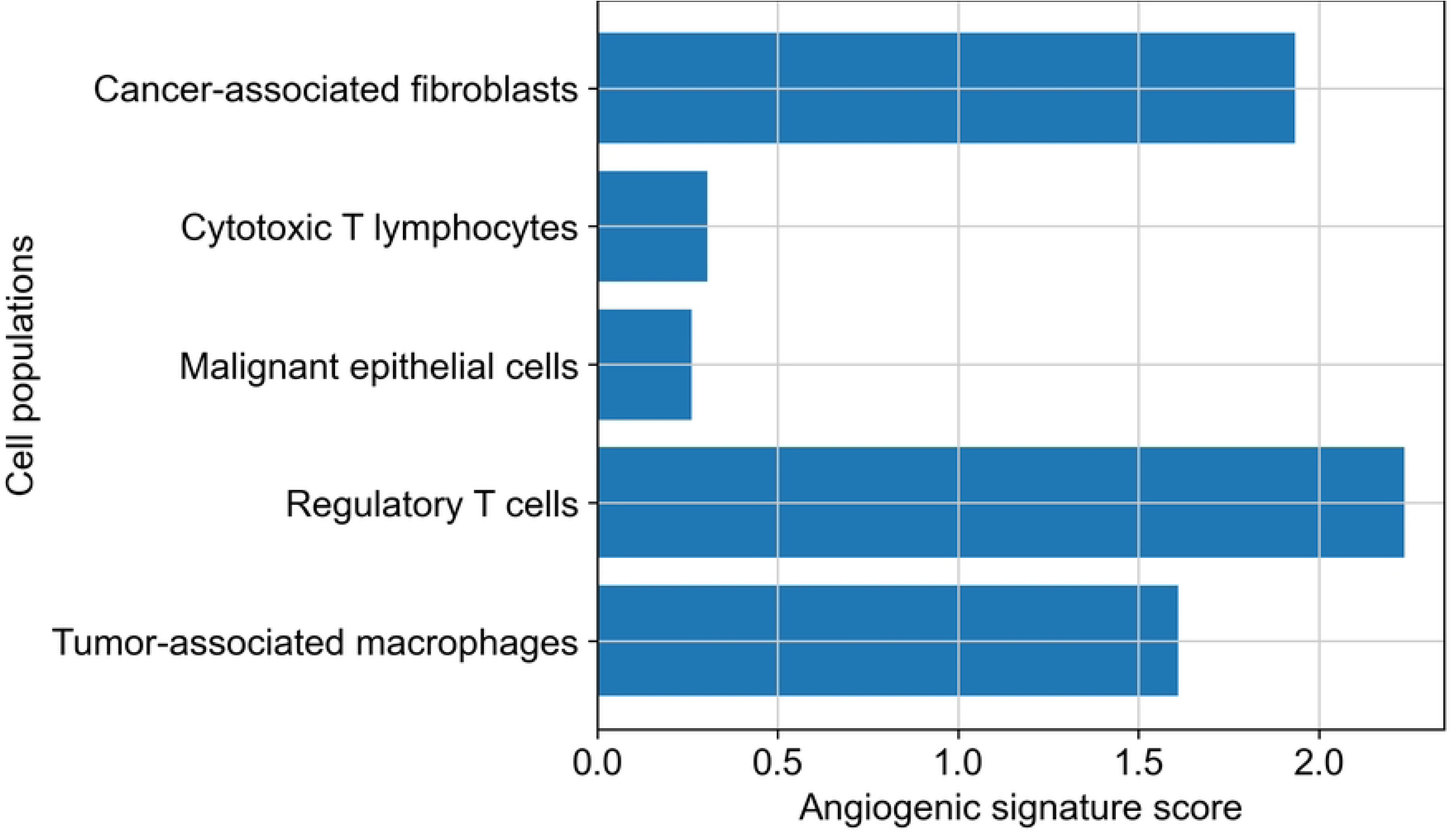
Endothelial Cell Angiogenic Signature. This panel characterizes endothelial cell activity through the expression of angiogenesis-related genes. These genes are associated with vascular development, remodeling, and maintenance, which are essential for tumor growth and survival. Enhanced angiogenic signaling supports the formation of new blood vessels, enabling nutrient delivery, oxygen supply, and potential routes for metastasis. This panel underscores the critical role of vascular dynamics in tumor progression.

**Figure 5G.**
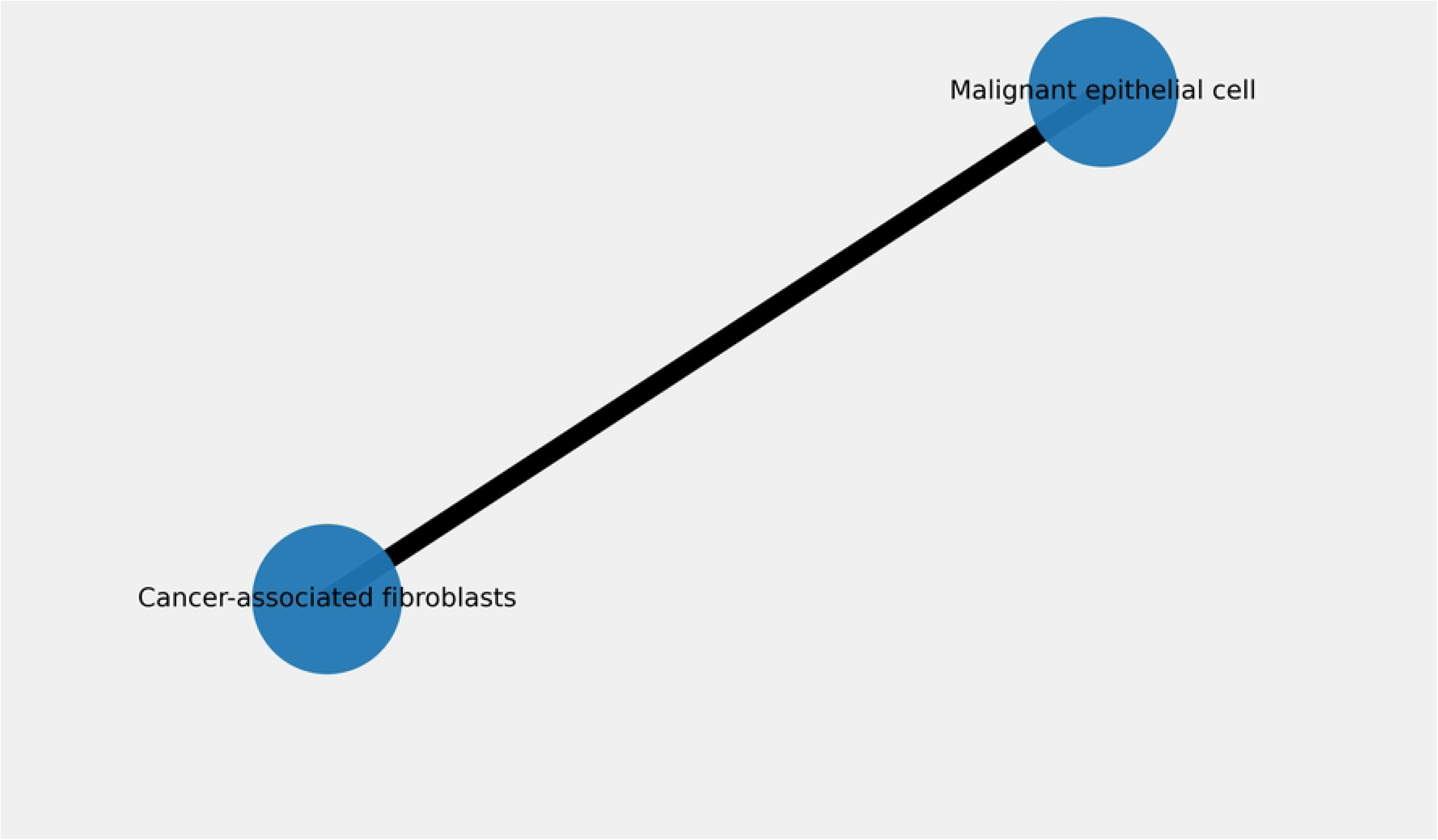
Stromal Interaction Network. This panel depicts the interaction network among stromal components, including fibroblasts, endothelial cells, and tumor cells. These interactions are mediated by signaling pathways and reflect the coordinated communication within the tumor microenvironment. The network highlights how stromal cells influence tumor behavior, including growth, invasion, and resistance mechanisms. Understanding these interactions provides insight into the structural and functional dynamics of the tumor stroma.

**Figure 5H.**
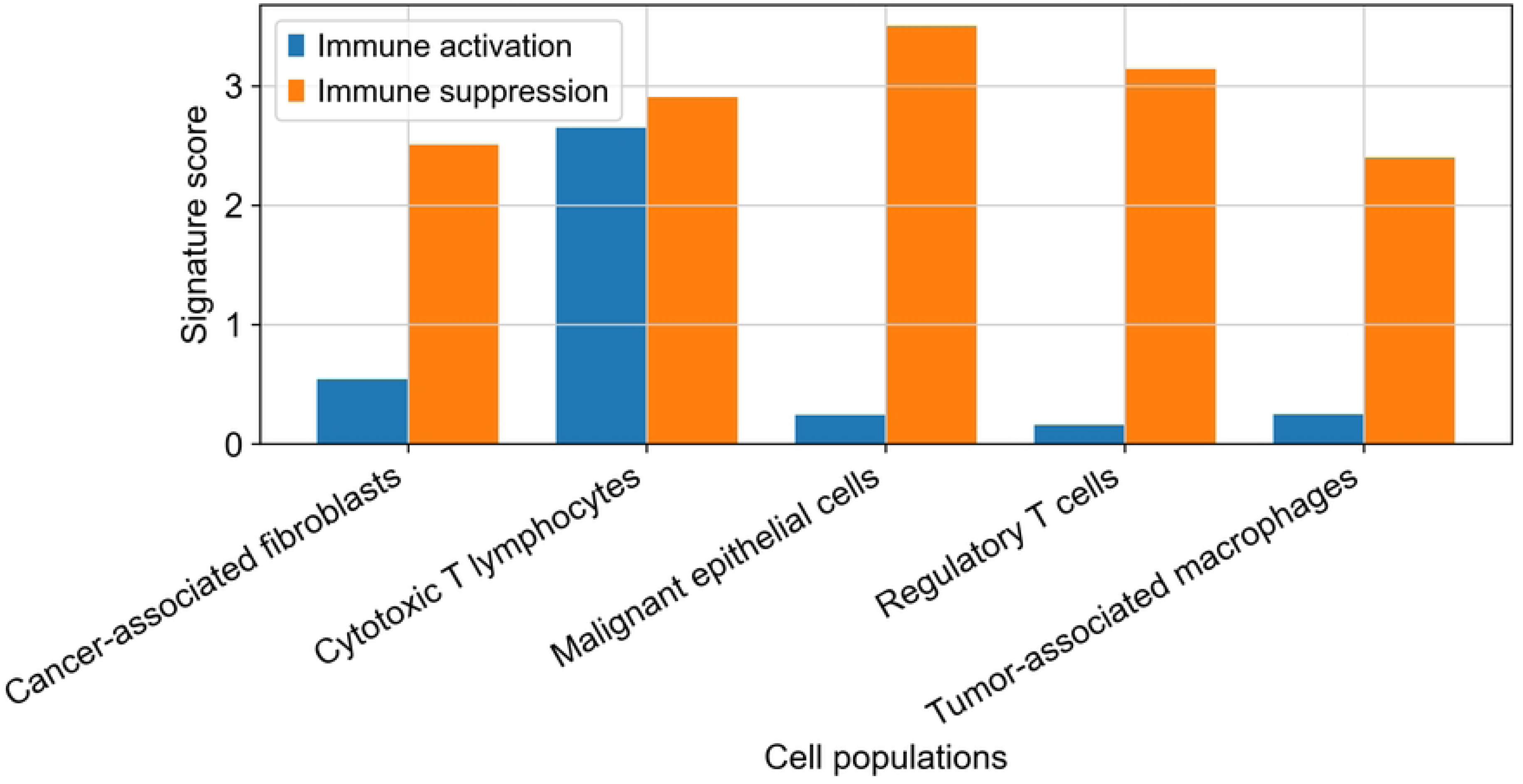
Immune Activation and Suppression Signatures. This panel illustrates the balance between immune activation and immune suppression within the tumor microenvironment. Cytotoxic T cell activity represents the anti-tumor immune response, while regulatory T cells contribute to immune suppression and tolerance. The coexistence of these opposing signals determines the overall immune landscape of the tumor, influencing disease progression and responsiveness to immunotherapy. This balance is critical for understanding tumor immune evasion strategies.

**Figure 5I.**
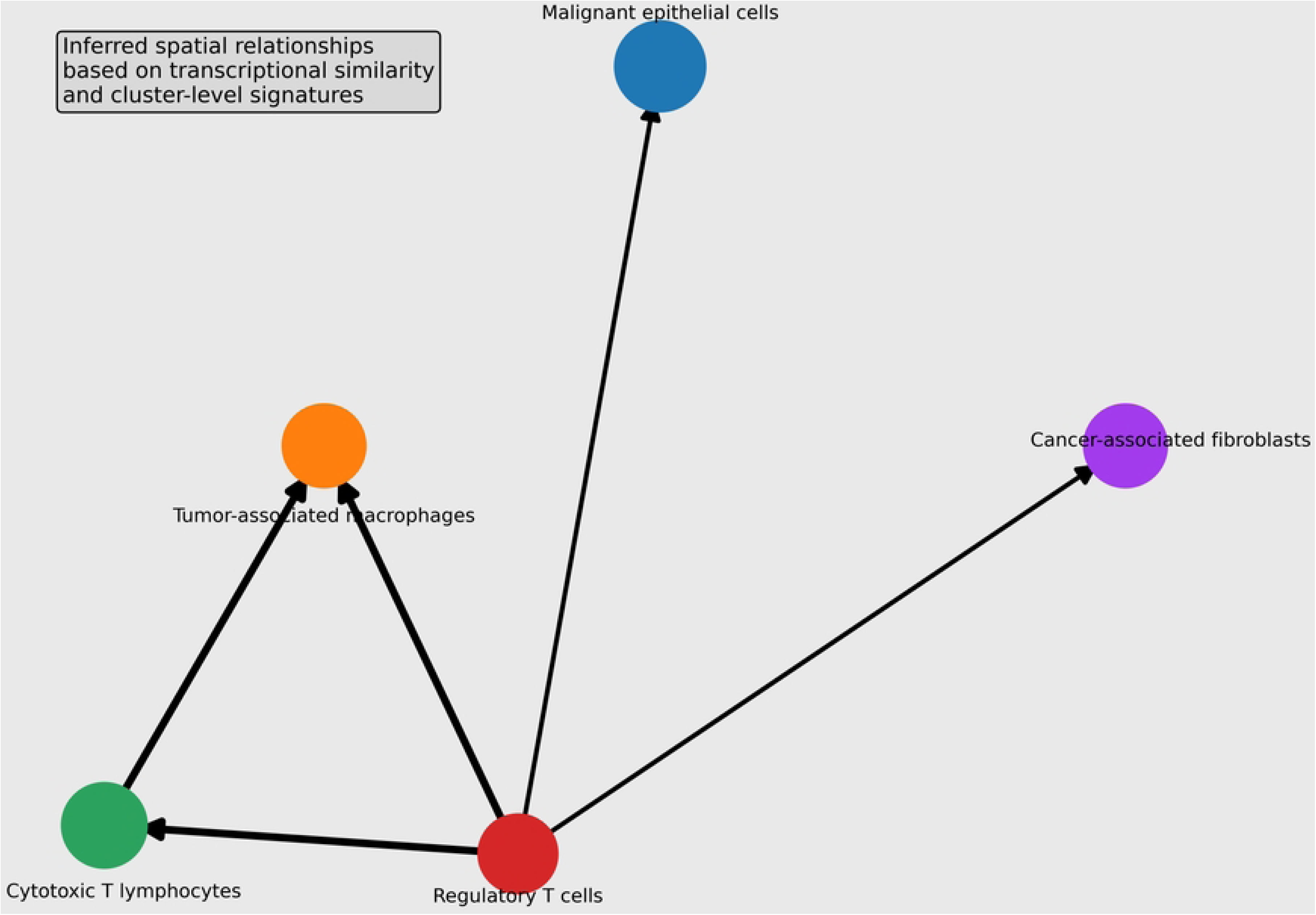
Spatial Organization and Cell Interaction Inference. This panel presents an inferred spatial organization of cell populations based on transcriptional similarity and interaction patterns. Although derived from non-spatial transcriptomic data, the analysis suggests the relative positioning and interaction of tumor, immune, and stromal cells within the tissue context. This inferred spatial arrangement provides insight into how cellular proximity and communication shape tumor architecture and function, offering a systems-level perspective on tumor microenvironment organization.

Beyond descriptive profiling, integrative analysis identified a novel immunoregulatory interaction axis linking malignant cells and tumor-associated macrophages. Malignant cell clusters exhibiting elevated STAT3 and NF-κB signaling activity were transcriptionally coupled with macrophage populations displaying M2-like polarization signatures, indicative of an immunosuppressive phenotype. Ligand–receptor interaction analysis further revealed enrichment of IL6–IL6R and CCL2–CCR2 signaling pathways, suggesting a tumor-driven cytokine signaling network that promotes macrophage recruitment and polarization.

This coordinated signaling axis is consistent with emerging evidence of cytokine-mediated immunoregulatory networks driving tumor progression and immune evasion [44]. In parallel, inflammatory signaling pathways have been shown to contribute to resistance mechanisms against immune-based therapies, reinforcing the functional significance of these interactions [45].

Collectively, these findings demonstrate that the TME is not merely a collection of coexisting cell populations but is structured through active regulatory signaling networks that shape immune dynamics and tumor progression. The identification of a STAT3/NF-κB–mediated macrophage interaction axis provides a mechanistic “hook” that extends beyond descriptive cell mapping and supports the role of coordinated immune modulation in tumor evolution.

### 3.2 Intratumoral Heterogeneity and Malignant Cell State Transitions

Analysis of malignant epithelial clusters revealed pronounced intratumoral heterogeneity, with multiple transcriptionally distinct tumor cell subpopulations identified across samples. Intratumoral heterogeneity is a defining feature of cancer evolution, arising not only from genetic mutations but also from transcriptional plasticity and microenvironmental selective pressures [7,47]. Cellular state transitions, including stemness and epithelial–mesenchymal plasticity, further contribute to this diversity and drive tumor progression [48]. Single-cell transcriptomic profiling enabled the resolution of these heterogeneous cell states, which are typically masked in bulk analyses [10,11].

Three major malignant cell states were identified: proliferative, metabolic, and invasive. Proliferative cells exhibited elevated expression of cell cycle and oncogenic signaling genes, with strong activation of MYC, a regulator of tumor growth and metabolic reprogramming [1,46]. A second population showed transcriptional signatures of hypoxia-driven metabolic adaptation, marked by increased HIF1A expression, indicating adaptation to oxygen-limited tumor environments [49]. A third subpopulation displayed invasive and angiogenic features, characterized by upregulation of VEGFA, consistent with enhanced vascularization and metastatic potential [6].

Importantly, these malignant states formed a continuous transcriptional spectrum, suggesting dynamic transitions rather than fixed sub-clonal identities. This supports the concept that tumor progression is driven by cellular plasticity and adaptive state switching, influenced by both intrinsic regulatory programs and external microenvironmental signals [49].

These findings extend the cellular heterogeneity described in Section 3.1 by demonstrating that malignant cells undergo functional reprogramming linked to tumor–microenvironment interactions, reinforcing the role of dynamic cellular plasticity in tumor progression and therapeutic resistance [7,47].

### 3.3 Differential Expression and Functional Pathway Enrichment

Differential gene expression analysis identified genes significantly upregulated in tumor-associated cell populations relative to reference groups, revealing key molecular pathways underlying tumor progression and microenvironmental interactions. Functional enrichment analysis using Gene Ontology (GO) and Kyoto Encyclopedia of Genes and Genomes (KEGG) highlighted coordinated activation of immune, inflammatory, and angiogenic signaling pathways [50, 51].

Among the most enriched pathways, immune checkpoint signaling was prominent, indicating active regulation of immune tolerance within the tumor microenvironment. These pathways are known to suppress cytotoxic T-cell activity and facilitate tumor immune evasion [52, 4]. The enrichment of checkpoint-associated genes supports the presence of an immunosuppressive environment consistent with the tumor–immune regulatory interaction identified in Section 3.1.

In parallel, inflammatory cytokine signaling pathways were significantly enriched, reflecting active communication between malignant and immune cells. Cytokine-mediated signaling plays a central role in immune cell recruitment, polarization, and tumor-associated inflammation [2,3]. Notably, the enrichment of cytokine pathways aligns with the observed IL6-associated signaling axis, reinforcing the role of cytokine-driven networks in coordinating tumor–microenvironment interactions [44].

Additionally, angiogenesis-related pathways were enriched across malignant and endothelial populations, characterized by genes involved in vascular development and endothelial activation. Angiogenesis supports tumor growth by providing oxygen and nutrient supply while facilitating metastatic dissemination [1,6].

Collectively, these findings demonstrate that tumor progression is driven by integrated signaling networks linking immune suppression, inflammation, and vascular remodeling. The convergence of these pathways suggests that tumor–microenvironment interactions are governed by coordinated regulatory programs rather than independent processes, providing mechanistic insight into cancer progression and immune evasion [2].

### 3.4 Pseudotime Dynamics of Tumor Progression

Trajectory analysis revealed continuous transcriptional gradients reflecting progressive phenotypic transitions within malignant cell populations. Pseudotime ordering placed tumor cells along a developmental continuum, indicating that tumor progression occurs through dynamic state transitions rather than discrete cellular states [37,38]. Early pseudotime states were characterized by proliferative epithelial-like profiles, whereas later stages exhibited transcriptional signatures associated with invasion and metastatic potential, consistent with prior single-cell studies of tumor evolution [10,11]. Progression along the pseudotime axis was marked by gradual activation of genes linked to cell migration, extracellular matrix remodeling, and angiogenic signaling, indicating a shift toward invasive phenotypes [1,2]. Increased expression of angiogenesis-related genes and stromal regulators in late-stage cells suggests coordinated activation of vascular and microenvironmental remodeling processes during tumor progression [5,6].

Importantly, the trajectory revealed transcriptional signatures consistent with epithelial– mesenchymal transition (EMT), a key process driving tumor invasion and metastasis [53, 54]. EMT- associated genes involved in cytoskeletal reorganization and cellular motility increased progressively along pseudotime, indicating the acquisition of mesenchymal-like phenotypes during tumor evolution.

These findings demonstrate that tumor progression is governed by continuous transcriptional reprogramming and cellular plasticity, linking proliferative, metabolic, and invasive states identified in Section 3.2. The alignment of EMT and angiogenic signaling with late pseudotime stages further supports the role of coordinated regulatory programs in driving tumor progression and microenvironmental adaptation [7].

### 3.5 Regulatory Network Control of Tumor Progression

Gene regulatory network analysis identified several transcription factors acting as central regulatory hubs within tumor-associated gene expression programs. Network topology analysis revealed that transcription factors including STAT3, NF-κB, MYC, and HIF-1α occupied highly connected nodes within the regulatory network, suggesting that these regulators orchestrate key signaling pathways controlling tumor progression and microenvironment interactions. These transcription factors have been widely implicated in cancer development and are known to regulate multiple downstream pathways involved in proliferation, inflammation, and metabolic adaptation [1,46].

Among these regulatory hubs, STAT3 emerged as a prominent driver of tumor-associated signaling networks. STAT3 regulates genes involved in immune suppression, tumor survival, and inflammatory signaling, and persistent STAT3 activation has been observed in many tumor types [55,56]. Similarly, the transcription factor NF-κB plays a critical role in linking inflammatory signaling with tumor progression by regulating cytokine production and immune responses within the tumor microenvironment [57]. Activation of NF-κB signaling has been associated with chronic inflammation and tumor-promoting immune responses in multiple cancers [57].

MYC was also identified as a key regulatory hub controlling transcriptional programs associated with tumor cell proliferation and metabolic reprogramming [1,46]. MYC regulates genes involved in cell cycle progression, ribosome biogenesis, and metabolic pathways that support rapid tumor growth. Additionally, HIF-1α emerged as a critical regulator of hypoxia-responsive pathways that promote angiogenesis and metabolic adaptation in oxygen-deprived tumor environments [49]. These transcription factors collectively form interconnected regulatory circuits that coordinate tumor growth, immune modulation, and microenvironmental adaptation.

To quantify the structural organization of the inferred regulatory network, network topology was evaluated using the clustering coefficient defined as:

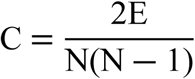

where Crepresents the clustering coefficient, Erepresents the number of edges in the network, and N represents the total number of nodes. High clustering coefficient values indicate dense regulatory interactions among transcription factors and their target genes, reflecting coordinated regulatory control of tumor-associated pathways [58]. The integrated regulatory architecture and hub gene connectivity patterns are illustrated in Figure 6A–C, while pathway integration and regulatory module clustering are shown in Figure 6D–F. Together, these findings highlight the central role of transcriptional regulatory networks in controlling tumor progression and shaping tumor microenvironment dynamics.

**Figure 6.** Integrated Regulatory Landscape of Tumor Progression. (A) Multi-layer network integrating gene regulatory interactions, ligand–receptor signaling, and pseudotime trajectory. (B) Central transcription factor hubs (STAT3, NF-κB, MYC, HIF-1α) and their downstream programs. (C) Pseudotime trajectory of malignant cells colored by transcriptional state (proliferative → invasive). (D) Schematic model summarizing coordinated regulatory modules driving TME remodeling and tumor progression.

**Figure 6A.**
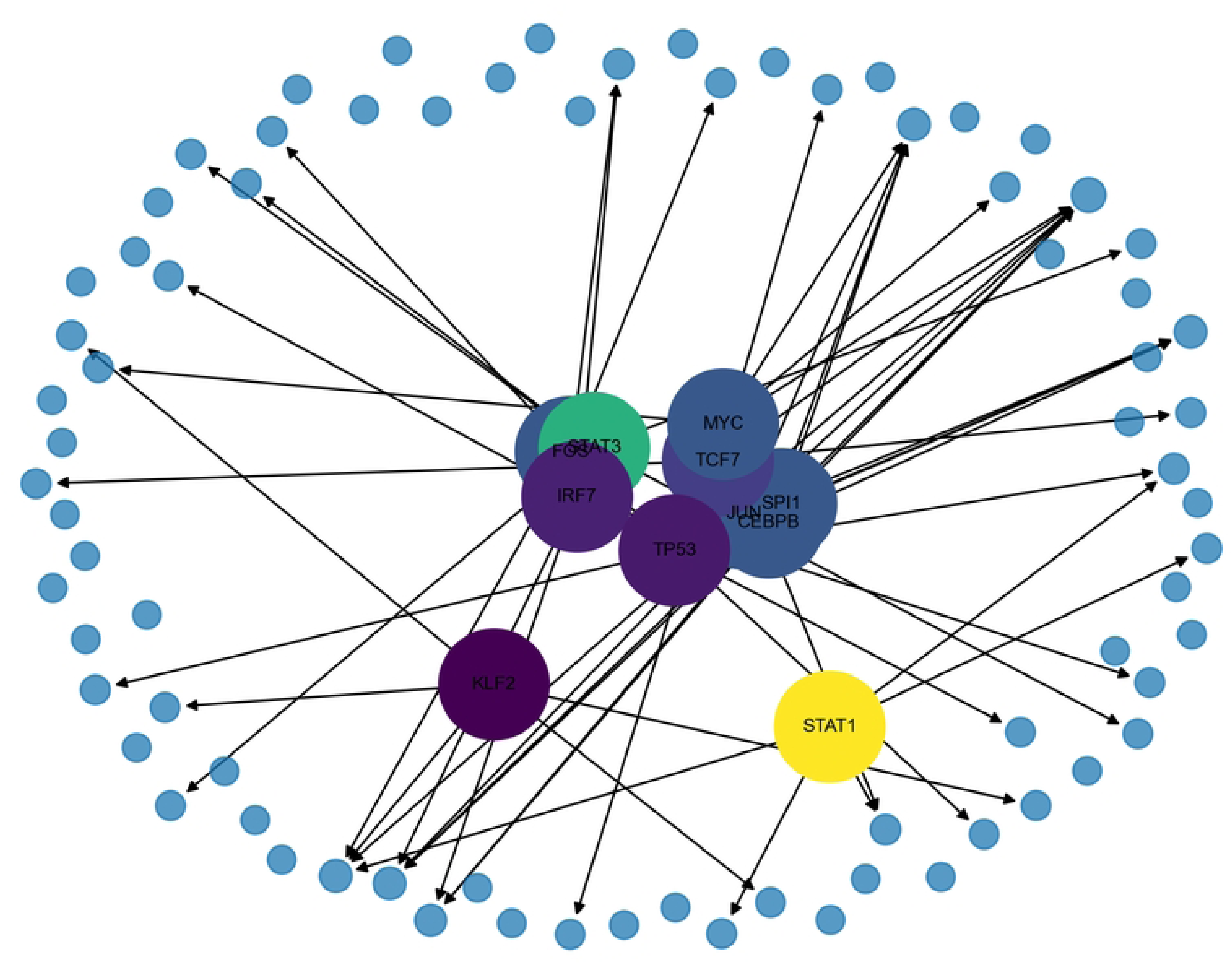
Integrated TF-Target Regulatory Network. This panel illustrates the integrated gene regulatory network (GRN) comprising transcription factors (TFs) and their downstream target genes. Nodes represent TFs and target genes, while directed edges indicate regulatory interactions inferred from co-expression analysis and transcription factor binding motif enrichment. Node size reflects regulatory importance, and node color represents cluster-specific activity levels, linking transcriptional regulation to distinct tumor cell states. This integrated framework combines GRN inference with functional activity profiling, providing a comprehensive view of transcriptional control mechanisms driving tumor progression.

**Figure 6B.**
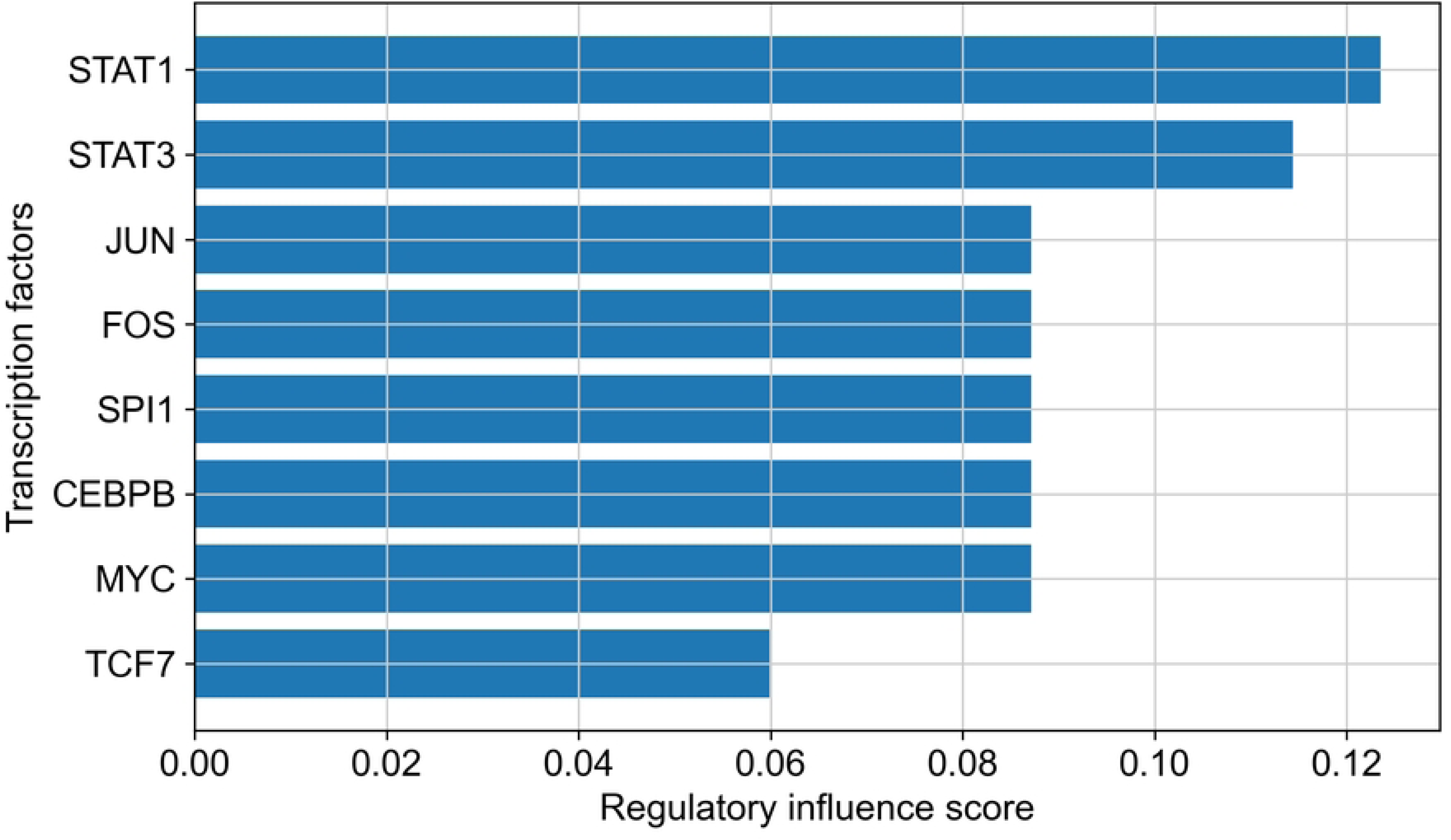
Transcription Factor Influence Ranking. This panel presents the ranking of transcription factors based on their regulatory influence within the network. Influence scores are derived from a combination of centrality metrics, connectivity, and activity levels across cell clusters. Highly ranked TFs, such as MYC, TP53, STAT3, NF-kB, and HIF1A, emerge as master regulators orchestrating key transcriptional programs associated with proliferation, stress response, and immune modulation. This ranking highlights critical drivers of tumor progression and potential targets for therapeutic intervention.

**Figure 6C.**
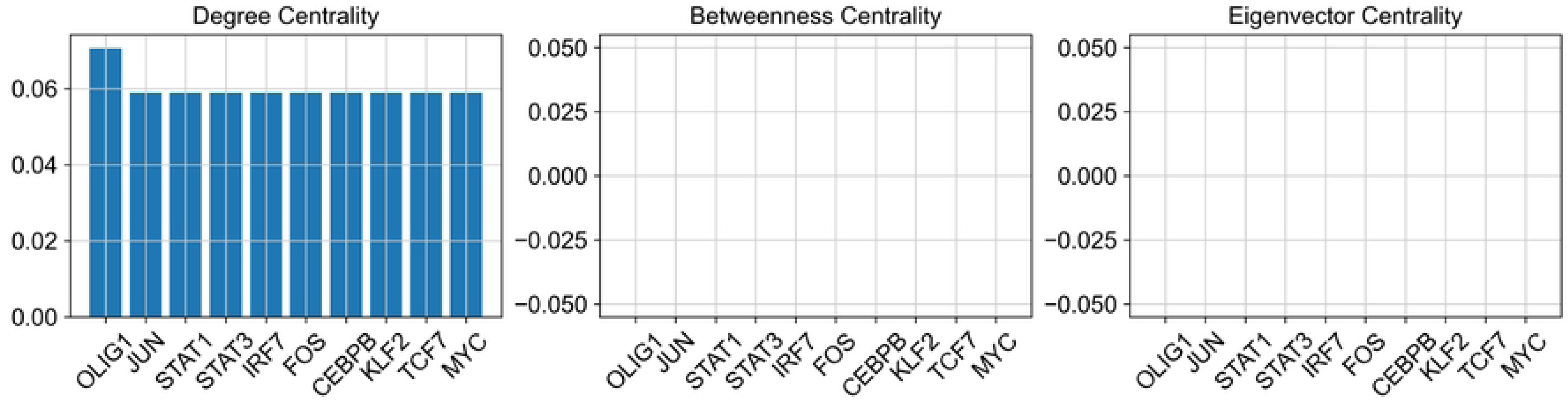
Hub Gene Centrality Analysis. This panel identifies hub genes within the regulatory network using centrality measures, including degree centrality, betweenness centrality, and eigenvector centrality. Hub genes such as TP53, MYC, JUN, and STAT3 occupy central positions in the network and act as key control points for information flow and regulatory coordination. Their prominence underscores their essential role in maintaining network structure and regulating complex transcriptional processes associated with tumor development and adaptation.

**Figure 6D.**
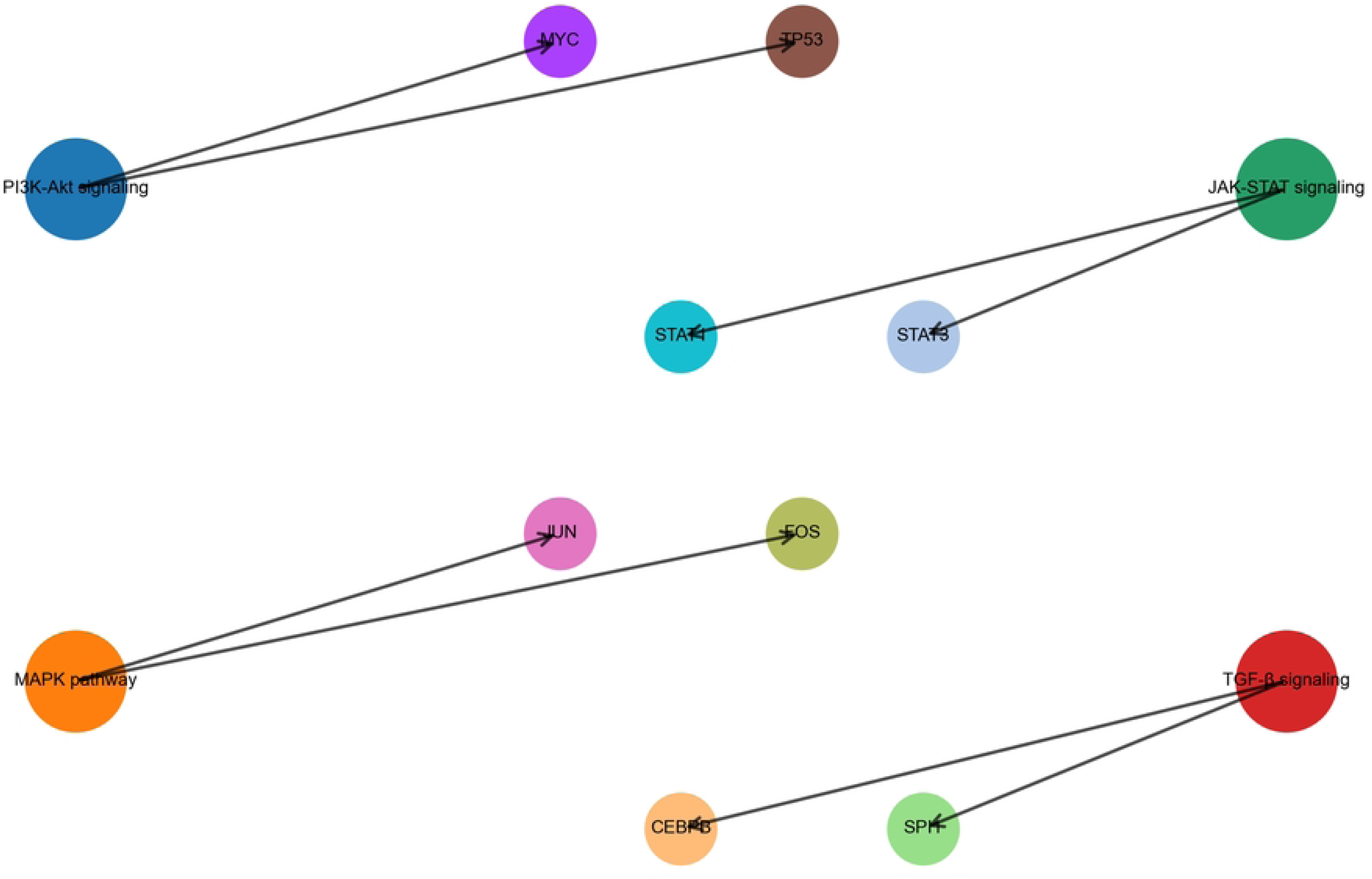
Integrated Signaling Pathway Interaction Map. This panel depicts the cross-talk between transcriptional regulatory modules and major signaling pathways, integrating gene regulatory networks with pathway-level interactions. Key pathways include PI3K-Akt signaling, MAPK signaling, JAK-STAT signaling, and TGF-P signaling. Connections between TFs and pathways illustrate how transcriptional regulation interfaces with signaling cascades to coordinate cellular responses. This multi-layer integration highlights the complexity of regulatory control and the interplay between intracellular signaling and gene expression during tumor progression.

**Figure 6E.**
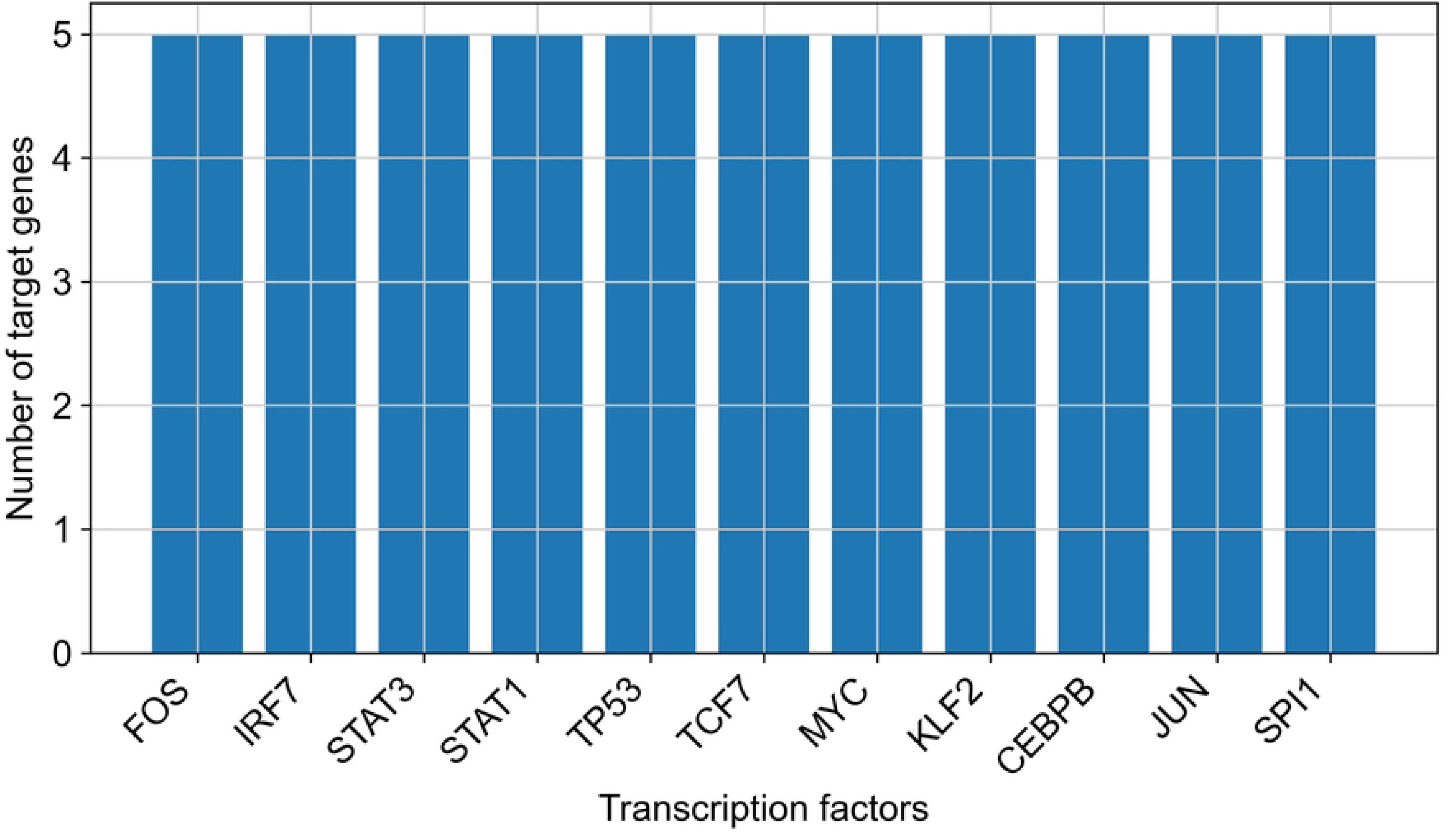
Regulatory Module Clustering. This panel shows the clustering of regulatory modules composed of co-regulated transcription factors and target genes. Modules are grouped based on shared regulatory patterns and functional similarity, with representative modules corresponding to biological processes such as cell cycle regulation, immune response, and hypoxia adaptation. These modules reflect coordinated transcriptional programs that govern different aspects of tumor progression, enabling a modular interpretation of complex regulatory networks.

**Figure 6F.**
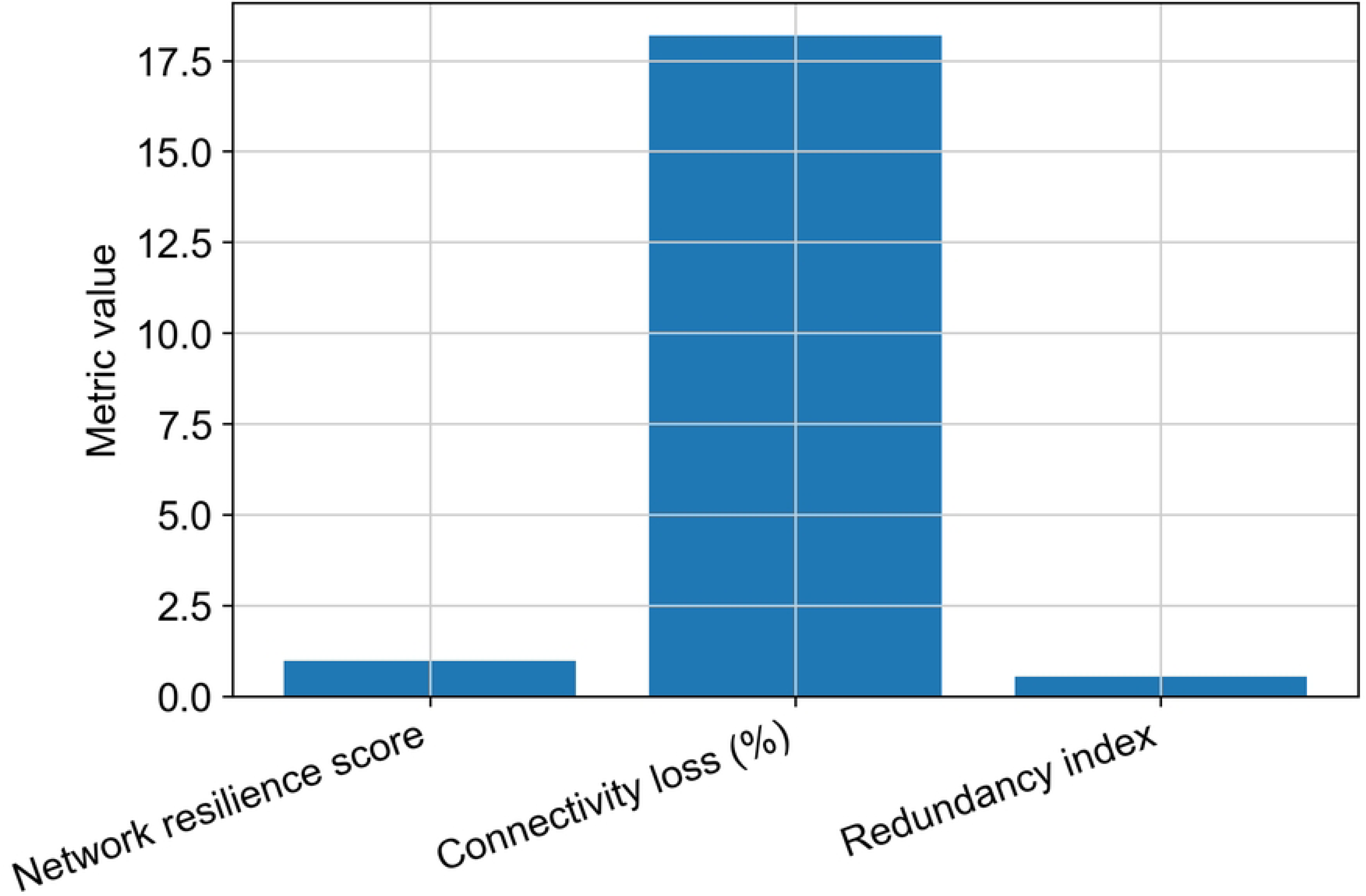
Network Robustness Analysis. This panel evaluates the robustness of the regulatory network under perturbations such as node or edge removal. Metrics including network resilience score, connectivity loss percentage, and redundancy index are used to assess stability and sensitivity. Robust networks maintain functional integrity despite disruptions, indicating the presence of redundant pathways and compensatory mechanisms. This analysis provides insight into the resilience of tumor regulatory systems and their ability to adapt to environmental and therapeutic pressures.

**Figure 6G.**
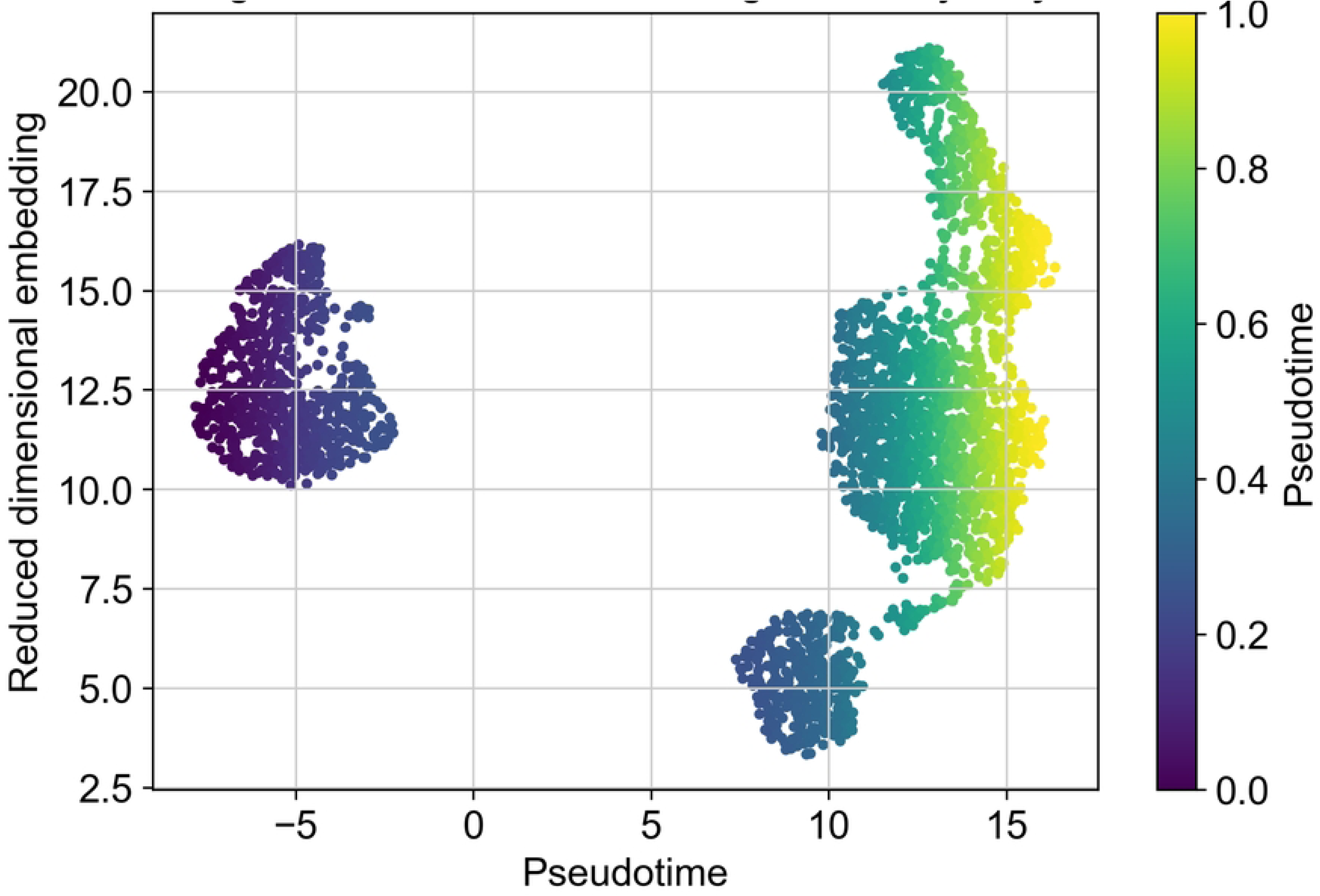
Pseudotime Tumor Progression Trajectory. This panel presents a pseudotime trajectory that orders cells along a continuum representing tumor progression. Cells are arranged based on transcriptional similarity, revealing dynamic changes in gene expression across different stages. The trajectory captures transitions from early to advanced tumor states, highlighting temporal progression and cellular differentiation within the tumor microenvironment. This analysis provides a dynamic perspective on tumor evolution and regulatory changes over time.

**Figure 6H.**
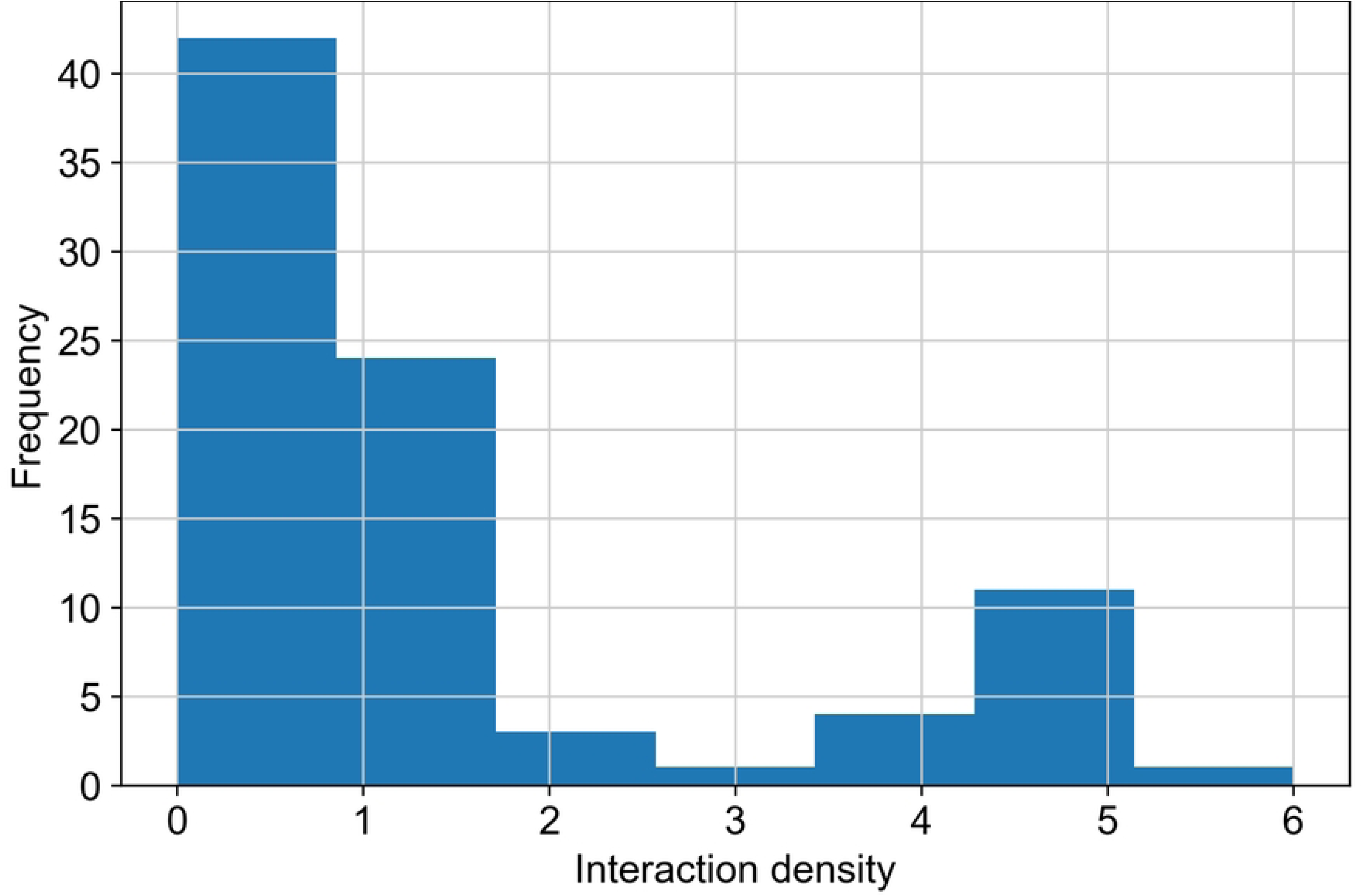
Gene Interaction Network Density. This panel illustrates the distribution of network density, representing the degree of connectivity among genes within the regulatory network. The histogram shows the frequency of interaction densities, with higher density regions indicating tightly connected gene modules. Such dense clusters reflect coordinated regulatory activity and functional specialization, emphasizing the complexity and interdependence of transcriptional networks in tumor biology.

**Figure 6I.**
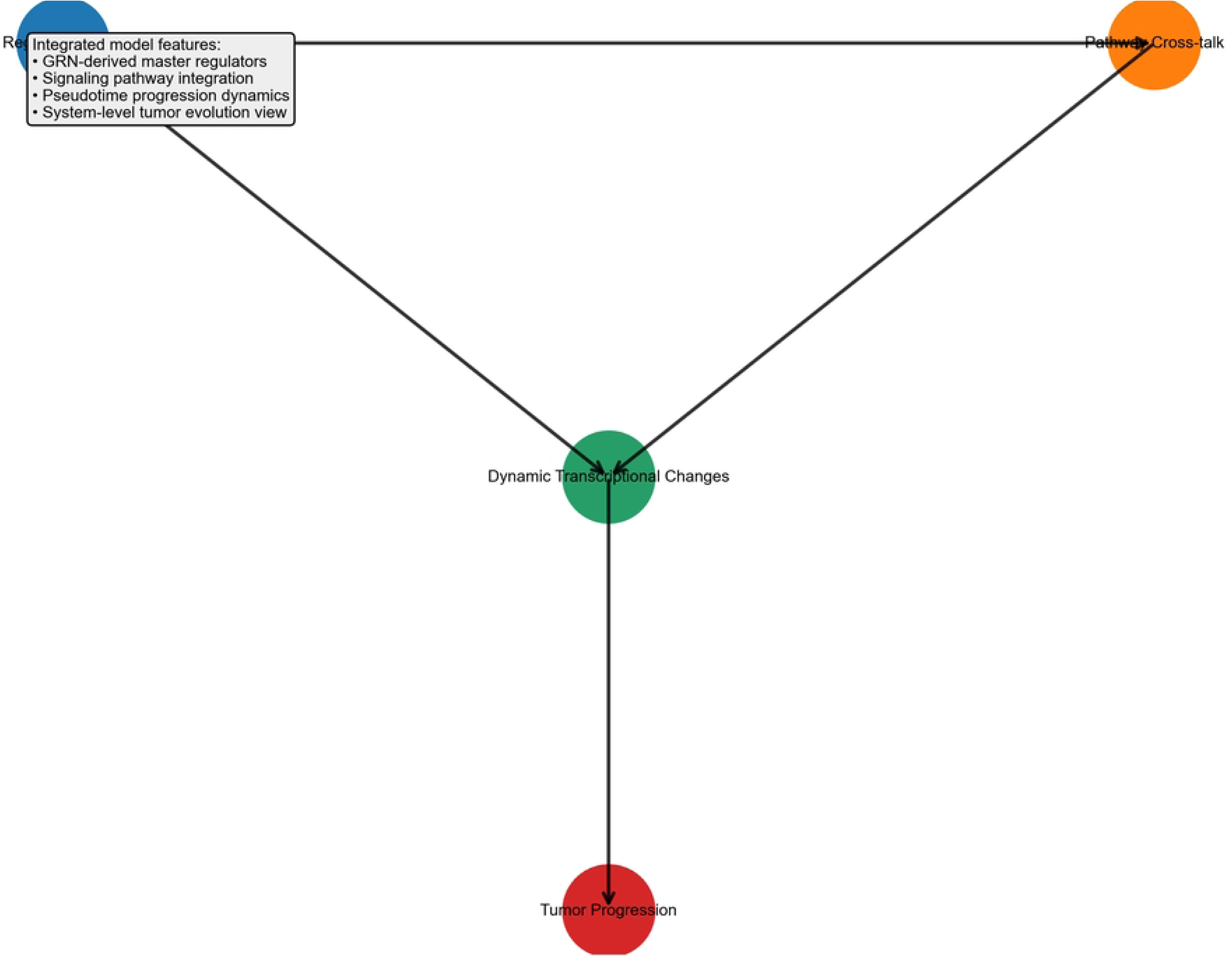
Integrated Tumor Progression Model. This panel presents a comprehensive model integrating gene regulatory networks, signaling pathways, and pseudotime dynamics into a unified framework of tumor progression. The model highlights key regulatory hubs driving progression, pathway cross-talk coordinating cellular responses, and dynamic transcriptional changes across tumor states. By combining multiple layers of analysis, this panel provides a systems-level understanding of tumor evolution, offering insights into the mechanisms underlying tumor growth, adaptation, and therapeutic resistance.

## 4. DISCUSSION

### 4.1 Biological Implications of Tumor Heterogeneity

The results of this study highlight the profound cellular heterogeneity that characterizes the tumor microenvironment and demonstrate how diverse malignant and non-malignant cell populations contribute to cancer progression. Intratumoral heterogeneity is a defining feature of tumor evolution, arising from genetic mutations, transcriptional variability, and selective pressures imposed by the tumor microenvironment [7,8]. Our findings support the idea that tumor progression is driven by transcriptional plasticity rather than fixed clonal states, consistent with observations from single-cell studies in glioblastoma and melanoma. In this context, tumor cells exhibit dynamic transitions between functional states, including proliferative, metabolic, and invasive phenotypes, reflecting adaptive responses to microenvironmental pressures and regulatory signaling networks [10,11]. These findings further suggest that tumor progression may also involve coordinated transcriptional regulatory circuits linking malignant cells and immunoregulatory macrophage populations, highlighting potential targets for therapeutic disruption of tumor microenvironment signaling networks [2–4,44].

The presence of transcriptionally distinct malignant subclones has important implications for therapeutic resistance. Targeted therapies often eliminate dominant tumor populations while leaving resistant subclones unaffected, enabling these populations to expand and drive disease recurrence [7,8]. The identification of MYC-driven proliferative states and HIF-1α–associated metabolic adaptation observed in our analysis suggests that tumor cells can activate alternative regulatory programs to maintain survival under environmental stress conditions [46,49]. Furthermore, the progressive activation of metastasis-associated transcriptional programs along the pseudotime trajectory (Figure 6G) indicates that tumor cells gradually acquire invasive capabilities during tumor evolution. These findings reinforce the importance of resolving cellular heterogeneity at single-cell resolution in order to understand tumor progression and develop more effective therapeutic strategies targeting multiple tumor subpopulations simultaneously [1,2].

### 4.2 Regulatory Networks Driving Tumor Progression

The regulatory network analysis conducted in this study revealed several transcription factors functioning as central regulatory hubs controlling tumor-associated transcriptional programs. Among these regulators, STAT3, NF-κB, MYC, and HIF-1α exhibited high connectivity within the inferred regulatory network (Figure 6A–C), indicating that they play major roles in coordinating gene expression patterns associated with tumor progression. These transcription factors have been widely implicated in cancer biology and regulate diverse processes including cell proliferation, immune modulation, metabolic adaptation, and angiogenesis [1,46].

STAT3 signaling has been shown to promote tumor survival and immune suppression by regulating inflammatory cytokine production and inhibiting anti-tumor immune responses [55]. Similarly, NF- κB functions as a central mediator of inflammation-driven tumor progression by linking inflammatory signaling with oncogenic transcriptional programs [57]. The identification of MYC as a regulatory hub further highlights its role in coordinating metabolic reprogramming and cell cycle progression in rapidly proliferating tumor cells [46]. Additionally, HIF-1α regulates hypoxia-responsive pathways that enable tumor cells to survive in oxygen-deprived microenvironments and promotes angiogenesis through activation of VEGF signaling [6,49].

Together, these interconnected regulatory modules form complex transcriptional circuits that govern tumor growth and adaptation to microenvironmental stress. The regulatory architecture illustrated in Figure 6D–F suggests that coordinated transcription factor activity plays a central role in orchestrating tumor progression and shaping tumor microenvironment dynamics.

### 4.3 Tumor Microenvironment Crosstalk

Our cell–cell communication analysis revealed extensive signaling interactions between malignant cells and surrounding immune and stromal populations within the tumor microenvironment. These interactions are critical determinants of tumor progression because they regulate immune suppression, angiogenesis, and extracellular matrix remodelling [2,3]. The ligand–receptor interaction networks identified in this study (Figure 4) demonstrate how tumor cells exploit signaling pathways to manipulate surrounding cellular populations.

Tumor-associated macrophages and regulatory T cells were found to contribute to immune suppression by producing cytokines and immune checkpoint molecules that inhibit cytotoxic T-cell responses [4, 47]. At the same time, cancer-associated fibroblasts promote stromal activation by remodeling the extracellular matrix and secreting growth factors that facilitate tumor invasion and metastasis [5]. Endothelial cells also play a crucial role in tumor progression by forming abnormal vascular networks that support tumor growth and provide pathways for metastatic dissemination [6]. These findings highlight how tumor–microenvironment interactions actively drive immune suppression, stromal remodeling, and tumor progression.

### 4.4 Study Limitations and Future Directions

While this integrative analysis provides a clearer view of tumor microenvironment (TME) heterogeneity, several technical and biological constraints remain. A primary limitation is the exclusive reliance on transcriptomic data from public repositories. As a result, we were unable to measure protein-level expression or post-translational modifications. Future integration of proteomic or phosphoproteomic data would be essential to confirm that the transcriptional programs identified here translate into functional protein activity.

Furthermore, although our cross-cancer analysis focused on non-small cell lung cancer and breast cancer, the findings may not yet be generalizable to all solid tumors. Validating the conserved nature of the STAT3/NF-κB–macrophage axis will require the use of even larger multi-cancer atlases and more diverse patient cohorts.

Another key consideration is that these findings are derived from dissociated single-cell suspensions, which inherently lack spatial architecture. Because the physical proximity of cells within the TME is vital for the ligand–receptor interactions we inferred [59], the next step will involve spatial transcriptomics to map these communication networks within intact tissue.

Lastly, it is important to emphasize that these results are primarily correlative. Establishing causality for the identified transcription factors and signaling pairs will require functional validation in co- culture systems or patient-derived organoids. Moving forward, combining these scRNA-seq insights with spatial and epigenomic data will offer the systems-level resolution needed to support the translation of these targets toward clinical application in precision oncology.

## 5. CONCLUSION

This study demonstrates the power of integrative single-cell RNA sequencing to dissect the cellular heterogeneity and regulatory architecture of the tumor microenvironment (TME) across multiple cancer types. By combining clustering, pseudotime trajectory inference, gene regulatory network modeling, and ligand–receptor interaction analysis, we resolved distinct populations of malignant epithelial cells, immune subsets, cancer-associated fibroblasts, and endothelial cells, and uncovered pronounced intratumoral transcriptional heterogeneity.

Our analyses identified key transcriptional hubs, including STAT3, NF-κB, MYC, and HIF-1α, that coordinate proliferation, metabolic adaptation, inflammation, and immune evasion. We also uncovered a cytokine-driven immunoregulatory axis linking malignant cells and tumor-associated macrophages via IL6–IL6R and CCL2–CCR2 signaling. These findings highlight how coordinated transcriptional programs and intercellular communication networks shape tumor progression and microenvironmental remodeling.

Collectively, this work provides a systems-level framework for understanding TME organization and underscores the value of multi-dataset integration in revealing conserved regulatory mechanisms. While the results are correlative and require functional validation, they generate actionable hypotheses for disrupting tumor–microenvironment crosstalk. Future integration with spatial transcriptomics, multi-omics approaches, and experimental models will further advance these insights toward precision oncology strategies that target both malignant cells and their supportive ecosystem.

## DECLARATIONS

### Ethics approval and consent to participate

Not applicable.

### Consent for publication

Not applicable.

### Availability of data and materials

The datasets analyzed in this study are publicly available from the Gene Expression Omnibus (GEO), European Nucleotide Archive (ENA), and Single Cell Portal repositories. Accession numbers are provided in Table 1.

**Table 1:** Table 1. Summary of scRNA-seq datasets used in this study. Includes dataset source, cancer type, sample size, and sequencing platform.

| Study | Cancer Type | Dataset Description | Platform | Approx. Cells | Key Features | Source | Accession |
| --- | --- | --- | --- | --- | --- | --- | --- |
| [21] | Non-Small Cell Lung Cancer (NSCLC) | Tumor microenvironment remodeling after neoadjuvant immunotherapy | 10x Genomics Chromium | ~30,000 | Immune reprogramming, TME remodeling, therapy response | GEO | **GSE131907** |
| [22] | Breast Cancer | Single-cell atlas of normal, preneoplastic, and tumorigenic breast tissue | 10x Genomics Chromium | ~50,000 | Tumor progression states, epithelial heterogeneity | GEO | **GSE161529** |
| [23] | Breast Cancer | Processed scRNA-seq dataset and downstream analysis objects from breast cancer atlas | 10x Genomics Chromium | ~40,000 | Data validation, reproducible pipeline, annotation refinement | GEO | **GSE176078** |
| [24] | Breast Cancer (HR+ / HER2-) | Tumor–stromal interaction analysis focusing on CAF–tumor cooperation | 10x Genomics Chromium | ~25,000 | CAF interaction, migration, tumor-stroma signaling | GEO | **GSE114725** |
| [25] | Multi-cancer / Immune-focused | Multi-omics integration for neutrophil heterogeneity using ML | Multi-platform (scRNA + omics) | ~20,000 | Immune heterogeneity, ML integration, neutrophil subtypes | GEO<br>ENA | **GSE157344 / PRJNA720124** |

**Table 2:**
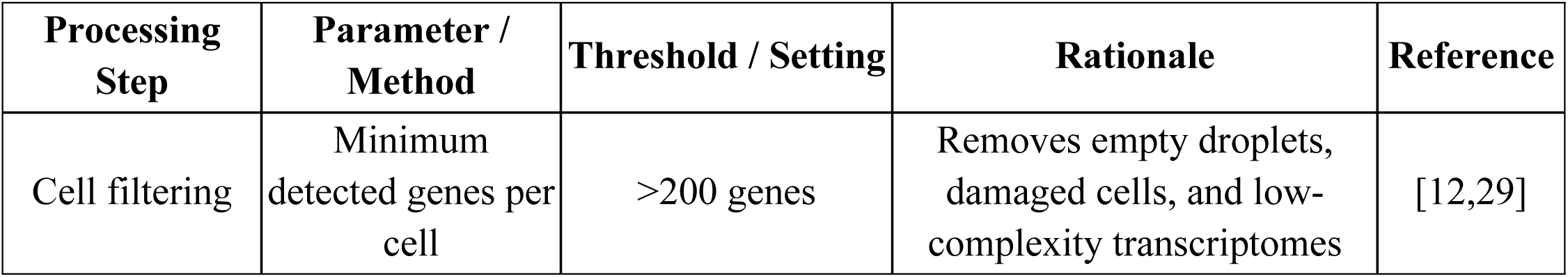
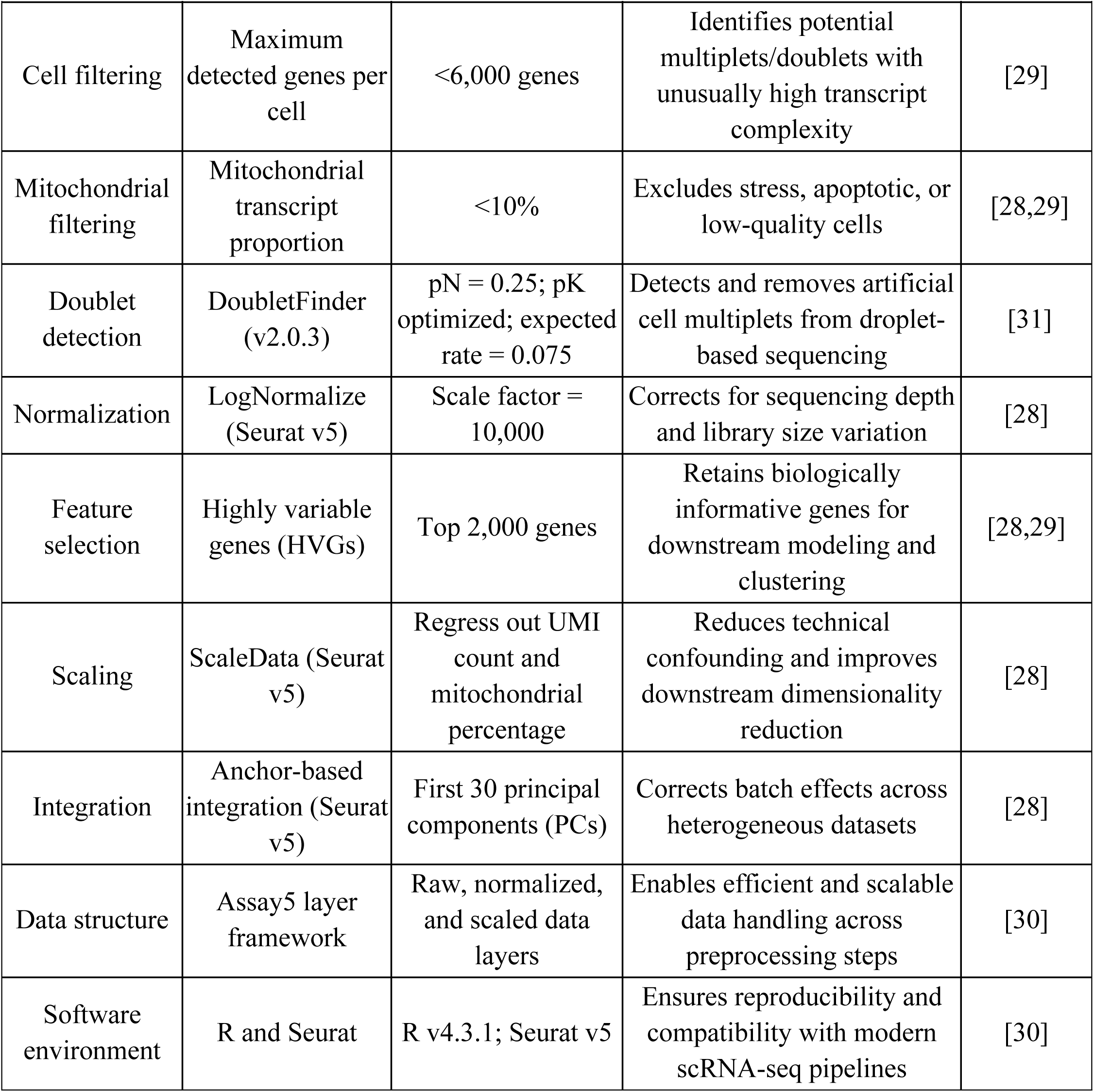
Quality Control and Preprocessing Parameters for Single-Cell Transcriptomic Analysis Lists filtering thresholds and criteria applied during scRNA-seq data preprocessing.

| Processing Step | Parameter / Method | Threshold / Setting | Rationale | Reference |
| --- | --- | --- | --- | --- |
| Cell filtering | Minimum detected genes per cell | >200 genes | Removes empty droplets, damaged cells, and low-complexity transcriptomes | [12,29] |
| Cell filtering | Maximum detected genes per cell | <6,000 genes | Identifies potential multiplets/doublets with unusually high transcript complexity | [29] |
| Mitochondrial filtering | Mitochondrial transcript proportion | <10% | Excludes stress, apoptotic, or low-quality cells | [28,29] |
| Doublet detection | DoubletFinder (v2.0.3) | pN = 0.25; pK optimized; expected rate = 0.075 | Detects and removes artificial cell multiplets from droplet-based sequencing | [31] |
| Normalization | LogNormalize (Seurat v5) | Scale factor = 10,000 | Corrects for sequencing depth and library size variation | [28] |
| Feature selection | Highly variable genes (HVGs) | Top 2,000 genes | Retains biologically informative genes for downstream modeling and clustering | [28,29] |
| Scaling | ScaleData (Seurat v5) | Regress out UMI count and mitochondrial percentage | Reduces technical confounding and improves downstream dimensionality reduction | [28] |
| Integration | Anchor-based integration (Seurat v5) | First 30 principal components (PCs) | Corrects batch effects across heterogeneous datasets | [28] |
| Data structure | Assay5 layer framework | Raw, normalized, and scaled data layers | Enables efficient and scalable data handling across preprocessing steps | [30] |
| Software environment | R and Seurat | R v4.3.1; Seurat v5 | Ensures reproducibility and compatibility with modern scRNA-seq pipelines | [30] |

**Table 3:**
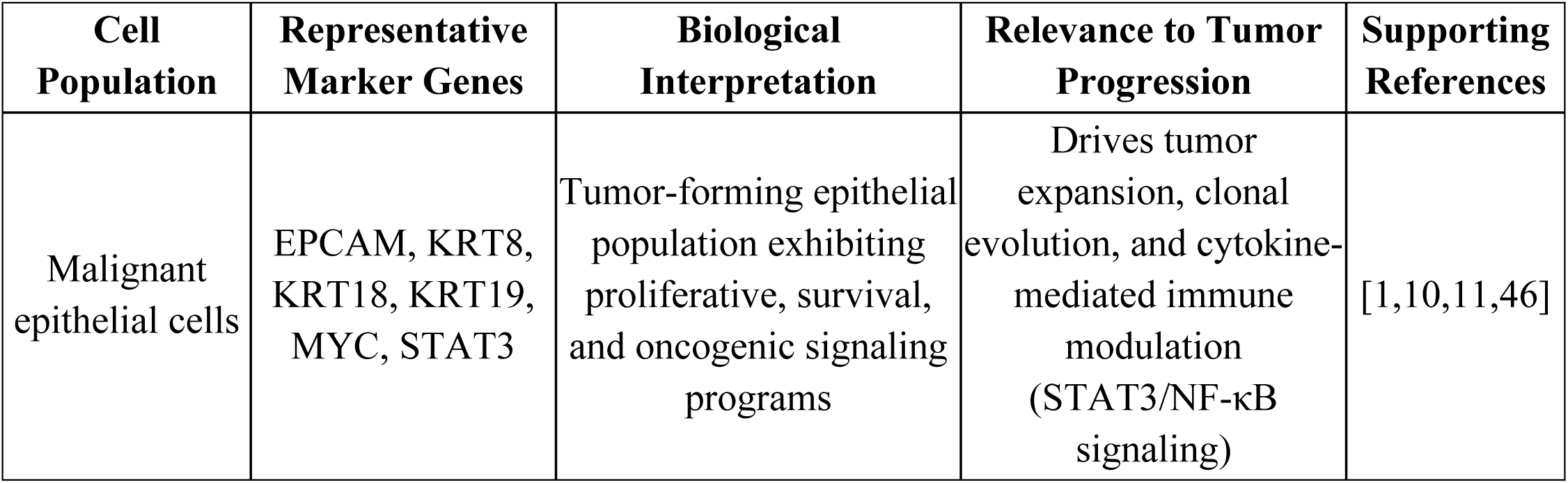
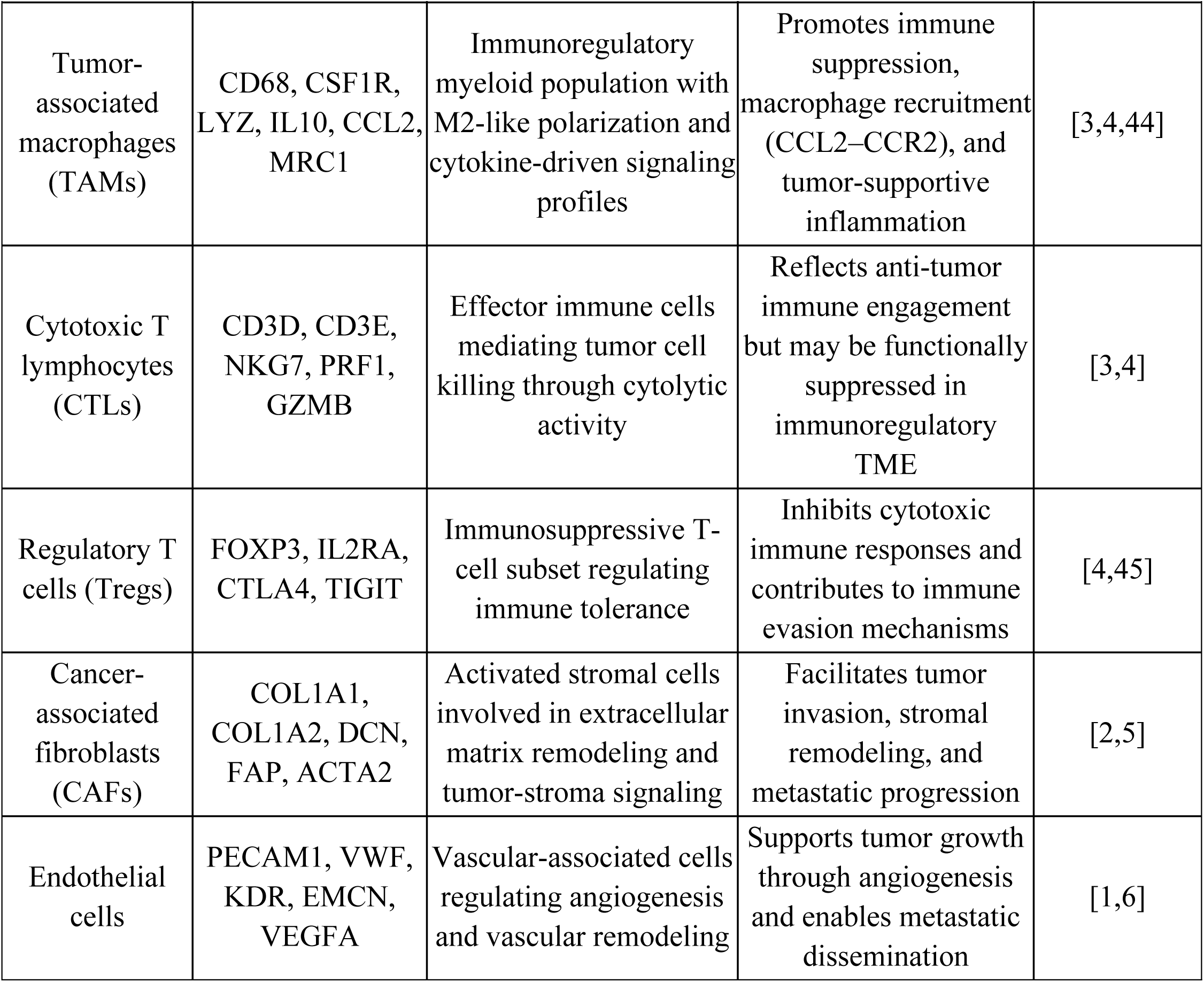
Major Cell Populations in the Tumor Microenvironment with Marker Genes and Functional Roles. Displays key marker genes used to identify major cell populations.

**Table 4:**
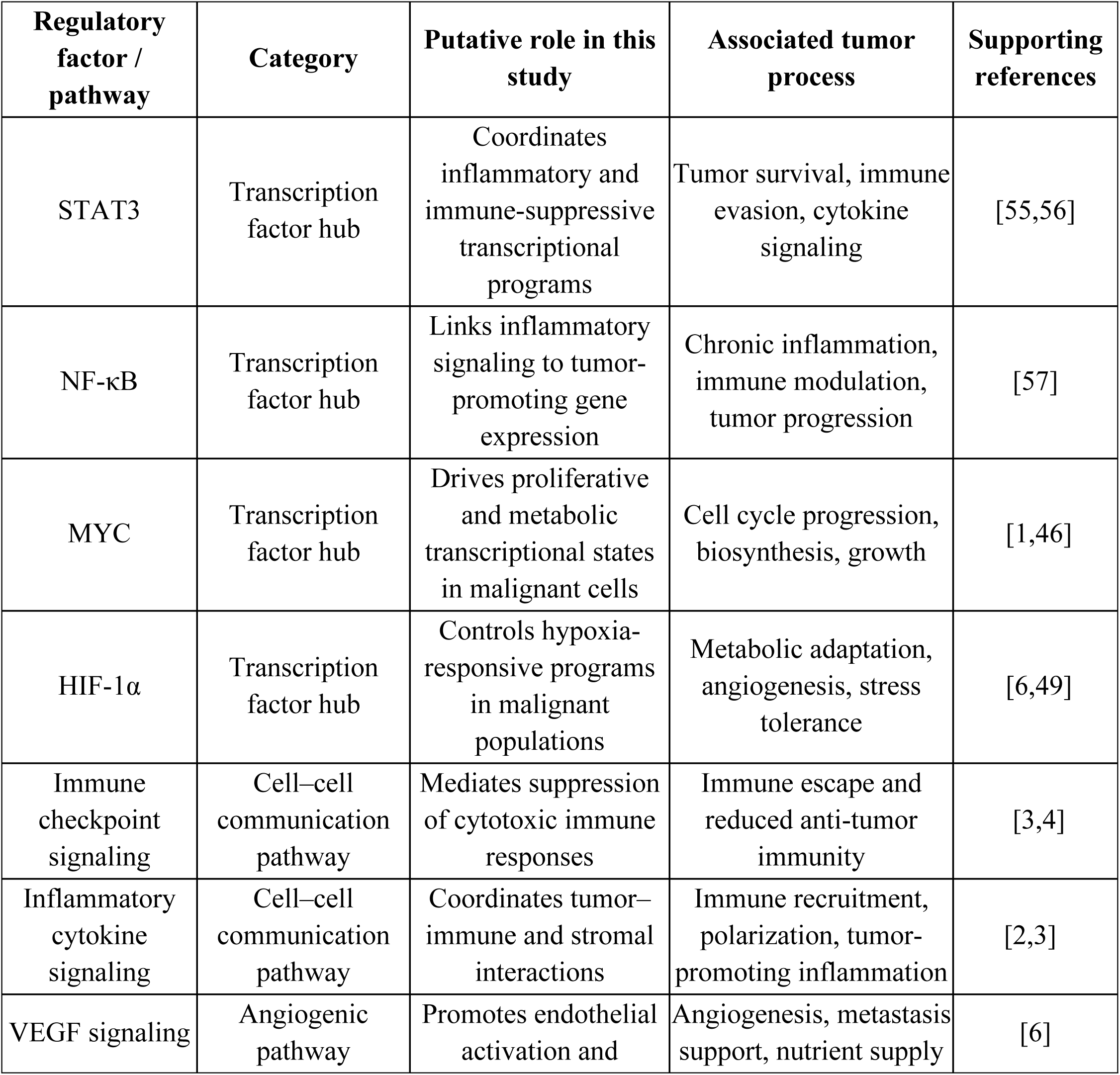
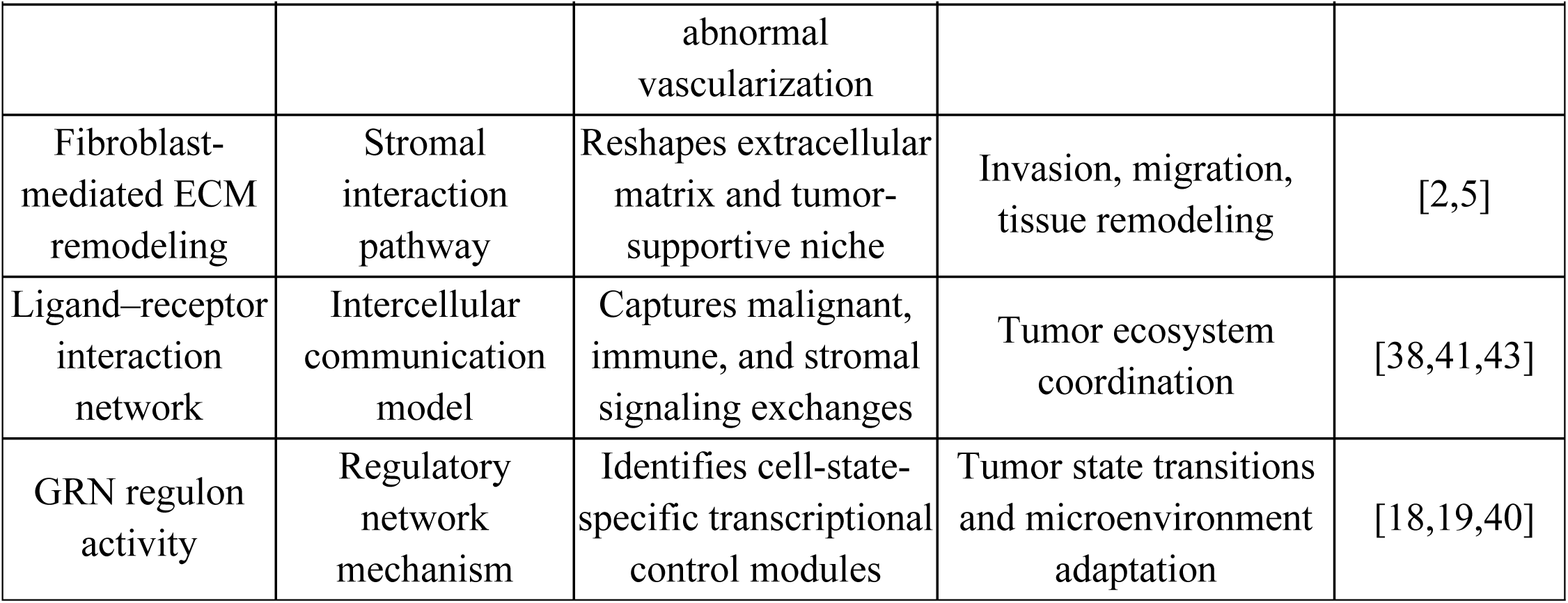
Key regulatory hubs and tumor microenvironment communication pathways identified in the study. Highlights major regulatory hubs and their functional roles in tumor progression.

### Competing interests

The authors declare that they have no competing interests.

## Funding

No specific funding was received for this study.

## Authors’ contributions

MOO conceived the study, performed the bioinformatic analyses, interpreted the results, drafted and revised the manuscript. CEE contributed to study design, data interpretation, visualization, and critically revised the manuscript. All authors read and approved the final manuscript.

## Notes

### Competing Interest Statement

The authors have declared that no competing interests exist.

